# Transcriptional fluctuations govern the serum dependent cell cycle duration heterogeneities in Mammalian cells

**DOI:** 10.1101/2022.05.06.490879

**Authors:** Vinodhini Govindaraj, Subrot Sarma, Atharva Karulkar, Rahul Purwar, Sandip Kar

**Affiliations:** Department of Chemistry, IIT Bombay, Powai, Mumbai 400076, India; Department of Biosciences and Bioengineering, IIT Bombay, Powai, Mumbai 400076, India

**Keywords:** Cell cycle duration variabilities, live-cell imaging, transcriptional fluctuation mathematical, computational modeling

## Abstract

Mammalian cells exhibit a high degree of intercellular variability in cell cycle period and phase durations. However, the factors orchestrating the cell cycle duration heterogeneities remain unclear. Herein, by combining cell cycle network-based mathematical models with live single-cell imaging studies under varied serum conditions, we demonstrate that fluctuating transcription rates of cell cycle regulatory genes across cell lineages and during cell cycle progression in mammalian cells majorly govern the robust correlation patterns of cell cycle period and phase durations among sister, cousin, and mother-daughter lineage pairs. However, for the overall cellular population, alteration in serum level modulates the fluctuation and correlation patterns of cell cycle period and phase durations in a correlated manner. These heterogeneities at the population level can be finetuned under limited serum conditions by perturbing the cell cycle network using a p38-signaling inhibitor without affecting the robust lineage level correlations. Overall, our approach identifies transcriptional fluctuations as the key controlling factor for the cell cycle duration heterogeneities, and predicts ways to reduce cell-to-cell variabilities by perturbing the cell cycle network regulations.

**Significance statement:** In malignant tumors, cells display a diverse pattern in cell division time. This cell-to-cell variability in cell cycle duration had been observed even under culture conditions for various mammalian cells. Here we used live-cell imaging studies to monitor FUCCI-HeLa cells and quantified the cell cycle period and time spent in different phases under varied serum conditions. We proposed a set of stochastic cell cycle network-based mathematical models to investigate the live-cell imaging data and unraveled that the transcription rate variation across cell lineages and during cell cycle phases explains every aspect of the cell cycle duration variabilities. Our models identified how different deterministic effects and stochastic fluctuations control these variabilities and predicted ways to alter these cell cycle duration variabilities.

## Introduction

Cell cycle period and phase durations of mammalian cells demonstrate a high degree of heterogeneity under culture conditions (1–9), and within a tumor micro-environment (10, 11). Often, such heterogeneities significantly influence the cell-fate decision-making (12–16). This means that these heterogeneities can be fine-tuned to design better therapeutics if the factors controlling these variabilities are known precisely. However, identifying these controlling factors is highly challenging due to the diverse and complex nature of cell cycle duration heterogeneities. For example, The cell cycle phase durations quantified at the single-cell level in proliferating lymphocytes (8, 17) using FUCCI-reporter (18) showed a highly variable S-G_2_-M time with strongly correlated G_1_ timings in the sibling cell pairs. A highly correlated cell cycle in sibling pairs has been observed in other mammalian cell types as well (19–22) with mothers and daughters having poor correlation in cell cycle timings, while cousins showing a significant correlation for the same (19–22). These observations are quite generic and cell-type independent, however, the precise understanding of the nature and origin of these heterogeneities remains elusive.

To understand such complex inheritance patterns of cell cycle durations in cell lineages, different kinds of mathematical modeling studies were employed. The model based on transition probability by Dowling et al. assumed that cells transit from one cell cycle phase to another randomly, and consequently create no/poor correlation in mother-daughter pairs (17). However, the cell cycle period of siblings are correlated as cells spent an equal proportion of division time in S-G2-M phases (17). A ‘kicked cell cycle model ‘ proposed by Sandler et al. considered that another oscillator like the circadian clock eventually influences cell cycle durations and causes the observed correlation pattern in cell lineages (20). Recent models using bifurcating autoregressive (BAR) approach suggest that the inheritance of more than one deterministic factor account for the observed correlation pattern in cell lineages (21, 22). It was evident that these modeling studies came up with a widely varied qualitative explanation for the inheritance pattern. However, none of these studies considered the well-established cell cycle gene-interaction network which dynamically controls the cell cycle progression and the associated variabilities (23, 24) within the cell lineages and even at the overall cellular population level. It can be envisaged that an appropriate network-based stochastic cell cycle model will provide the opportunity to identify the effect of various kinds of noise sources like, (i) intrinsic noise due to gene expression variabilities (25–27), (ii) extrinsic noises due to cell to cell variability in the transcription rates of cell cycle related genes within different lineages (28–32), epigenetic modifications during the cell cycle (33– 36), random partitioning of molecules during cell division (23, 24), etc., and (iii) even the deterministic factors that ultimately orchestrate the cell cycle duration heterogeneities and the inheritance pattern.

Moreover, in most of the previous single-cell studies (20–22), the heterogeneities in cell cycle duration have been quantified for a healthy growing condition. However, the cell cycle durations are known to get altered due to changes in the growth environment (17, 19), and in malignant tumors, the cells continue to proliferate even under minimal growth condition (7, 37). This suggests that the cell cycle durations and the corresponding heterogeneities associated with it may change according to the serum level. Earlier, it was shown that for a specific growth condition, most of the cell cycle period variation happens due to G_1_-phase duration variability while S-G_2_-M time remains relatively constant (38), while recent studies demonstrate that even S-G_2_-M phase duration can also be highly variable (17). However, the exact nature and to what extent the serum level influences the cell cycle duration heterogeneities, remain unresolved. In this article, we take a system biology approach to unravel how the change in serum level simultaneously modifies the heterogeneities of the cell cycle phase durations and inheritance pattern among the cell lineage pairs and for the overall cellular population under culture conditions. Our proposed network-based stochastic cell cycle model allowed us to address these pertinent questions; (i) Which is the factor that influentially governs the correlation pattern of cell cycle period and phase durations for cell lineage pairs under different serum conditions? (ii) Which noise source (Intrinsic or Extrinsic) majorly controls the variance of the cell cycle period and phase duration distributions? (iii) How do serum levels alter the fluctuation and correlation patterns of cell cycle period and phase durations for the overall cellular population? and (iv) Can we fine-tune these cell cycle period and phase duration heterogeneities for the overall cellular population to attain a therapeutically advantageous situation?

To answer these questions, we quantified the intercellular variability and lineage correlations in cell cycle period and phase durations from single-cell imaging data and investigated the origin of these variabilities and correlations using our network-based stochastic cell cycle model. Live-cell imaging of FUCCI-HeLa cells under different serum conditions depict that the variabilities and correlations observed in cell cycle and phase durations can be modulated in a serum-dependent manner. Our modeling study reconciled the experimental data and demonstrated that cell-to-cell variability in transcription propensities of the cell cycle regulatory genes is one of the major sources of variability in cell cycle period and phase durations. It further revealed that the memory of transcriptional activity in cell lineages leads to the correlation pattern observed in mother-daughter, sibling, and cousin pairs under varied serum conditions. Finally, we showed that it is possible to fine-tune the heterogeneities of cell cycle period and phase durations under a specific serum level without altering the inheritance pattern by just altering specific gene expression via inhibition of the p38-signaling (39, 40) pathway.

## Results

### Alteration in serum level influences the cell cycle period and phase duration heterogeneities

To investigate cell cycle progression at the single-cell level, we produced FUCCI (18) expressing stable HeLa cell-line (FUCCI-HeLa, **Materials and Methods**) to precisely quantify the cell cycle period and phase durations (**Fig. 1a**). We performed live-cell imaging studies with FUCCI-HeLa cells under two different serum concentrations 2% (low, **Movie S1** and **Movie S2, Fig. 1b**) and 10% (high, **Movie S3** and **Movie S4, Fig. 1c**) and measured the cell cycle period (T_CC_), and phase duration (T_G1_ & T_S-G2-M_) distributions by following FUCCI reporter trajectories from several single cells (**Fig. 1d-e**) (details in **Materials and Methods**). To calculate the statistics for T_CC_, T_G1_ & T_S-G2-M_ timings, we only considered those cells which completed their cell cycle during the imaging period. We observed across different replicates (**Table S1**) that cells exhibit a longer G_1_ and cell cycle mean durations for 2% serum (**Fig. 1d, Fig. S1**, and **Table S1**) in comparison to 10% serum condition (**Fig. 1e, Fig. S1**, and **Table S1**), while the mean S-G_2_-M duration shows a slight increase or no perceptible change (**Table S1**) under 10% serum concentration (**Fig.1d-1e**). While T_G1_ variability remains higher than T_S-G2-M_ variability under both the serum conditions, the mean CV (coefficient of variation) of T_G1_ and T_CC_ duration decreases, and T_S-G2-M_ increases at 10% serum compared to 2% serum (**Fig. 1f** and **Table S1**).

**Fig. 1.**
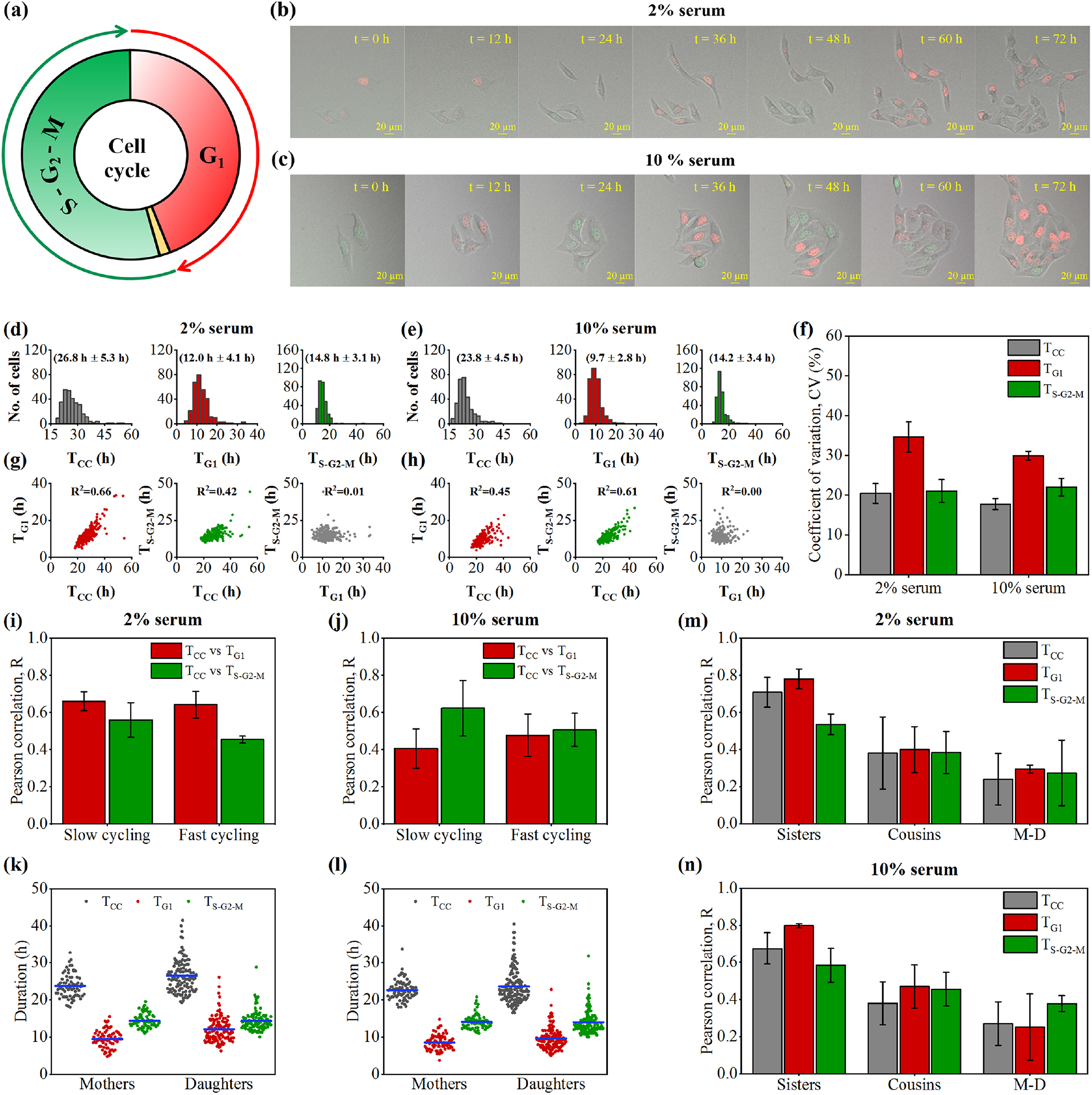
Live cell imaging of FUCCI-HeLa cells quantifies the serum-dependent cell cycle period and phase duration heterogeneities. (**a**) Schematic depiction of FUCCI-reporter expression dynamics during the cell cycle. Live-cell imaging snapshots (at 12 h intervals) of FUCCI-HeLa cells under (**b**) 2% serum, and (**c**) 10% serum conditions. **T_CC_, T_G1_** and **T_S-G2-M_** distributions quantified from single-cell imaging data at (**d**) 2% serum (n=300, replicate 1) and (**e**) 10% serum (n=300, replicate 1) (4 replicates are performed, **Table S1**). (**e**) CV of **T_CC_, T_G1_** and **T_S-G2-M_** distributions were observed across 4 individual replicates (**Table S1**) performed at 2% and 10% serum conditions. Correlations among **T_CC_, T_G1_** and **T_S-G2-M_** were observed from the single-cell imaging at (**g**) 2% serum (n=300, replicate 1) and (**h**) 10% serum (n=300, replicate 1) (4 replicates are performed, **Table S1**). Correlation between **T_CC_, T_G1_** and **T_S-G2-M_** in slow and fast cycling cells at (**i**) 2% serum and (**j**) 10% serum for 4 individual replicates (**Table S1**). **T_CC_, T_G1_** and **T_S-G2-M_** distributions for mother and daughter cells (replicate 1) at (**k**) 2% serum and (**l**) 10% serum (4 replicates are performed, **Table S1**). Observed correlations (Mean ± standard deviation) for **T_CC_, T_G1_** and **T_S-G2-M_** in sisters, cousins and mother-daughter pairs across 4 individual replicates (**Table S1**) at (**m**) 2% serum and (**n**) 10% serum conditions.

Thus, one can presume that the T_G1_ and T_S-G2-M_ are mutually adjusting in a correlated manner during the cell cycle progression. However, the T_G1_ and T_S-G2-M_ showed poor correlation under both serum conditions (**Fig. 1g-1h** and **Fig. S1**). This indicates that the phase durations do not mutually influence each other i.e. a shorter T_G1_ does not cause a longer T_S-G2-M_ or vice versa, which had been previously reported as well (42). Interestingly, our experiments revealed that under a 2% serum condition, T_CC_ correlates to a higher extent with T_G1_ in comparison to T_S-G2-M_ (**Fig. 1g**). This confirms that T_G1_ being highly variable (**Fig. 1d**) indeed greatly influences the T_CC_ at 2% serum condition.

However, higher serum concentration (10%) decreases the T_CC_ vs T_G1_ correlation and increases the T_CC_ vs T_S-G2-M_ correlation (**Fig. 1h**) suggesting that T_S-G2-M_ variability dictates the T_CC_ variability at 10% serum, even though the magnitude of T_G1_ variability is higher than the corresponding T_S-G2-M_ variability (**Fig. 1e**). These observations state that variations in serum condition modulate the T_CC_, T_G1_, and T_S-G2-M_ heterogeneities in a specific manner at the population level.

### Cells within a subpopulation and in cell lineages show distinct cell cycle period and phase duration pattern

Often, a population-level cellular response can be influenced by a subpopulation of cells with distinct behaviour (43, 44). To investigate this aspect, we have divided the cellular population into two subpopulations based on the average T_CC_ (**Fig. S2A**) to understand the contribution of the slow and fast cycling cells in the observed correlation pattern between T_CC_ vs T_G1_ and T_CC_ vs T_S-G2-M_ under varied serum conditions. We find that under low serum (2%) conditions, both fast and slow-cycling cells have a high T_CC_ vs T_G1_ correlation (**Fig. 1i** and **Fig.S2b**). However, at 10% serum condition, slow-cycling cells have an increased T_CC_ vs T_S-G2-M_ correlation than the fast-cycling cells (**Fig. 1j** and **Fig.S2b**). This indicates that slow-cycling cells cause the higher T_CC_ vs T_S-G2-M_ correlation for the overall cellular population at 10% serum. Further, we analyze this continuously dividing the cellular population by following the cells over specific cell lineages which eventually produces subpopulations of mother-daughter, sister-sister, and cousin-cousin pairs. Previous studies had demonstrated that mother-daughter, sister-sister, and cousin-cousin pairs show distinct correlation patterns of cell cycle period and phase durations (17, 19–22). We examined similar subpopulations and observed that on an average mother cell population had a faster cycling time than the daughter population under both 2% and 10% serum levels (**Fig. 1k-1l**) with poor correlation in T_CC_ and phase timings between mother and daughter pairs (**Fig. 1m-1n**). We observed a highly correlated T_CC_ among sister pairs (**Fig. 1m-1n**) with a comparatively higher correlation in T_G1_ than T_S-G2-M_ (17, 19–22). This suggests that the sister cells execute the G_1_-phase in a coordinated manner than the S-G_2_-M phase, where the phase duration could get altered among the sister cells due to various reasons. Cousin pairs showed a moderate level of correlation which was in between the sisters and mother-daughter pairs (**Fig. 1m-1n**). Importantly, serum level does not affect the T_CC_, T_G1_, and T_S-G2-M_ correlations in sisters, cousins, and mother-daughter pairs significantly (**Fig. 1m-1n**). However, it was hard to disentangle the underlying factors which critically govern such kind of fluctuation and correlation patterns under different serum conditions explicitly by only analyzing these experimental observations.

### A stochastic cell cycle model to analyze the serum dependent cell cycle period and phase duration heterogeneities

Over the years, many cell cycle models (45–52) have been developed to address different aspects of mammalian cell-cycle regulations in a context-dependent manner. In a similar spirit, we developed a minimalistic cell cycle network-based (**Fig. 2a**) mathematical model (**Table S2**) to investigate the origin of heterogeneities observed under different serum conditions. The model (**Fig. 2a**) includes three different modules which consist of regulatory interactions controlling different phases of the cell cycle to generate a cycling cell. The first module (**Module-I, Fig. 2a**, and **Fig. S3**) depicts the early G_1_ phase regulation along with the restriction point control mechanism that organizes the decision-making for the cell cycle commitment by sensing the serum level (53, 54). This module contains serum-mediated Myc and CycD activation that initiates the phosphorylation of Rb protein by the Cyclin-dependent kinase (CDK) complex Cdk1-CycD to partially activate the transcription factor E2F1. The second module (**Module-II, Fig. 2a**, and **Fig. S3**) delineates the molecular events controlling the G_1_-S transition and the S-phase activities which primarily depend on the two important Cyclin-dependent kinase regulators, CDK2-CycE and CDK2-CycA. These CDK-cyclin complexes overcome the inhibitions by the Cyclin-dependent kinase inhibitors (CKI ‘s in the form of p21 and p27) and Cdh1 to help the cells cross the restriction point by complete activation of E2F1 which coordinates the G_1_-S transition and active S-phase related genes. The third module (**Module-III, Fig. 2a**, and **Fig. S3**) describes the molecular regulation of the G_2_ and M phases in a nutshell. It considers that the Cdk1-CycB complex in the G_2_-phase overcomes the inhibition by Wee1, and gets activated by Cdc25 due to positive-feedback regulation. It further activates Cdc20 in M-phase which allows the cells to exit from the M-phase by activating Cdh1. Additionally, we have introduced the Cdt1 and Geminin proteins in our model to measure the T_CC_, T_G1_, and T_S-G2-M_ precisely from our model.

**Fig. 2.**
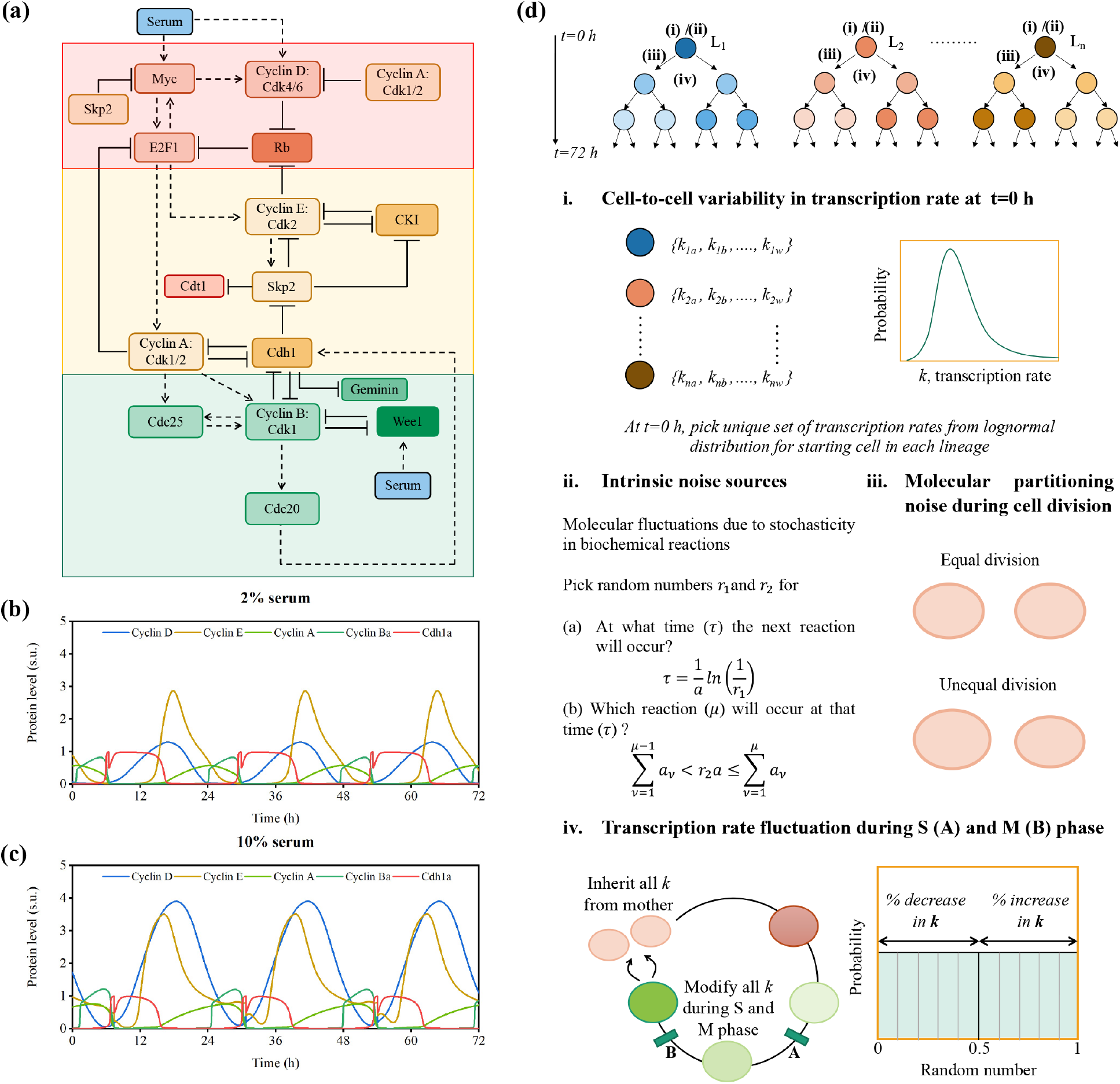
Proposed cell cycle network-based stochastic-mathematical model to understand the experimentally observed serum-dependent cell duration heterogeneities. (**a**) The minimalistic cell cycle network under the influence of serum. (Solid and dotted arrows indicate the direct and indirect activation processes. Hammerhead arrows indicate inhibition processes.) **Module-I** depicts early G0 to G1 regulations. **Module-II** contains the optimal G1-S transition and S-phase regulations. **Module-III** represents the G2-M phase interactions in a simplified manner (Details in **Supplementary Text-1**). The temporal dynamics for Cyclins and active Cdh1 at (**b**) 2%, and (**c**) 10% serum. (**d**) Numerical simulation strategy to simulate different cell lineages by incorporating extrinsic ((**i**) cell-to-cell transcription rate variability, (**ii**) symmetric and asymmetric cell division event, (**iii**) transcription rate variation during cell cycle process due to epigenetic modifications, etc.) and intrinsic ((**ii**) due to molecular fluctuations for low copy numbers of proteins and mRNA ‘s) noise sources experienced by the mammalian cells under culture conditions.

A detailed description of different modules and corresponding molecular events are provided in the **supplementary material**. We developed an ordinary differential equation (ODE) based mathematical model (**Table S3**) for the detailed cell cycle network (**Fig. 2A** and **Fig. S3**) and the kinetic parameters for the model are depicted in **Table S4**. The model reproduced the experimentally observed mean T_CC_, T_G1_, and T_S-G2-M_ (**Fig. 2b** for 2% and **Fig. 2c** for 10% serum) under specific serum doses for FUCCI-HeLa cells. Here, we made sure that the number of molecules of mRNAs and proteins is reasonably higher, as in mammalian cells (55–58) (**Fig.2b-c**), these numbers are considerably higher than budding (59–61) or fission yeast cells (62). Such a deterministic model sets the stage for the stochastic model, where both intrinsic and extrinsic fluctuations can be introduced systematically.

We developed the stochastic simulation protocol (**Fig. 2d**) to simulate cell cycle lineages by picking up random mother cells at t=0. For each mother cell, at time t=0, we have chosen transcription rates for each cell cycle gene from log-normal distributions, where the mean of the distributions are the mean deterministic transcription rates of the corresponding genes with a specific CV (28) (**Fig. 2d(i)**). The idea behind transcription rate variability of cell cycle genes stems from the recent observations, where it was suggested that cell-to-cell variability in the propensity to transcribe mRNAs plays a dominant role in gene expression fluctuations (28–30). It was observed that in each cell, the transcriptional activity for a gene fluctuates around a gene-specific mean level with a high correlation in mean transcriptional activity between sisters and mother-daughter pairs, suggesting the existence of transcriptional memory across generations (29, 30). These studies claim that the daughter cells inherit certain factors controlling transcription in a correlated manner but at the same time transcribe genes quite differently from the corresponding mother.

Thus, we added transcription rate fluctuations across cell lineages while implementing Gillespie Stochastic Simulation Algorithm (SSA) to quantify the intrinsic fluctuations (**Fig. S4**) in the expression of genes (**Fig. 2d(ii)**) for the overall cell cycle network (**Table S2** and **Table S3**). Additionally, we have incorporated the extrinsic noise sources in the form of random equal or unequal partitioning of proteins and mRNAs among the two daughter cells during cell division (23)(**Fig. 2d(iii)**), and transcription rate variability of the cell cycle regulatory genes in different phases of the cell cycle (**Fig. 2d(iv)**). Here, we assumed that transcription rates get modified due to epigenetic modifications in S-phase (during DNA-synthesis) (63–65)and M-phase (while sister chromatids get segregated among the two daughter cells) (31–36). We followed many generations (for 72 h) in the form of lineage trees (**Fig. 2d**) implementing the above-mentioned simulation protocol (**Fig. 2d** and **Fig. S5**) by picking up a random mother cell with transcription rates of all regulatory genes drawn from log-normal distributions (mean deterministic transcription rate with 20% CV). We assumed that during certain stages of the S-phase and M-Phase, these transcription rates either increase or decrease up to 6-30% (details in **Method** section and **Fig. S5**).

### Model simulations reconciled the experimentally observed cell cycle period and phase duration heterogeneities

The above-mentioned stochastic simulation protocol (**Table S5**) qualitatively reproduced the mean and variances of the T_CC_, T_G1_, and T_S-G2-M_ distributions for similar numbers of numerically simulated single cells by following several cell lineages under both 2% (**Fig. 3a** and **Fig. S6a-b**) and 10% (**Fig. 3b** and **Fig. S6a-b**) serum conditions. The numerical simulation displayed that under both the serum conditions, the CV of the T_CC_ slightly decreases at 10% serum, however, T_G1_ distribution CV decreases, and T_S-G2-M_ distribution CV either increases or stays the same as the serum dose increases from 2% to 10% (**Fig. 3c**). Intriguingly, even in our numerical simulations, we observe that at 2% serum condition, T_CC_ correlates to a higher extent with T_G1_ in comparison to T_S-G2-M_ (**Fig. 3d** and **Fig. S6c-d**), whereas T_CC_ correlates more with the T_S-G2-M_ at 10% serum (**Fig. 3e** and **Fig. S6c-d**). Moreover, the model predicts that the ratio (R) of the CV ‘s of the T_G1_ and T_S-G2-M_ distributions (over three simulation replicates, **Table S5**) relatively increases by ∼21% as the serum level changes from 2% (R_2%_ = 0.76) to 10% (R_10%_ = 0.92). Interestingly, we have observed similar features in the experimental data (over four experimental replicates, **Table S1**, and **Fig. 1f**) as well, where the relative increase in R is ∼21.3% as the serum level goes from 2% (R_2%_ = 0.61) to 10% (R_2%_ = 0.74). Thus, our model qualitatively captures the relationship between the changes in the correlation and fluctuation pattern of the cell cycle phase duration distributions as the serum level varies from 2% to 10%. This analysis suggests that by altering the fluctuation pattern of these phase duration distributions, one can shift the correlation pattern between T_CC_ Vs T_G1_ and T_CC_ Vs T_S-G2-M_ and vice-versa. How to achieve such kind of alteration and what will be its implications, remained an open question.

**Fig. 3.**
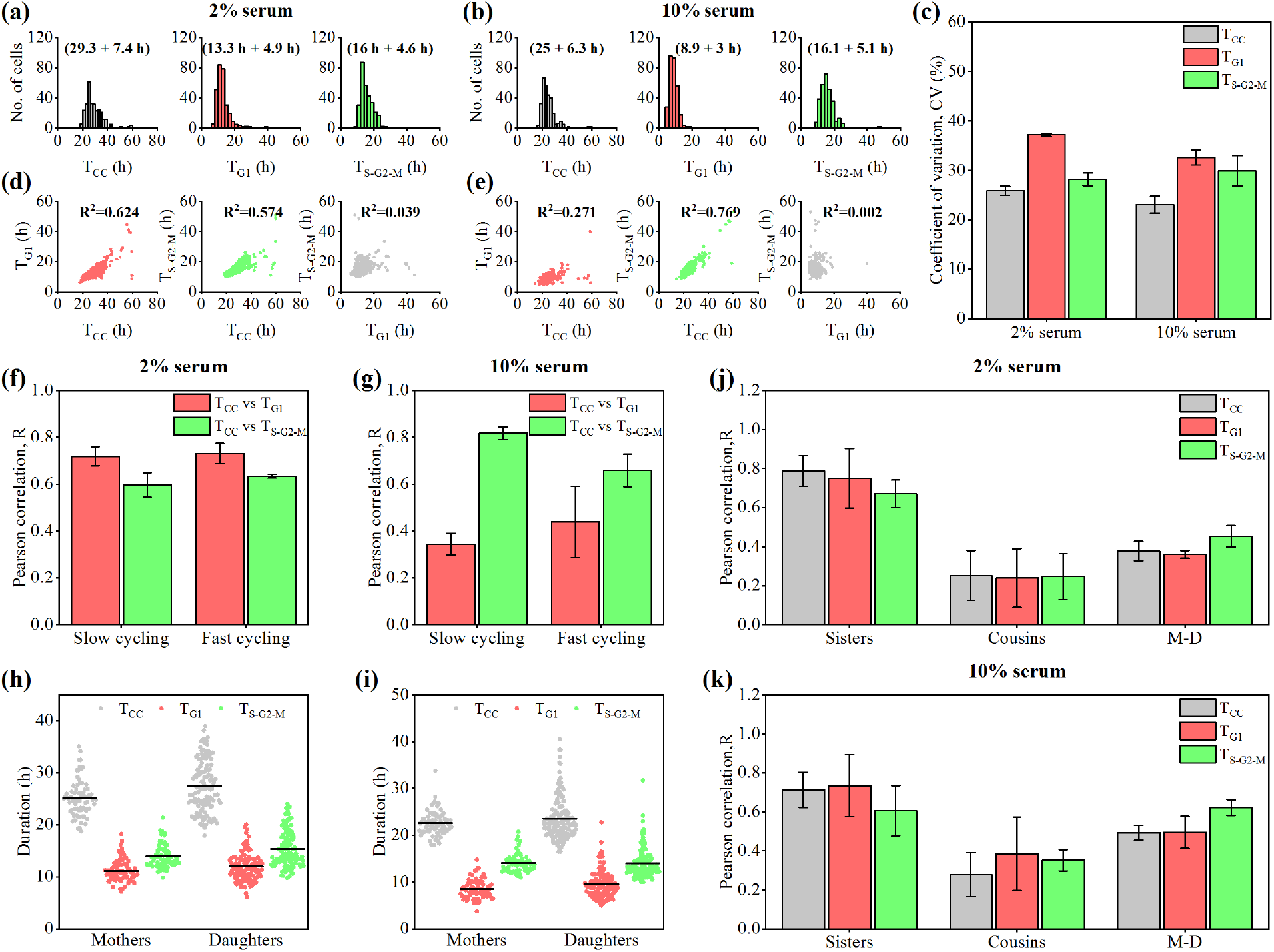
Stochastic model simulations corroborate the experimentally observed T_CC_, T_G1_ and T_S-G2-M_ heterogeneities. Distributions of T_CC_, T_G1_ and T_S-G2-M_ obtained from stochastic simulation at (**a**) 2% serum (n=300, replicate 1) and (**b**) 10% serum (n=300, replicate 1). (**c**) Average CV in T_CC_, T_G1_, and T_S-G2-M_ was obtained at 2% serum and 10% serum (from 3 replicate simulations, 300 cells each), respectively. Correlation pattern of T_CC_, T_G1_ and T_S-G2-M_ at (**d**) 2% serum (n=300, replicate 1) and (**e**) 10% serum (n=300, replicate 1). Mean and standard deviation of correlation between T_CC_, T_G1_ and T_S-G2-M_ in slow and fast cycling cells at (**f**) 2% serum and (**g**) 10% serum. Scatter plot showing the distribution of T_CC_, T_G1_, and T_S-G2-M_ in mothers and daughters (replicate 1) at (**h**) 2% serum and (**i**) 10% serum. Mean and standard deviation of T_CC_, T_G1_, and T_S-G2-M_ correlations quantified for sisters, cousins, and mother-daughter pairs performed at (**j**) 2% serum and (**k**) 10% serum (3 numerical replicates, **Table S6**).

A subpopulation level analysis (**Fig. S2A**) using the model demonstrates that for 2% serum, both slow and fast cycling cells have a high T_CC_ vs T_G1_ correlation (**Fig. 3f**). However, for 10%serum, T_CC_ vs T_S-G2-M_ correlation increases in slow-cycling cells (**Fig. 3g**). Our simulation further exhibits that the cycling time of mothers is comparably faster than the daughters under both 2% and 10% serum conditions (**Fig. 3h-i** and **Fig. S7a-b**). Our model simulation predicts that the difference in cycling time between mothers and daughters is observed due to an increase in cell density and cell-cell contact in the microenvironment under culture conditions (67, 68).

Fascinatingly, our model simulations reproduced that in cell lineages, the T_CC_, T_G1_, and T_S-G2-M_ are highly correlated in sister pairs under both serum conditions, which is substantially higher than the cousin-cousin and mother-daughter pairs (**Fig. 3j-k**) as observed in our experiments. Our model analysis under 2% and 10% serum conditions suggests that the correlations of T_CC_, T_G1_, and T_S-G2-M_ among sisters, mother-daughter, and cousin-cousin pairs are kind of independent of serum conditions. This is in line with our experimental observations (**Fig. 1m-n**) and is in contrast with the previous studies (17, 20, 22) where only a single serum condition was employed to report such correlations. We will elucidate how these correlations remain independent of the serum doses in the coming sections. However, our study, for the first time revealed that such correlations in cell lineage pairs for cell cycle period and phase durations can be explained from a generic stochastic cell cycle network-based model. Our stochastic model performed quite efficiently to reconcile such complex experimental observations and could potentially unravel the crucial factors governing the T_CC_, T_G1_, and T_S-G2-M_ heterogeneities.

### Transcriptional fluctuation governs the cell cycle period and phase duration heterogeneities

The proposed model (**Fig. 2**) includes all the essential deterministic regulations, intrinsic noise, and different extrinsic noise sources, which eventually drive the heterogeneous cell cycle progression in mammalian cells. Thus, it provides the opportunity to decipher the effect of these regulatory processes on T_CC_, T_G1_, and T_S-G2-M_ heterogeneities by considering different reduced versions (**Table-1**) of our initially proposed full network-based stochastic model (**M-1, Fig. 4a** and **Table-1**). First, we performed simulation by considering only intrinsic fluctuations, where each starting mother cell for different cell lineages has the deterministic mean transcription rates for all the genes, and during division is partitioning the molecular regulators unequally among the two daughters (**M-2, Table-1**). Under this scenario, the mean T_CC_ and T_G1_ decrease with the increased serum level without much variation in the T_S-G2-M_. However, this version of the model substantially underestimates (**M-2, Fig. 4b**, and **Table-S6**) the T_CC_, T_G1_, and T_S-G2-M_ variabilities in comparison to the experimental observation (**Fig. 4a**). We observed a bit of sister-sister correlation of T_CC_ at 2% serum, which was absent at 10% serum level. However, no perceptible cousin-cousin or mother-daughter correlations were found at 2% or 10% serum levels (**M-2, Fig. 4b**, and **Table-S6**). This suggests that extrinsic fluctuations in the form of transcription rate variation may significantly affect such a correlation pattern found in cell lineage analysis.

To verify this idea, using a deterministic model (**M-3, Table-1**), we simulated the cell lineages by including the transcription rate variabilities for each starting mother cell for a specific lineage, which during division is partitioning the molecular regulators unequally among the two daughters. Thus, the intrinsic variabilities due to inherent molecular fluctuations are absent in the **M-3** model. **M-3** model shows higher fluctuations in T_CC_, T_G1_, and T_S-G2-M_ (more than **M-2** Model)) with growing transcription rate fluctuations (CV=0% to 40%, for the respective log-normal distributions **Table-S7**) and produces a high correlation in T_CC_ of sister pairs for both 2% and 10% serum conditions (**M-3, Fig. 4c**, and **Table-S6**). Intriguingly, it reveals that if all the cells in different lineages have identical transcription rates, then a highly correlated T_CC_, T_G1_, and T_S-G2-M_ can be observed for sisters, cousins, and even for mother-daughter pairs **(M-3, Fig. 4c**, and **Table-S6)**. This displays that transcriptional variability across cell populations considerably contributes to the fluctuation pattern of T_CC_, T_G1_, and T_S-G2-M_ and causes the high T_CC_ correlations among sister pairs. This demonstrates that transcriptional rates can indeed get inherited (30, 35) by both the daughters from their mother to cause such high correlations.

**Fig. 4.**
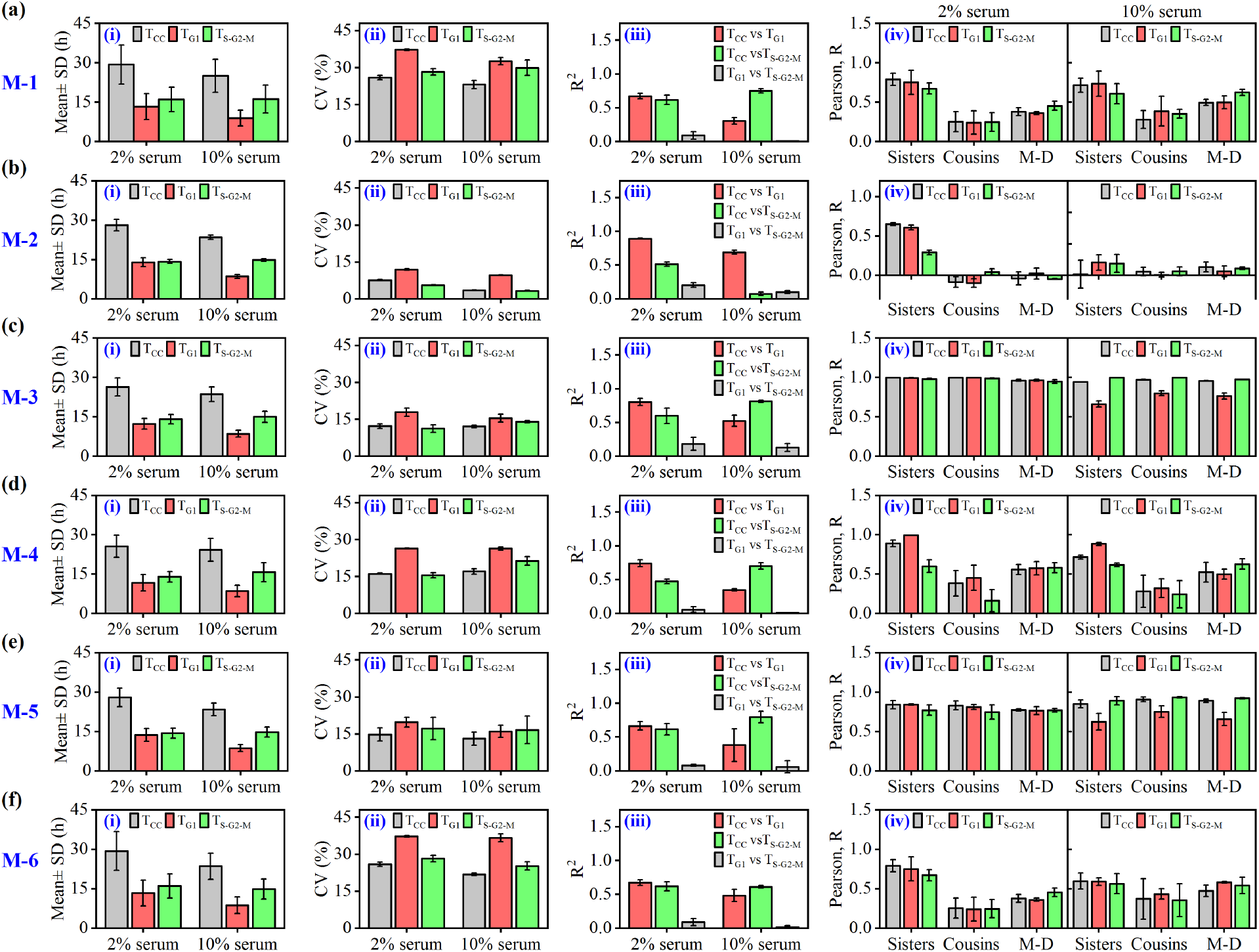
Simulation results at 2% and 10% serum conditions for different model variants with unequal division. Calculated (i) mean T_CC_, T_G1_, and T_S-G2-M_ (± standard deviation (SD) for Replicate I), (ii) mean CV (± SD), (iii) mean correlation coefficient (R^2^) (± SD) of T_CC_, T_G1_ and T_S-G2-M_, and (iv) mean Pearson correlation coefficient (R) (± SD) obtained for the T_CC_, T_G1_, and T_S-G2-M_ for sisters, cousins and mother-daughter pairs selected from different cell lineages for the model variants **(a)** M-1, **(b)** M-2, **(c)** M-3, **(d)** M-4, **(e)** M-5 and **(f)** M-6 defined in **Table-1**.

However, experimentally we do not observe such a high correlation in cousins and mother-daughter pairs, which was confirmed by our full stochastic model (**Fig. 4a** and **Table-S6**). Here comes the importance of the transcription rate variations introduced by us during the S-phase and M-phase, respectively, which take into account the epigenetic modifications (31–36, 63–66) due to genome reorganization during DNA synthesis and separating the sister-chromatids to the respective daughter cells. We numerically included the effect of these events in the model by altering the extent of transcription rate variability for each transcript by using a uniformly distributed random number and adjusted the transcription rates accordingly during S and M-phases, respectively (**Table-S8**, See **Method** for details). Including this feature of transcription rate variation in the **M-4** model (**M-4, Table-1**) leads to a lower correlation of T_CC_, T_G1_, and T_S-G2-M_ among cousins and mother-daughter pairs, without affecting the sister-sister correlations (**M-4, Fig. 4d, Table-S6** and **Table S8**).

This gets further confirmed when we ignored transcription rate variation (**M-5, Table-1**) during the cell cycle in our full model. We observe an increasing level of cousin-cousin and mother-daughter correlation in the **M-5** model simulation (**M-5, Fig. 4e**, and **Table-S6**), which is absent in the experimental findings. This implies that there should be sufficient variation of transcription rates in S and M phases during each cell cycle of every individual cell to create reduced T_CC_, T_G1_, and T_S-G2-M_ correlation in mother-daughter and cousin pairs (**Table S8**). Considering equal partitioning of molecular regulators during cell division (**Table S9** and **Fig. S8**) instead of unequal partitioning only increases the T_G1_ correlation in sisters, cousins, and mother-daughter pairs a bit with a greater increase in T_G1_ correlation at 10% serum **(Table-S9)**.

It is clear from the above analysis that the serum availability has less influence in controlling the correlation of T_CC_, T_G1_, and T_S-G2-M_ of sisters, cousins, and mother-daughters. However, our full model (**M-1, Table-1**) analysis reconciled the fact that serum affects the variability of the individual phase duration distributions considerably (**Fig. 3a-c** and **Fig. 4a**) at the population level, as an increase in serum from 2% to 10%, reduces mean duration and CV of T_G1_ and increases the mean T_S-G2-M_. This consequently reverses the nature of the correlation pattern of T_CC_ with T_G1_ and T_S-G2-M_ (at 2% T_CC_ correlates more with T_G1_, but at 10% T_CC_ correlates more with T_S-G2-M_) (**Fig. 3d-e** and **Fig. 4a**). How change in serum level induces such kind of correlation reversal? To answer this intriguing question, we look back into our proposed cell cycle network (**Fig. 2a**) and find that serum not only activates the initiators (CycD and Myc (69–71)) of the cell cycle in the G_1_-phase, it further activates the Wee1 expression (72, 73) in G_2_ phase. Activation of Wee1 by serum essentially prolongs the T_S-G2-M_, as the serum dose is increased from 2% to 10%. Our reduced model (**M-6, Table-1**) simulation (assuming no serum mediated activation of Wee1) revealed that both the mean T_G1_ and T_S-G2-M_ and related fluctuation decreases, as we increase the serum level from 2% to 10% (**M-6, Fig. 4f**, and **Table-S6**). Consequently, in both 2% and 10% serum conditions, the CV of T_G1_ remains considerably higher than the CV of T_S-G2-M_. This indicates that the serum-dependent cell cycle network functions in a highly coordinated manner to control the overall heterogeneities of the T_CC_, T_G1_, and T_S-G2-M_. At this end, we perform a detailed sensitivity analysis of the model parameters (**Fig. S9**), which demonstrates that the model generated T_CC_, T_G1_, and T_S-G2-M_ change either marginally or moderately, even if we modify any related kinetic parameters of the model. This shows that our model predictions are quite robust.

### Inhibiting p38-signaling alters the cell cycle duration heterogeneities at low serum condition

The observations made from the **M-6** model (**Table 1**) simulations indicate that it may be possible to perturb the serum-dependent cell cycle regulatory network in a specific manner to control the related heterogeneities. Since cells commit to active cell cycle progression during the G_1_-phase of the cell cycle (7), reducing the CV of the T_G1_ distribution for an actively proliferating population of cells under low serum conditions may lead to novel therapeutic insights to get rid of the cancerous cells within a malignant tumor, where cells can proliferate even under serum depleted conditions (7). Literature studies suggest that p38, a stress-signaling protein kinase, retards G_1_ progression by inhibiting the transcription of Myc and CycD and promotes S-G_2_-M progression by activating Plk1 kinase under normal growth conditions in mammalian cells (74–77). Hence, we hypothesize (**Fig. S10a**) that inhibiting p38 protein-mediated signaling at a 2% serum condition may modulate the mean and variance of T_G1_, which will lead to an increase in the correlation of T_CC_ and T_S-G2-M_, as observed under 10% serum condition (**Fig. 3e**).

We incorporated the concept of inhibiting p38 signaling by introducing the effect of thep38-inhibitor in a phenomenological manner (**Table S10**) in our deterministic model (See **Supplementary text** for details) and the model simulation at 2% serum condition shows a decrease and an increase in T_G1_ and T_S-G2-M_, respectively (**Fig. S10b**). The stochastic simulation at 2% serum condition in presence of p38-inhibitor demonstrates reduced mean and variability of T_G1_ (**Table S11**). The model simulation further predicts that the CV of T_CC_, T_G1_, and T_S-G2-M_ at 2% serum condition in presence of p38-inhibition (**Fig. 5a** and **Table S11**) will show a trend as observed for 10% serum condition without any inhibition (**Fig. 1f**). This effect produced in the CV of T_CC_, T_G1_, and T_S-G2-M_ will accompany with a higher correlation between T_CC_ and T_S-G2-M_ (**Fig. 5b**), which is in stark contrast with the control simulation performed at 2% serum condition (**Fig. 5b** and **Table S11**) without p38-inhibitor. At the lineage level, model simulation exhibited a slight drop in the sister-sister T_CC_ correlation due to a drop in the T_S-G2-M_ correlation (**Fig. 5c**) under the p38 inhibition condition, while for the mother-daughter pair, these correlations remain unaffected (**Fig. 5c**). However, the cousin-cousin T_CC_ correlation increases slightly under the p38-inhibition (**Fig. 5c** and **Table S11**) condition due to an increase in the T_S-G2-M_ correlation.

**Table 1.**
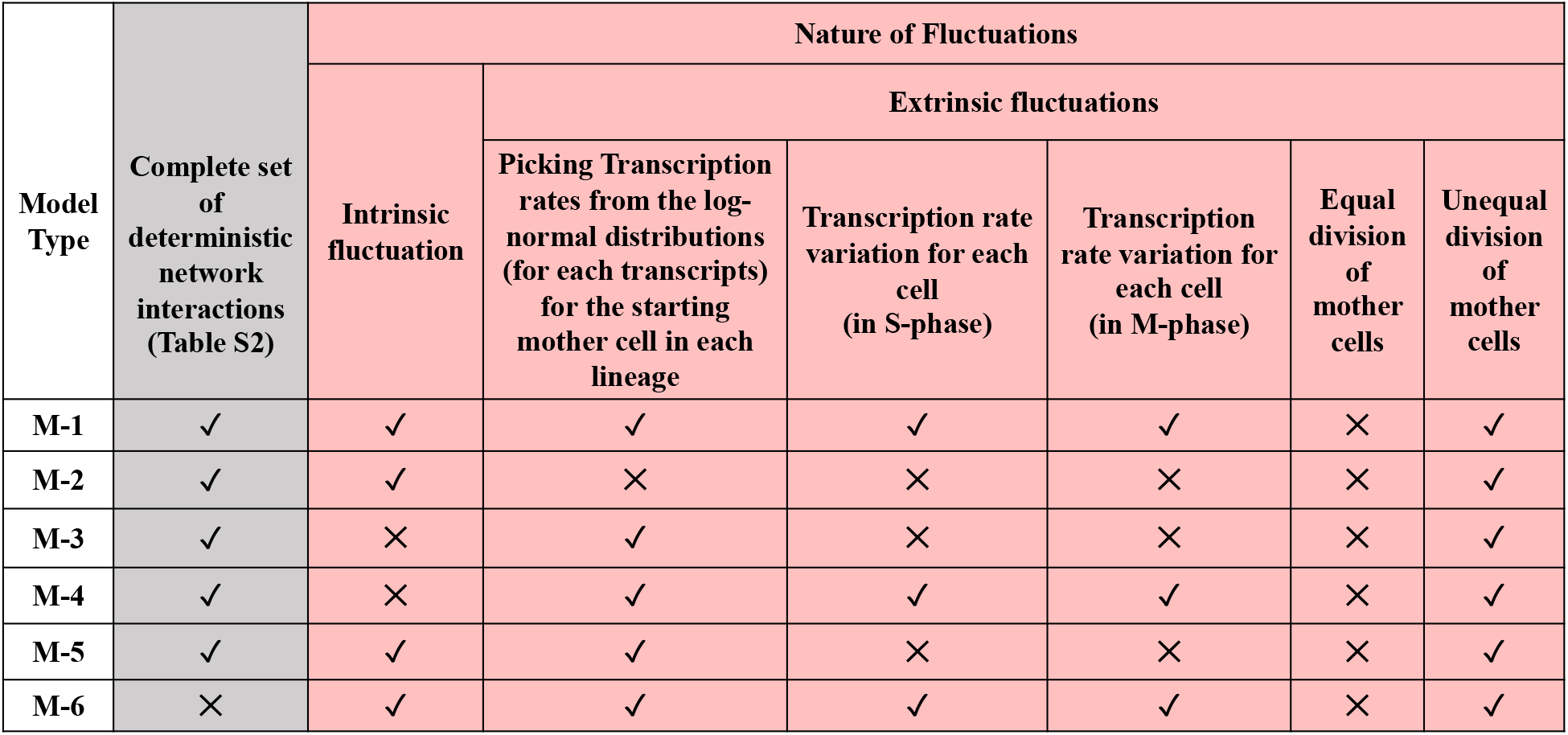
Different model variants are considered to unravel the contributions of various factors in controlling heterogeneities in T_CC_, T_G1_, and T_S-G2-M_ for mammalian cells.

| Model Type | Complete set of deterministic network interactions (Table S2) | Nature of Fluctuations |  |  |  |  |  |
| --- | --- | --- | --- | --- | --- | --- | --- |
|  |  | Intrinsic fluctuation | Extrinsic fluctuations |  |  |  |  |
|  |  |  | Picking Transcription rates from the log-normal distributions (for each transcripts) for the starting mother cell in each lineage | Transcription rate variation for each cell (in S-phase) | Transcription rate variation for each cell (in M-phase) | Equal division of mother cells | Unequal division of mother cells |
| M-1 | ✓ | ✓ | ✓ | ✓ | ✓ | × | ✓ |
| M-2 | ✓ | ✓ | × | × | × | × | ✓ |
| M-3 | ✓ | × | ✓ | × | × | × | ✓ |
| M-4 | ✓ | × | ✓ | ✓ | ✓ | × | ✓ |
| M-5 | ✓ | ✓ | ✓ | × | × | × | ✓ |
| M-6 | × | ✓ | ✓ | ✓ | ✓ | × | ✓ |

**Fig. 5.**
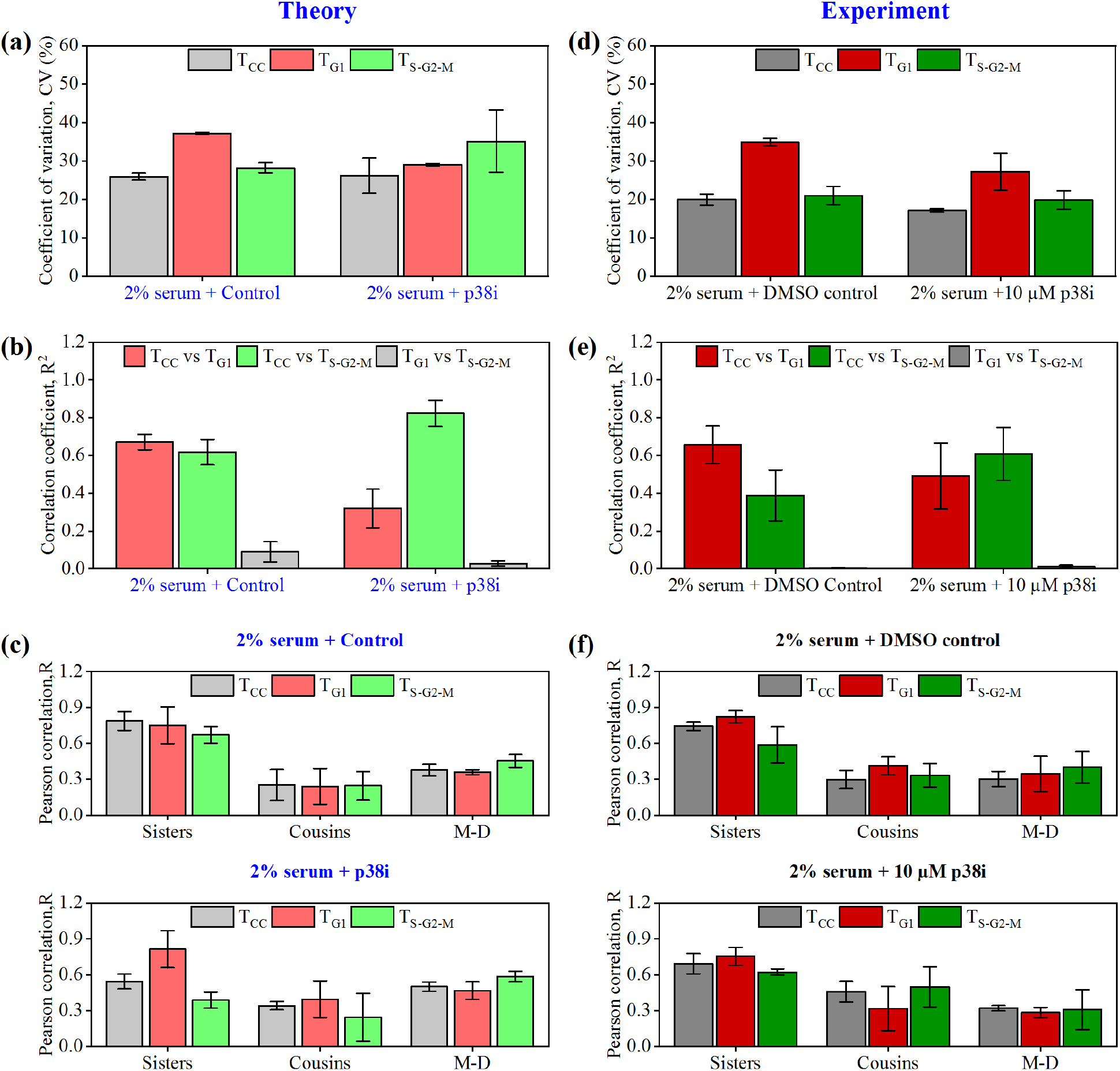
Inhibition of p38-signaling reduces the variability of the G1-phase and reverses the correlation pattern under low (2%) serum conditions. Numerically simulated 300 cells under only 2% serum (control) and 2% serum + p38-signaling inhibitor (10 μM). **(a)** mean CV (± SD) of T_CC_, T_G1_, and T_S-G2-M_ for all 3 numerical replicates, **(b)** mean Correlation coefficients (± SD) of T_CC_, T_G1_, and T_S-G2-M_, and **(c)** mean Pearson correlation (± SD) of T_CC_, T_G1_ and T_S-G2-M_ calculated from cell lineages for sisters, cousins, and mother-daughter pairs. Live-cell image analysis of nearly 300 FUCCI-HeLa cells at 2% serum (control) and 2% serum with 10 μM p38-signaling inhibitor. **(d)** experimental mean CV (± SD) of T_CC_, T_G1_, and T_S-G2-M_ for all 3 replicates, **(e)** experimental mean Correlation coefficients (± SD) of T_CC_, T_G1_, and T_S-G2-M_, and **(f)** experimental mean Pearson correlation (± SD) of T_CC_, T_G1_, and T_S-G2-M_ calculated from cell lineages for sisters, cousins, and mother-daughter pairs.

To verify our model predictions, we cultured FUCCI-HeLa cells at 2% serum along with ap38 inhibitor and performed live-cell imaging. The analysis of live-cell imaging revealed a decrease in mean T_G1_ with an increased mean T_S-G2-M_ **(Table S12)** under the p38 inhibitor condition compared to the control cell population, where the T_G1_ variability decreases **(Table S12)**. Moreover, experimentally quantified CV of T_G1_ at 2% serum condition in presence of p38-inhibition is considerably lower than the control situation as predicted by our model simulation (**Fig. 5d** and **Table S12**). Intriguingly, under 2% serum condition, T_CC_ vs T_G1_ correlation is found to be lower than the T_CC_ vs T_S-G2-M_ in the presence of p38 inhibitor **(Fig. 5e** and **Table S12)**, which is in agreement with the model prediction. Interestingly, under 2% serum along with p38-inhibitor, the T_CC_, T_G1_ and T_S-G2-M_ correlations among the sisters and mother-daughter pairs remain similar to that observed under the control situation **(Fig. 5f** and **Table S12)**. However, the cousin-cousin pair shows a lower correlation under the p38-inhibitor condition **(Fig. 5f** and **Table S12)**. Thus, the p38-inhibition study under 2% serum displays that T_CC_, T_G1_, and T_S-G2-M_ heterogeneities at the overall cell population level can be fine-tuned to create lower variability in T_G1_, which may have therapeutic relevance to get rid of the unwanted carcinogenic cells within a tumor.

## Discussion

Identifying the governing factors that organize the cell duration heterogeneities is a challenging task (8, 9, 12, 17, 19–22). In this article, we combined live-cell imaging studies of FUCCI-HeLa cells (**Fig. 1**) with an appropriate cell cycle network-based stochastic mathematical model (**Fig. 2**) and unraveled the major factors that control the heterogeneities associated with the T_CC_, T_G1_, and T_S-G2-M_ in a serum-dependent manner (**Fig. 6**). First, we demonstrated that the transcription rate variability in different cell lineages present within the cellular population is responsible for the high correlations in T_CC_ among sister pairs across cell lineages (**Fig. 6a**, and **Fig. 3-4**). By employing a specific model variant (**M-4** Model, **Table-1**), we revealed that the transcription rates alteration due to epigenetic modifications during S-phase and M-phases lead to a moderate to low correlation in cousins and mother-daughter pairs without affecting the high sister-sister correlation (**Fig. 6a**, and **Fig. 4d**). These findings substantiate the idea that cell-to-cell variability in the transcription of mRNA for different regulatory genes is the major source of variability in mammalian cells (28–30).

**Fig. 6.**
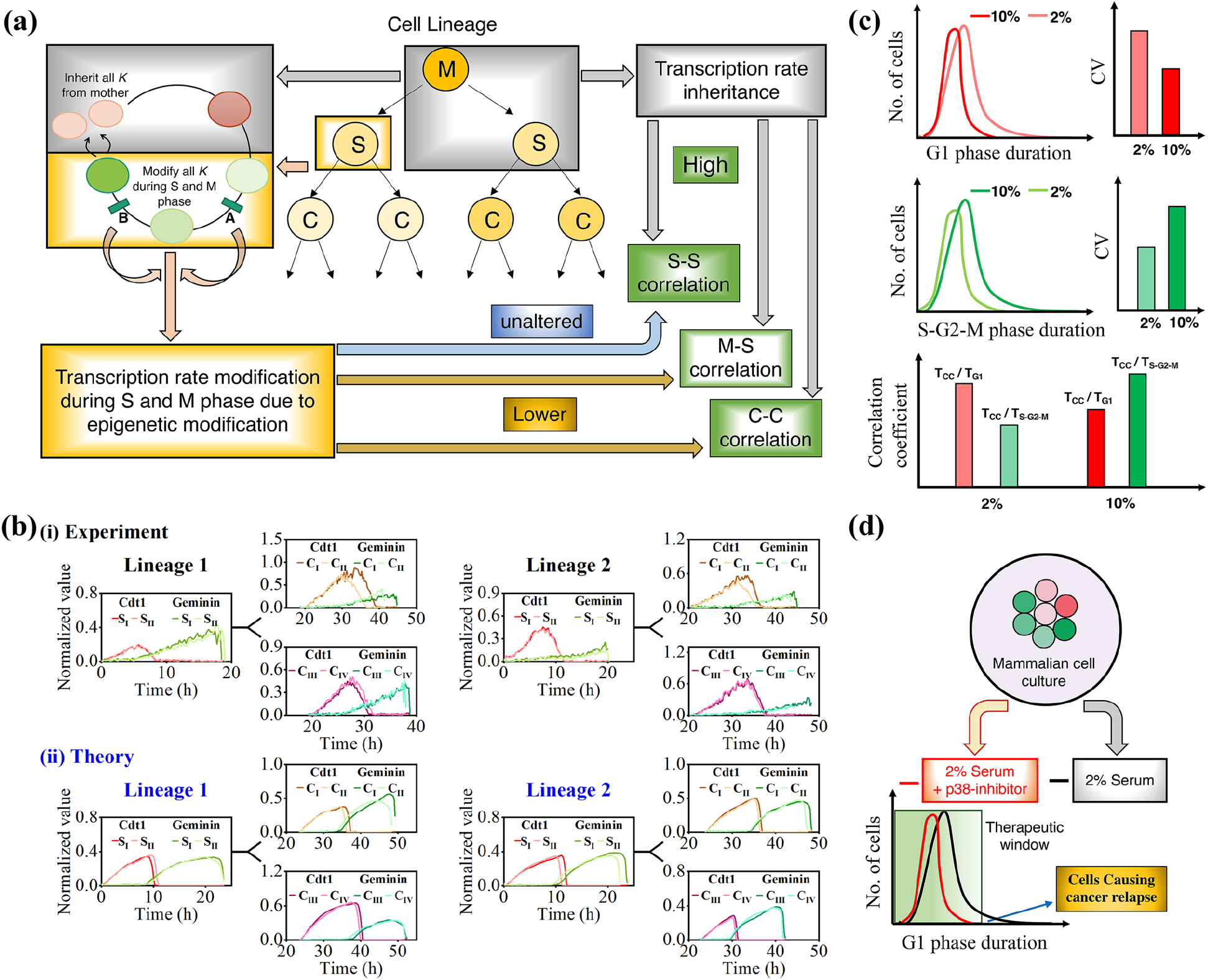
Transcription rate variation across cell lineages and during cell cycle phases govern the cell duration heterogeneities. **(a)** Schematic depiction of how transcription rate variation across cell lineages and during cell cycle orchestrate the correlation of T_CC_, T_G1_, and T_S-G2-M_ in sisters, cousins, and mother-daughter pairs. **(b)** Correlated transient dynamics of Cdt1 and Geminin during G1-phase which often gets decorrelated during S and M-phases; **(i)** Experiment, and **(ii)** Theory. **(c)** Schematic depiction of the alteration in mean, SD, and CVs of the T_G1_ and T_S-G2-M_ due to serum doses and its effect on correlation patterns of T_CC_ vs T_G1_ and T_CC_ vs T_S-G2-M_. **(d)** Inhibition of p38-signaling can lower the mean and variance of G1 phase duration even under low serum conditions.

Further, these studies suggested that daughter pairs inherit the transcription rates from the respective mother in a correlated manner (28–30), and it causes the correlated transcription pattern of various genes during the G_1_-phase among the daughter pairs. Our analysis with Cdt1 and Geminin time courses for two representative experimental (**Fig. 6b(i)**) and theoretical (**Fig. 6b(ii)**)cell lineages revealed that the high sister-sister correlation is indeed emanated from the highly correlated transcriptional activity of the cell cycle genes (we showed Cdt1 and Geminin) during G_1_-phase, due to inheritance of same transcriptional rates for all genes by both the sisters from their mother. However, mothers T_CC_ remain less correlated with the daughters T_CC_ as mothers only spent a few hours with the transcription rate after the epigenetic modification in M-phase, which are inherited by the respective daughters during cell division. Any loss in the correlation between sisters arises from intrinsic molecular fluctuations and unequal molecular partitioning during cell division (**Fig. 4**). Due to epigenetic modifications, the transcriptional rates get altered during the S and M phases, and the respective daughters come up with different transcription rates for all the genes before the next cell division. However, this effect leads to a significant drop in the cousin-cousin correlation (**Fig. 1m-n, Fig. 3j-k**, and **Fig. 6a-b**) as the extent of epigenetic modifications is different in respective sisters (**Fig. 6b**). It also caused the desynchrony in T_S-G2-M_ (**Fig. 6b**) and little loss of T_CC_ correlation among sister pairs. This observation is in contrast with that made by Dowling et al. (17) where they concluded that cells spend a near equal proportion of T_CC_ as T_S-G2-M_ and that creates the correlation among sibling pairs. These are important model predictions that can be experimentally probed in the future by performing in-situ epigenetic modifications in the cycling population of cells. Interestingly, the model simulation and experiments showed that the change in serum level did not modify the above-mentioned correlations appreciably (**Fig. 1m-n**, and **Fig. 3j-k**) among sisters, cousins, and mother-daughter pairs, demonstrating the robust nature of these correlations. Thus, our model provides a highly plausible and realistic explanation of the cell lineage pair correlations from a cell cycle network-based dynamical model.

Second, we captured the subtle effect of serum modulation on the overall cellular population of FUCCI-HeLa cells. We observed that shifting the serum level from 2% to 10% caused a decrease and an increase in mean, variance, and CV of T_G1_ and T_S-G2-M_, respectively (**Fig. 6c** and **Fig. 1d-f**), which lead to higher T_CC_ vs T_S-G2-M_ correlation at 10% serum in comparison to higher T_CC_ vs T_G1_ correlation at 2% serum (**Fig. 6c** and **Fig. 1g-h**). These observations are quite counterintuitive and in contrast with what had been reported for budding yeast cell cycle (78, 79), where an increase in glucose dosage in media increases the number of proteins and mRNAs within the cells, which leads to a decrease in the mean as well as variances of T_CC_, T_G1_, and T_S-G2-M_. Our model predicted that for mammalian cells, a coordinated effect of serum mediated activation of CycD and Myc in the G_1_-phase and Wee1 in the S-G_2_-M phase, and the extrinsic noise due to transcription rate fluctuation across cell lineages and during the S and M-phases during cell cycle progression produced such kind of unique fluctuation and correlation pattern (**Fig. 3a-e**). The slowly growing cells mostly dictated the overall variability pattern of T_G1_ and T_S-G2-M_ for the overall cellular population under different serum levels (**Fig. 1i-j** and **Fig. 3f-g**), however, the mother ‘s T_CC_ was always faster than the daughter ‘s T_CC_ (**Fig. 1k-l** and **Fig. 3h-i**) as observed in case of budding yeast cells (78).

Third, we showed that it is possible to perturb the cell cycle regulatory network to achieve a lower mean and variance of T_G1_ for a proliferating population of the cell under lower (2%) serum conditions by employing a p38-signaling inhibitor (**Fig. 6d** and **Fig. 5**). This may lead to effective therapeutic strategies to get rid of the slowly proliferating cells within the tumor population, as cells in tumors do survive and some of them evade chemotherapies due to higher T_G1_ variability under minimal growth conditions (**Fig. 1d** and **Fig. 5d**). Cells in presence of p38-inhibitor (with 2% serum) behaved as if they were under 10% serum condition, and demonstrated a signature high T_CC_ vs T_S-G2-M_ correlation (**Fig. 5e**). However, the correlation of T_CC_, T_G1_, and T_S-G2-M_ for cell lineage pairs remained grossly unaltered even under inhibitory conditions (**Fig. 5f**). Importantly, our stochastic model simulation adequately captured these features of T_CC_, T_G1_, and T_S-G2-M_ heterogeneities (**Fig. 5a-c**). This demonstrates that the T_CC_, T_G1_, and T_S-G2-M_-related heterogeneities can be influenced by just intervening at the cellular network level. Finally, by employing different model variants (**Table-1**), we showed that intrinsic fluctuations hardly affected the variances of T_CC_, T_G1_, and T_S-G2-M_ due to high copy numbers of mRNAs and proteins in mammalian cells (55–58), but modified the extent of variabilities originated due to transcription rate modifications (**Fig. 4** and **Table-S6**).

In conclusion, our modeling study systematically establishes that transcription ratevariation across cell lineages and during cell cycle progression majorly governs the cell cycle period and phase duration heterogeneities both at the cell lineage and overall population level. This contrasts the ideas of previous studies, where either it was suggested that circadian clock mediated cell cycle modulation (12, 20), or hidden long-range memories of growth and cell cycle period(22) caused the cell cycle duration heterogeneities across cell lineages. In our study, we considered a realistic cell cycle network model and addressed various facets of cell duration heterogeneities both at the cell lineage and at the overall population level under different serum conditions to elucidate the contribution of both stochastic and deterministic effects that organized the cell cycle duration heterogeneities. We demonstrated that the T_CC_, T_G1_ and T_S-G2-M_ related correlation in the cell lineage pairs remained almost unaltered under external perturbations suggesting that these features of cell cycle duration heterogeneities were extremely robust. However, for the overall cellular population, the mean and variances of T_CC_, T_G1_, and T_S-G2-M_ can be systematically altered by perturbing the cell cycle regulatory network. We believe that these insights will invoke new ideas to alter the mammalian cell cycle period and phase durations advantageously to produce novel therapeutic strategies.

## Materials and Methods

### Cell culture

HeLa cells and FUCCI -HeLa cells were cultured in DMEM medium (Hi Media, AT007) supplemented with 10% FBS (Gibco) and 1% Penicillin-Streptomycin (Hi Media,) at 37°C and 5% CO_2_.

### Generation of a stable cell line expressing FUCCI

Lentiviral plasmids containing the FUCCI genes -mKO2-hCdt1 (30/120) and mAG-hGeminin (1/110) were kindly gifted by Prof Atsushi Miyawaki, Lab for Cell Cycle Dynamics, RIKEN Brain Science Institute, Japan. The packaging and envelope plasmids (pCAG-HIVgp and pCMV-VSV-G-RSV-REV) were purchased from RIKEN BRC DNA BANK, Japan, and were used for lentivirus production. Plasmids containing mKO2-hCdt1 (30-120) and mAG-hGeminin (1-110) were co-transfected with envelope and packaging plasmids into LentiX-293T cells (Takara bio) to generate lentiviral particles. High-titer viral solutions were prepared and used for transduction. HeLa cells were first transduced with the virus-containing mAG-hCdt1 (30-120) gene and those cells which express RFP were sorted using FACS. Next, transduction was done on these sorted RFP positive cells using the virus-containing mKO2-hGem (1-110) gene. FACS sorting was done based on RFP fluorescence to obtain double-positive cells expressing FUCCI.

### Live cell imaging and data extraction

For live-cell imaging, on day 1, ∼ 10,000 FUCCI expressing HeLa cells were seeded per well of 6 well plates in opti-MEM media supplemented with FBS (2% or 10%) and 1% penicillin-streptomycin. On day 2, live-cell imaging was done using Zeiss Observer Z1 inverted fluorescence microscope fitted with a high-speed microlens-enhanced Nipkow spinning disc (Yokogawa CSU-X1 automated model) in a temperature (37°C) and CO_2_ (5%) controlled incubation stage. Ahalogen lamp is used as a light source along with Alexafluor 488 and Rhodamine filters. 10 positions were selected and images were taken every 15 min interval for all the 10 positions selected for 3 days.

Zen software was used to process the .czi files and create .tiff images for tracking. To obtain FUCCI trajectory in single cells, each cell was tracked by clicking the center of the nucleus using the ‘Manual tracking ‘ plugin available in ImageJ software. The positions (x, y) of the tracked cell are stored as a text file. An ImageJ script was written to create ROI using the x, and y coordinates and measure the mean fluorescence intensity at each channel for each cell. The duration between birth and the next division gives the cell cycle duration for a cell. The time corresponding to maximum RFP intensity is considered as the end of G_1_. The S-G_2_-M duration is calculated by subtracting G_1_ time from total cell cycle time. The distribution graphs were plotted using Origin software.

### Deterministic simulation

The differential equations used for deterministic simulation (**Table S3**) and the parameter values (**Table S4**) are given in the **supplementary material**. Using these, we have simulated the cell cycle model using a CVODE solver in XPPAUT 8.0 software and obtained periodic oscillation of the cell cycle network components for 2% and 10% serum levels (the ODE file will be available upon request). In this regard, we have tried to keep the number of proteins and mRNA in accordance with that reported for mammalian cells and performed some preliminary bifurcation analysis to set the deterministic model in the appropriate parametric regime.

### Simulation in cell lineages

To follow cells in lineages in our simulation similar to experiments, we developed an algorithm (**Fig. S5)**, where we followed cells from Generation 0 to at most Generation III in each lineage for ‘n ‘ lineages during a 72-h simulation period. To perform stochastic simulation of models-M-1, M-2, M-5, and M-6 **(Table-1)**, ‘Direct method of Gillespie algorithm ‘ (80) was used by incorporating the various sources of extrinsic variabilities (**Fig. 2d)** as described in **Table-1**. The details protocol for the simulation method is provided in **Fig. S5** and the corresponding figure legend. The code for stochastic simulation was written in Fortran language (the code will be available upon request). By adopting a similar lineage simulation algorithm, we investigated the role of various extrinsic noise sources (**Fig. 2d)** in the absence of intrinsic fluctuations, by developing a Matlab code (the code will be available upon request) for the models-M-3 and M-4 **(Table-1)**. For these models, the differential equations (**Table S3**) were solved using Matlab ODE solver (ODE15s) with the parameters mentioned in **Table S4**.

To obtain cell cycle period, G_1_, and S-G_2_-M duration distributions, we collect the cell cycle and phase timings from all the cells that had a complete cycle during the simulation period of 72 hrs. The cell cycle time is calculated using the birth and division time for each cell. Our simulation also keeps track of Cdt1 and Geminin dynamics during the simulation for each cell. The time difference between birth and the Cdt1 peak is considered as the end of G_1_. The S-G_2_-M time is calculated from the total cell cycle period and G_1_ time. For distribution graphs and variability calculation under different serum conditions, all the cells with a complete cycle from all the lineages were considered. All possible mother-daughter, sister, and cousin pairs with complete cell cycle duration irrespective of the generation was used to determine correlations in the cell cycle period and phase durations.

## Acknowledgments

We thank Prof Atsushi Miyawaki, Lab for Cell Cycle Dynamics, RIKEN Brain Science Institute, Japan for generously providing FUCCI plasmids. We thank Cell Culture Facility at the Department of Chemistry, IIT Bombay, the Spinning Disc Confocal Microscope facility at BSBE, IIT Bombay, and FACS facilities at BSBE and CRNTS, IIT Bombay for allowing us to use them for this work. We thank IIT Bombay for providing the TA fellowship to VG. This work is supported by the funding agencies **DBT, India** (Grant no. **BT/PR11932/BRB/10/1315/2014**) and **SERB, India** (Grant no. **CRG/2019/002640**, Grant no. **MTR/2020/000261** and Grant no. **EMR/2014/000500**).

