## Supplementary file for "Transcriptional fluctuations govern the serum dependent cell cycle duration heterogeneities in Mammalian cells"

### SI Text

#### Detail description of the proposed cell cycle network

The proposed cell cycle network consists of three modules to represent the cell cycle progression for a mammalian cell. All the biochemical reactions (provided in **Table S2**) governing the interactions in **Fig. 2a** are curated from existing literature on mammalian cell cycle regulation (1–12). Reactions governing the interactions between network components in the proposed network (**Fig 2**) are described in details in **Table S2**. The differential equations for each molecular species (**Table S3**) are written mostly using mass-action kinetics with the exception of Michaelis-Menten and Hill-kinetics terms for some activation and degradation processes which are described in a phenomenological manner. Half-lives of some mRNAs and proteins are taken from literature and converted into respective rate constants (**Table S4**). However, most of the other kinetic parameters in our model are adjusted to recreate the experimentally observed average cell cycle period, $G_1$ and S-$G_2$-M durations under different serum conditions, where we captured the reported dynamics of different proteins and mRNA as per available experimental literature phenomenologically. Reactions provided in **Table S2** can then be readily used for stochastic simulation by employing Gillespie's stochastic simulation algorithm (SSA).

**Construction of Module I:**

The first module is the restriction point module that senses the amount of serum and makes the decision for cell cycle commitment (2, 4, 12). Serum stimulation induces the transcription of Myc mRNA via an upstream signalling protein MV. Myc mRNA is translated to form Myc protein, which in turn activates transcription of E2F1 mRNA. E2F1 protein translated from its mRNA forms a heterodimeric complex DE with its partner Dp1 protein and serves as transcription factor for its own production and also synthesis of several proteins such as Myc, Cyclin E and Cyclin A to facilitate S phase entry and progression (2, 9, 13–17). However, during $G_1$ phase Rb protein translated from Rb mRNA inhibits DE by forming Rb:DE complex and DE will be released from the complex only when Rb inactivation occurs due to phosphorylation of Rb initially by Cyclin D:Cdk4 and later by Cyclin E:Cdk2 complexes (2, 9, 14, 15, 18, 19). Serum and Myc promotes transcription of Cyclin D mRNA and the translated Cyclin D protein forms complex with Cdk4 to form Cyclin D:CdK4 (represented as CycD in our model) and phosphorylates free Rb and Rb bound to DE complex leading to partial activation of E2F1. Additionally, E2F creates positive feedback for its activation by promoting its own transcription and also promoting transcription of Myc and Cyclin E to escape from Rb mediated repression. While monophosphorylated Rb can still bind to DE (iDE complex), the second phosphorylation on monophosphorylated Rb done by Cyclin E:Cdk2 leads to complete release and activation of E2F1. After S phase entry, the levels of Myc and Cyclin D:Cdk4 have to go down to low levels during mid and late S phase. This is achieved by the activation of $SCF^{Skp2}$ and Cyclin A, respectively (20, 21). E2F1 level also goes down due to Cyclin A:Cdk2/1 mediated phosphorylation and eventual degradation (22, 23). All the above reactions (R1-R47) are modelled in Equations 2-16 (**Table S3**).

**Construction of Module II:**

DE promotes transcription of Cyclin E and Cyclin A, that forms complex with their respective Cdk partners to facilitate S phase entry and progression. However, during $G_1$ phase, Cyclin Kinase Inhibitors (p27 and p21, denoted as CKI in our network) translated from CKI mRNA are present in high levels and inhibits Cyclin E:Cdk2 activity by forming Cyclin E:Cdk2:CKI complex (11, 19, 24, 25). On the other hand, Cyclin A:Cdk1/2 is subjected to $APC^{Cdh1}$ mediated degradation (7, 27) Activation of Cyclin E:Cdk2 can happen only when CKIs are subjected to ubiquitin mediated degradation during late $G_1$ phase by the E3 ligase $SCF^{Skp2}$ (26, 28).Skp2, a cofactor for SCF is translated from Skp2 mRNA and is not available for binding until late $G_1$ due to $APC^{Cdh1}$ mediated degradation in $G_1$ phase (28). Thus, free Cyclin E:Cdk2 accumulated in small amounts creates positive feedback loop for its release from Cyclin E:Cdk2:CKI complex by two different ways: (i) phosphorylates CKIs directly and facilitates their $SCF^{Skp2}$ mediated degradation, (ii) phosphorylates Skp2 and prevents it from $APC^{Cdh1}$ mediated degradation (29, 30). Cyclin A translated from Cyclin A mRNA phosphorylates Cdh1 and causes inactivation of $APC^{Cdh1}$ to relieve Cyclin A:Cdk2/1 and Skp2 from $APC^{Cdh1}$degradation (27, 31). Thus, Cyclin E and Cyclin A work together to overcome the inhibition CKI and Cdh1 and facilitate S phase entry. After S phase entry, CycE:Cdk2 complex is degraded by $SCF^{Skp2}$ (32, 33). All the above reactions (R48-R86) are modelled in Equations 15-28 (**Table S3**).

**Construction of Module III:**

The final module controls the entry into mitosis followed by cell cycle exit (1, 5, 8, 10, 34). After S phase entry, accumulated CycA:Cdk2/1 activates transcription of Cyclin B mRNA, which gets translated to form Cyclin B protein, which forms CycB:Cdk1 complex (35). However, Cyclin B:Cdk1 complex will be phosphorylated immediately by Wee1 kinase transcribed from Wee1 mRNA, whose synthesis is dependent on Serum. Cyclin A:Cdk2/1 also activates the transcription of Cdc25 mRNA which gets translated to Cdc25 protein. However, Cdc25 phosphatase is inactive until it gets phosphorylated by CycB:Cdk1. CycB:Cdk1 slowly accumulates during S phase and creates a positive feedback loop for its own activation by phosphorylating Cdc25 and Wee1 to activate and inactivate them via dephosphorylation and phosphorylation respectively (5, 10, 34, 36, 37). Entry into the mitosis is marked by switch-like activation of CycB:Cdk1. During end of mitosis, Cyclin B:Cdk1 creates a negative feedback with a delay by activating Cdc20 which dephosphorylates and activates its inhibitor, Cdh1(1, 8). The exit from the cell cycle happens when the cell cycle activators level go down and the inhibitors are activated again. As a result, an increase in the levels of Rb and CKI occurs during late mitosis. This event marks the end of mitosis and hence the cell cycle exit. Since, Cdh1 and Skp2 oscillates antagonistically it leads to the oscillatory dynamics of their targets, Geminin and Cdt1, respectively. All the above reactions (R87-R129) are modelled in Equations 26-45 (**Table S3**). We must emphasize that we have taken the minimalistic interactions that can lead to the exit module. However, there are more interactions involved in the actual mammalian cell cycle regulation which can make it even more perfect model but will contains more numbers of kinetic parameters. Thus, we worked with this minimal network model.

**Interactions governing p38 mediated cell cycle regulation:**

During $G_1$ phase of the cell cycle, activation of p38 activates the phosphatase PP2A which in turn causes ERK dephosphorylation and inactivation which in turn can cause decreased synthesis of proteins like Myc and Cyclin D (38). Hence, p38 decreases Cyclin D protein levels by negatively regulating Cyclin D transcription (39), phosphorylating and degrading Cyclin D protein (**Fig. S10A**) (40–42). p38 also activates cyclin kinase inhibitors such as p16, p21, p27 (**Fig. S10A**) and further inhibits $G_1$ progression and promotes cell cycle arrest (43, 44). We have incorporated the effect of p38 inhibition using two phenomenological terms in Cyclin D synthesis and CKI synthesis (**Table S10**).

Studies have shown that normal p38 is needed for proper spindle formation and activity timely progression through mitosis (45–47). Inhibition of p38 activity is known to cause defective spindle formation and mitotic arrest. We incorporated the effect of p38 inhibition by using a phenomenological inhibition term in synthesis of Cdc25 (**Table S10**), a positive regulator of mitotic progression.

#### Supplementary figures

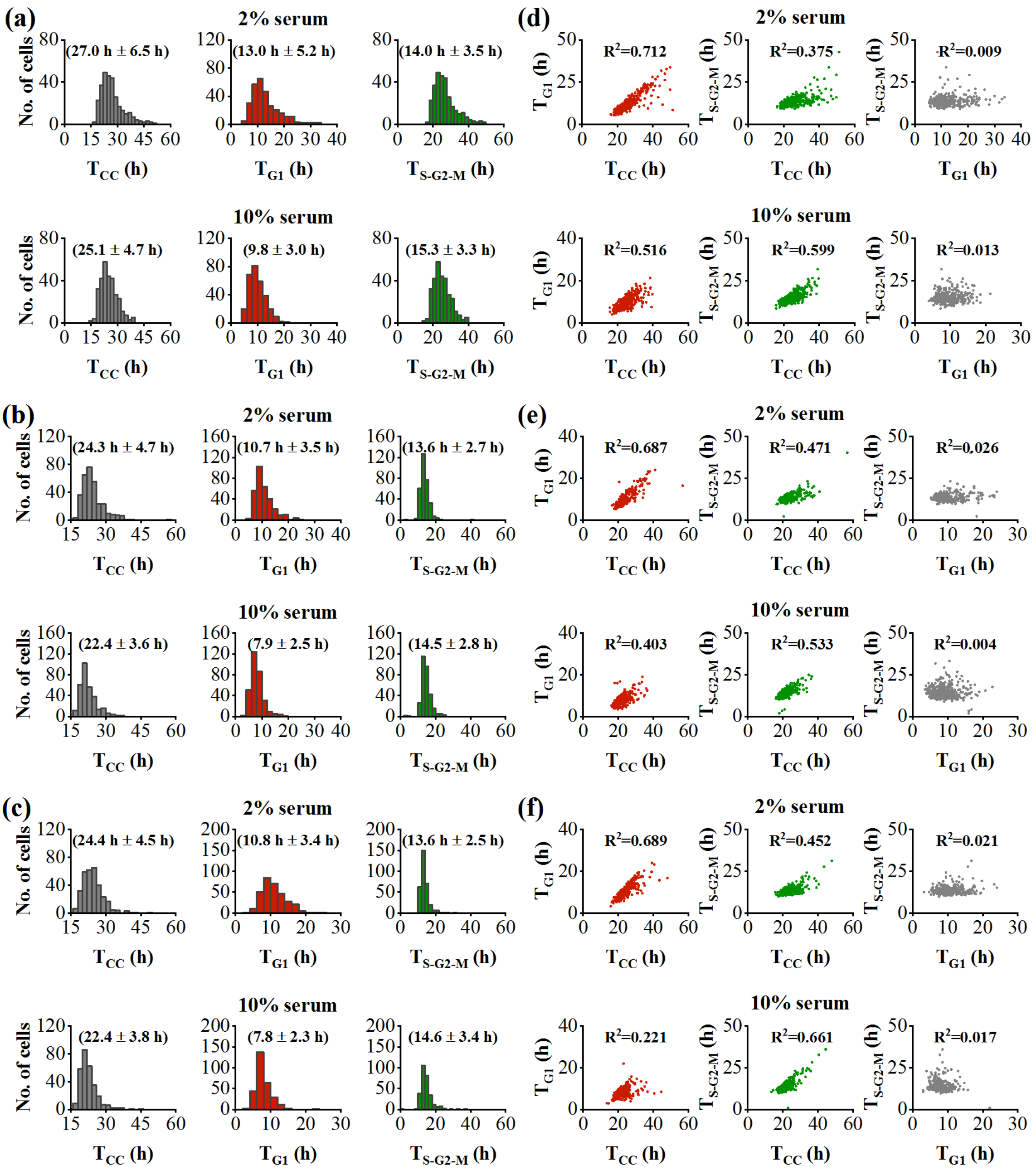


**Fig. S1.** Cell cycle, $G_1$ and $S$-$G_2$-M distribution quantified from single cell imaging data of FUCCI-HeLa cells at 2% serum and 10% serum for (a) replicate II, (b) replicate III and (c) replicate IV. Correlations among cell cycle period and phase durations observed from the single cell imaging data of HeLa at 2% serum and 10% serum for (c) replicate II, (d) replicate III and (e) replicate IV.

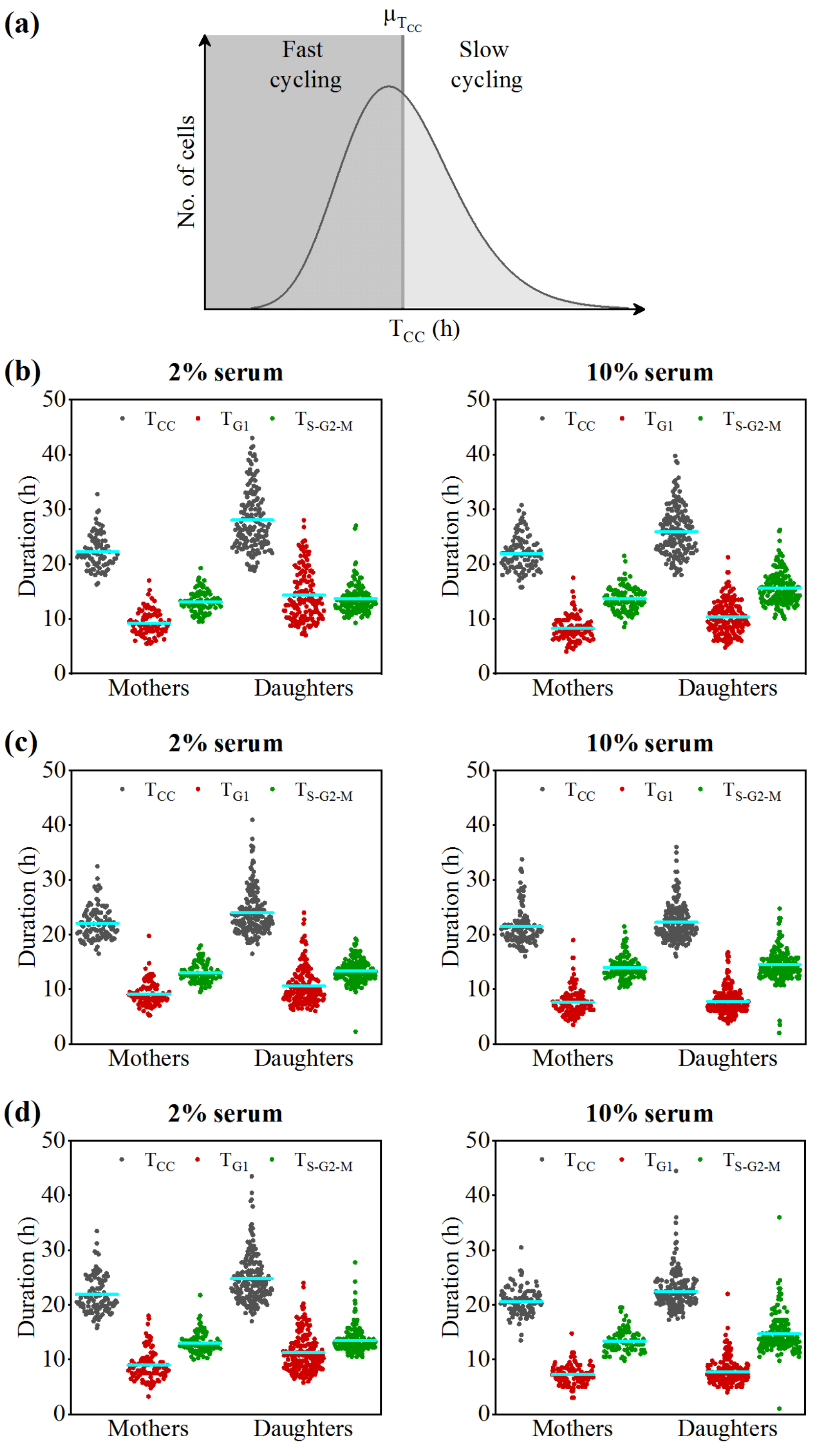


**Fig. S2.** (a) Schematic depiction for classifying cell population as slow and fast cycling based on average cell cycle period. Cell cycle period and phase durations distribution for mother and daughter cells quantified the from live cell imaging data of FUCCI-HeLa cells at 2% serum and 10% serum for (b) replicate II, (c) replicate III and (d) replicate IV.

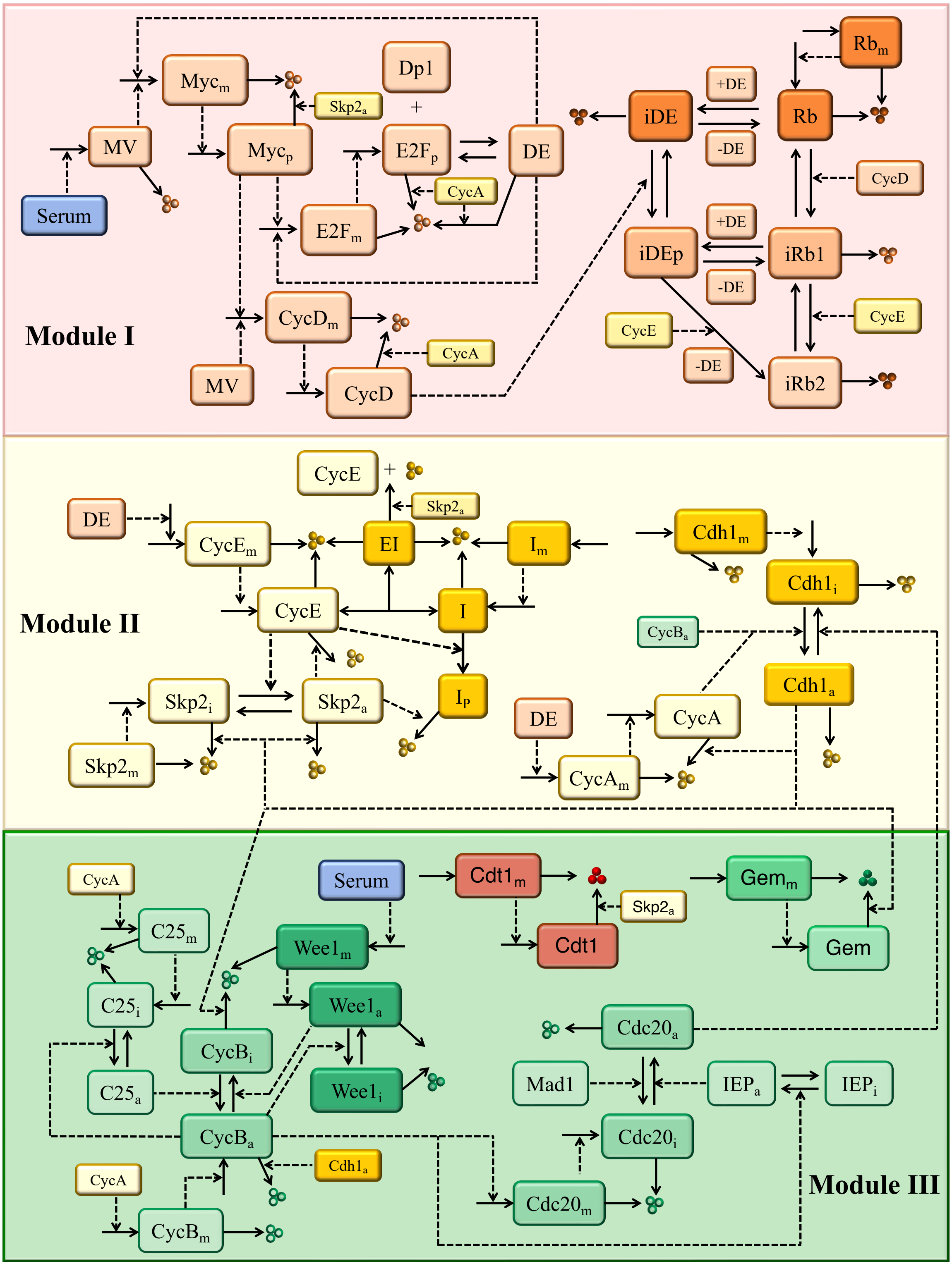


**Fig. S3. Detailed network interactions governing the different modules of cell cycle network proposed in Fig. 2(a).** Solid and dashed arrows represent biochemical reactions and catalytic activation processes respectively. The details of the regulatory interactions are described in Table S2 and SI Text. Corresponding kinetic equations and description of the parameters are described in Tables S3 and S4.

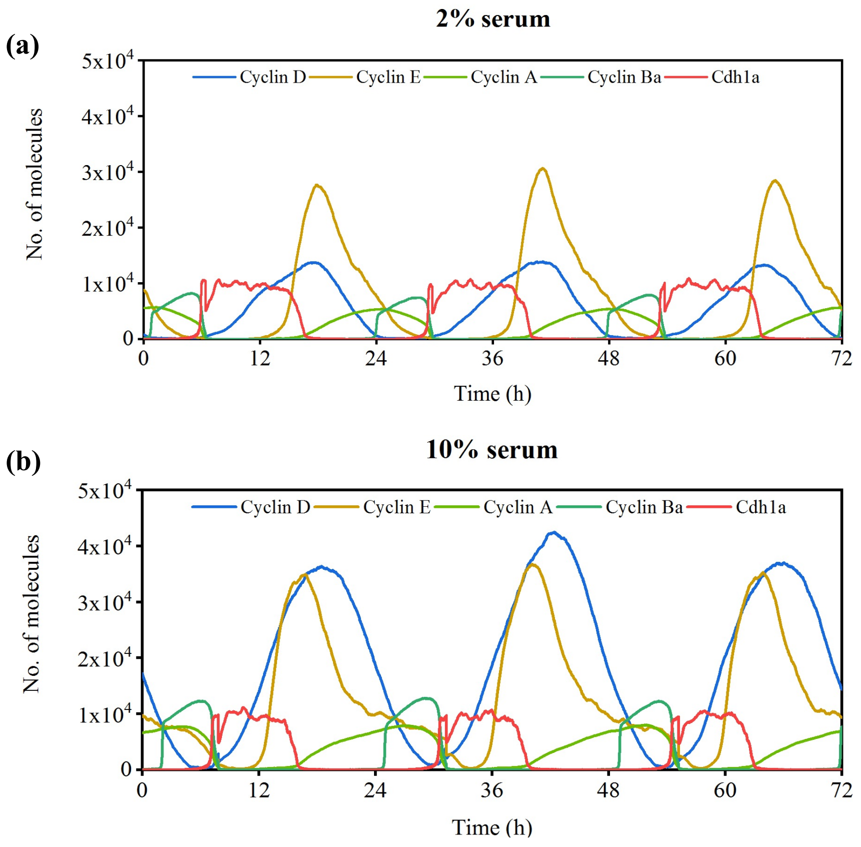


**Fig. S4.** Stochastic simulation time course trajectories for some of the important network components at (a) 2% serum and (b) 10% serum in the presence of only intrinsic noise (Fig. 2d(ii)).

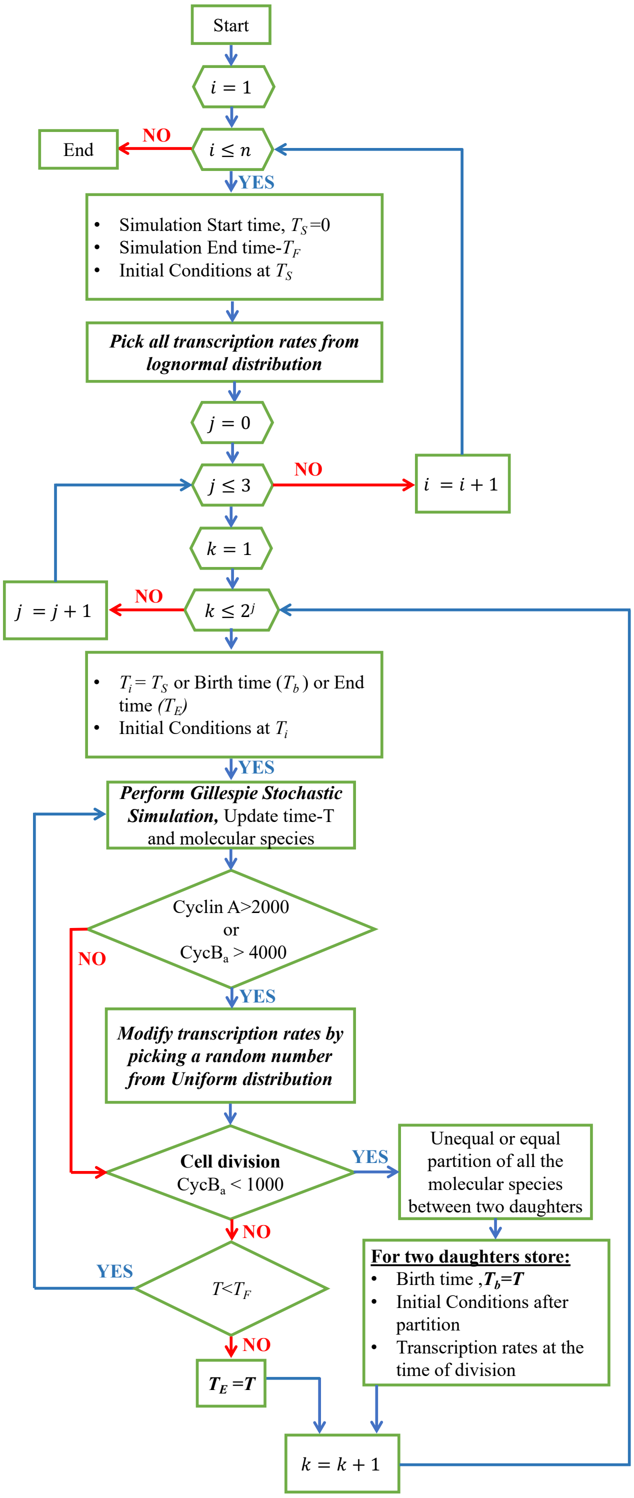


**Fig. S5. Stochastic simulation algorithm followed to implement the model simulation of the proposed network in cell lineages by incorporating various noise sources mentioned in Fig. 2(d).** Simulation for each starting mother cell ($0^{th}$ Generation, $j$ and $1^{st}$ Cell, $k$) in any cell lineage, $i$ starts with a start time, $T_S=0\ h$, end time, $T_E=72\ h$ and initial conditions for all the molecular species at $T_S$ obtained from deterministic simulation. Each transcription rate for the starting mother cell is chosen randomly from a lognormal distribution having corresponding deterministic rate as mean of the distribution. Gillespie stochastic simulation gets executed for the cell in which after each time step of the simulation, molecular species count gets updated for the occurred biochemical reaction. At each time step of the simulation, program also checks the criteria for transcription rate variation during cell cycle and for cell division. For transcription rate variation, a random number is picked from a uniform distribution between 0 to 1 and each rate is either decreased or increased x%, based on whether the picked random number for the rate is less than or greater than 0.5 respectively. Transcription rate variation occurs twice during simulation for any cell, when (a) Cyclin A level crosses 2000 as the level increases (indicating S phase entry) and (b) $CycB_a$ level crosses 4000 as the level increases (indicating M phase entry). In our simulation, cell division occurs when $CycB_a$ level crosses 1000 while it decreases, upon which all the molecular species are partitioned equally or unequally between two daughter cells. For unequal partition during cell division, a random number (r) is chosen from a normal distribution with mean 0.5 and 10% CV, based on which one of the daughter cells gets fraction ‘r’ and the other daughter receives (1-r) fraction of each molecular species respectively. The amount of each molecular species after partition, transcription rates of the mother at the time of division and division time stored in arrays with unique index for each cell. These information are needed for subsequent simulation of the daughter cells in the order in which they are added in the array in each generation of the lineage.

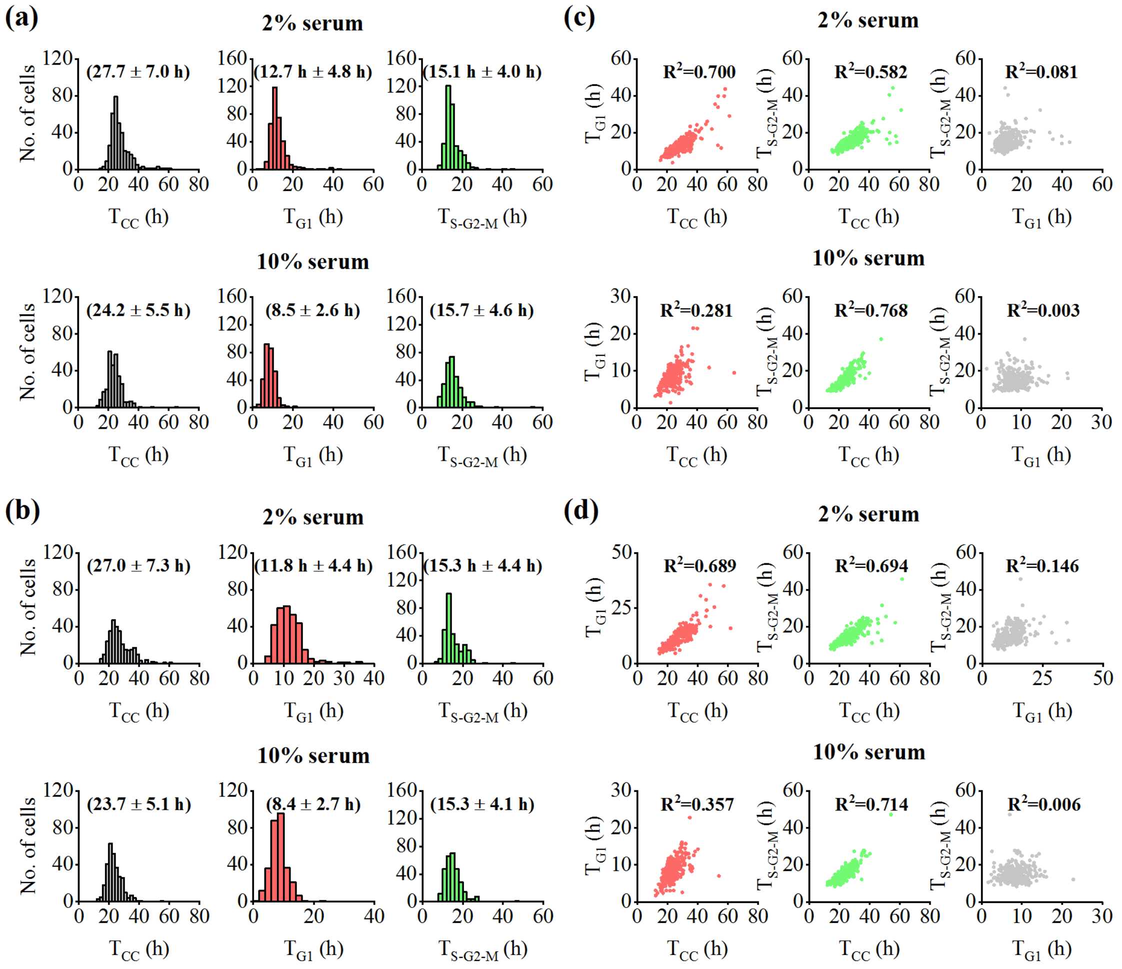


**Fig. S6.** Cell cycle, $G_1$ and S-$G_2$-M distribution quantified from stochastic simulation performed with Model M-1, Table-1 at 2% serum and 10% serum for (a) replicate II and (b) replicate III. Correlation pattern of cell cycle and phase timings observed from stochastic simulation performed with Model M-1, Table-1 at 2% serum and 10% serum for (c) replicate II and (d) replicate III.

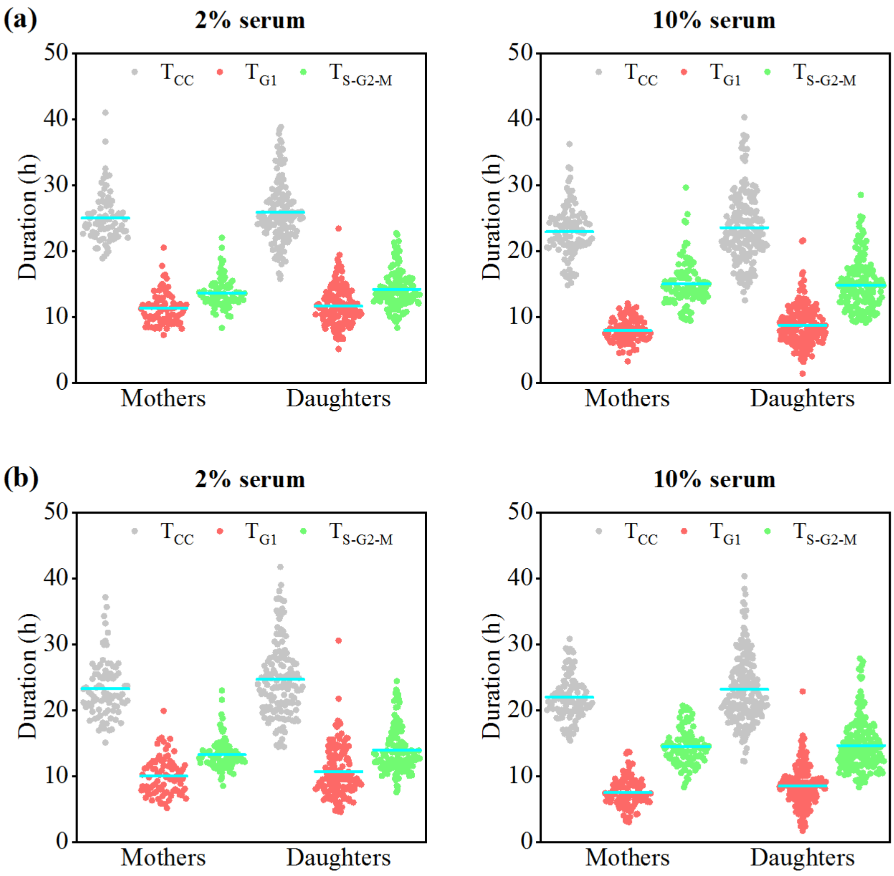


**Fig. S7.** Scatter plot showing the distribution of mothers and daughters from stochastic simulations performed with Model M-1, Table-1 at 2% serum and 10% serum for (a) replicate II and (b) replicate III.

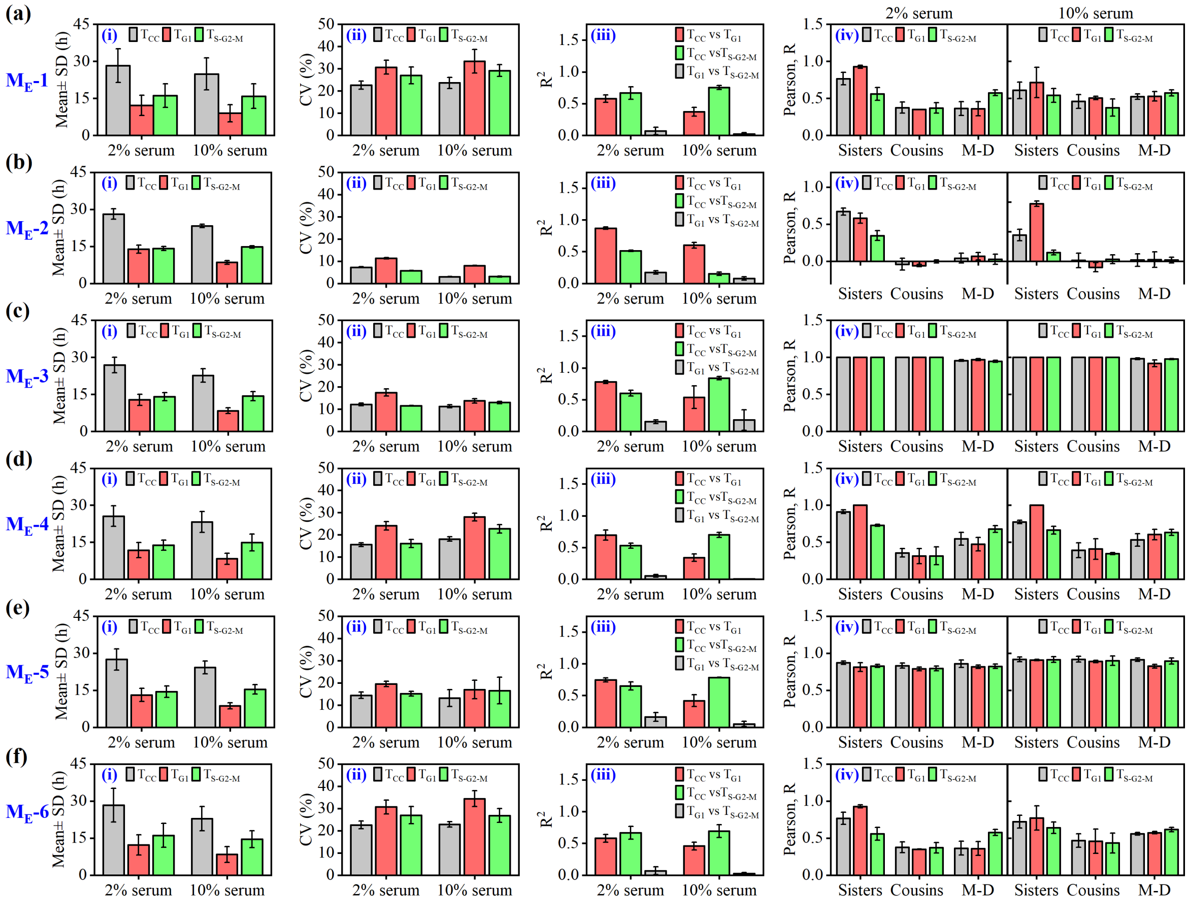


**Fig. S8. Simulation results at low (2%) and high (10%) serum concentrations from different model variants with equal division.** Calculated (i) Mean ± standard deviation (SD) for Replicate I, (ii) average coefficient of variation (CV), (iii) average correlation coefficient ($R^2$) of cell cycle period and phase durations, and (iv) average Pearson correlation coefficient (R) obtained for the cell cycle period and phase durations for sisters, cousins and mother-daughter pairs selected from different cell lineages (from three replicate simulations) for the respective model variants **(a)** M-1, **(b)** M-2, **(c)** M-3, **(d)** M-4, **(e)** M-5 and **(f)** M-6 defined in **Table-1** with equal division.

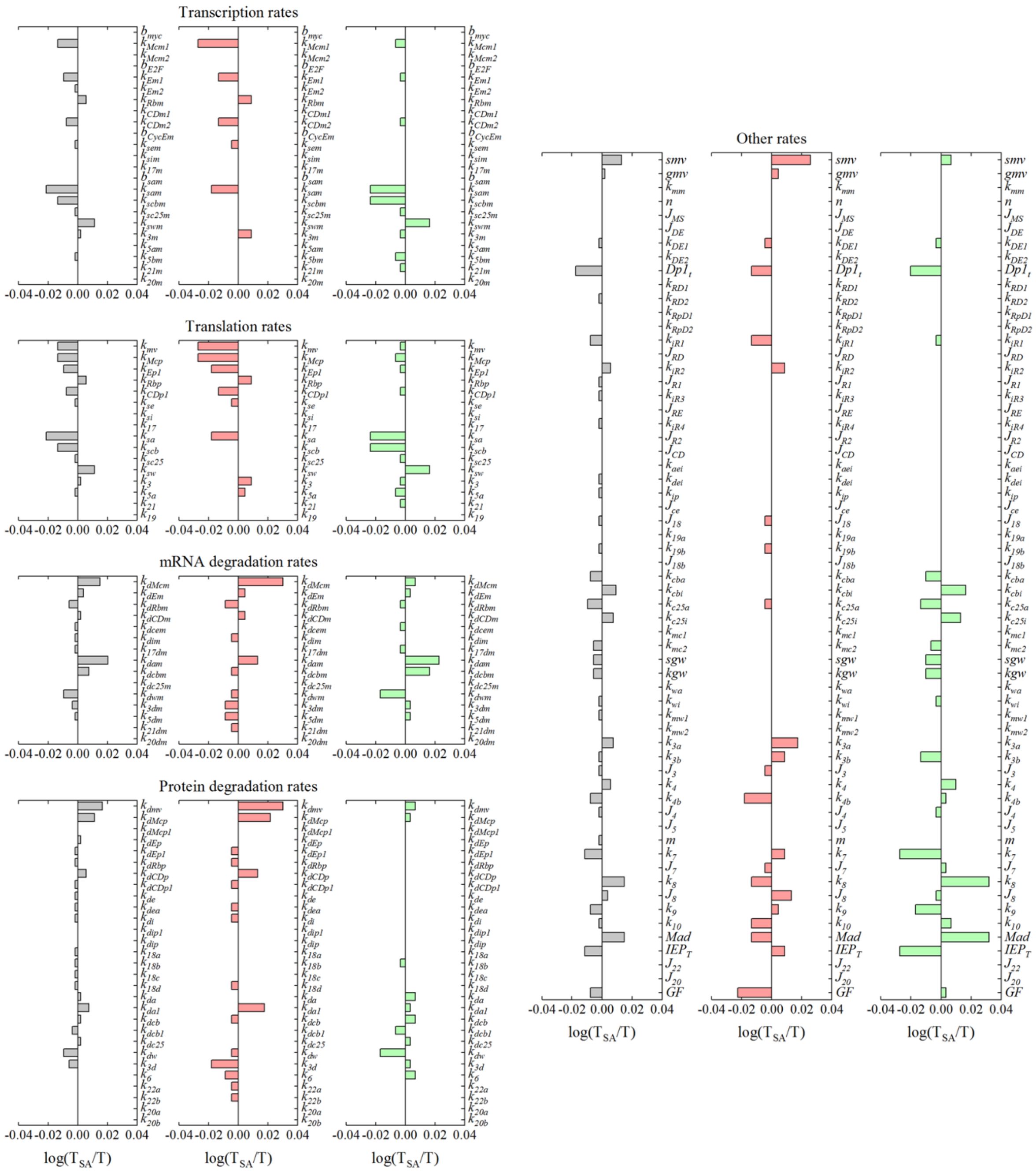


**Fig. S9.** Sensitivity analysis of the model parameters by taking cell cycle time period (Grey bar), $G_1$ duration (Red bar) and S-$G_2$-M duration (Green bar) as sensitivity criteria. Parameters were increased by 20% to analyse the sensitivity of the respective parameters.

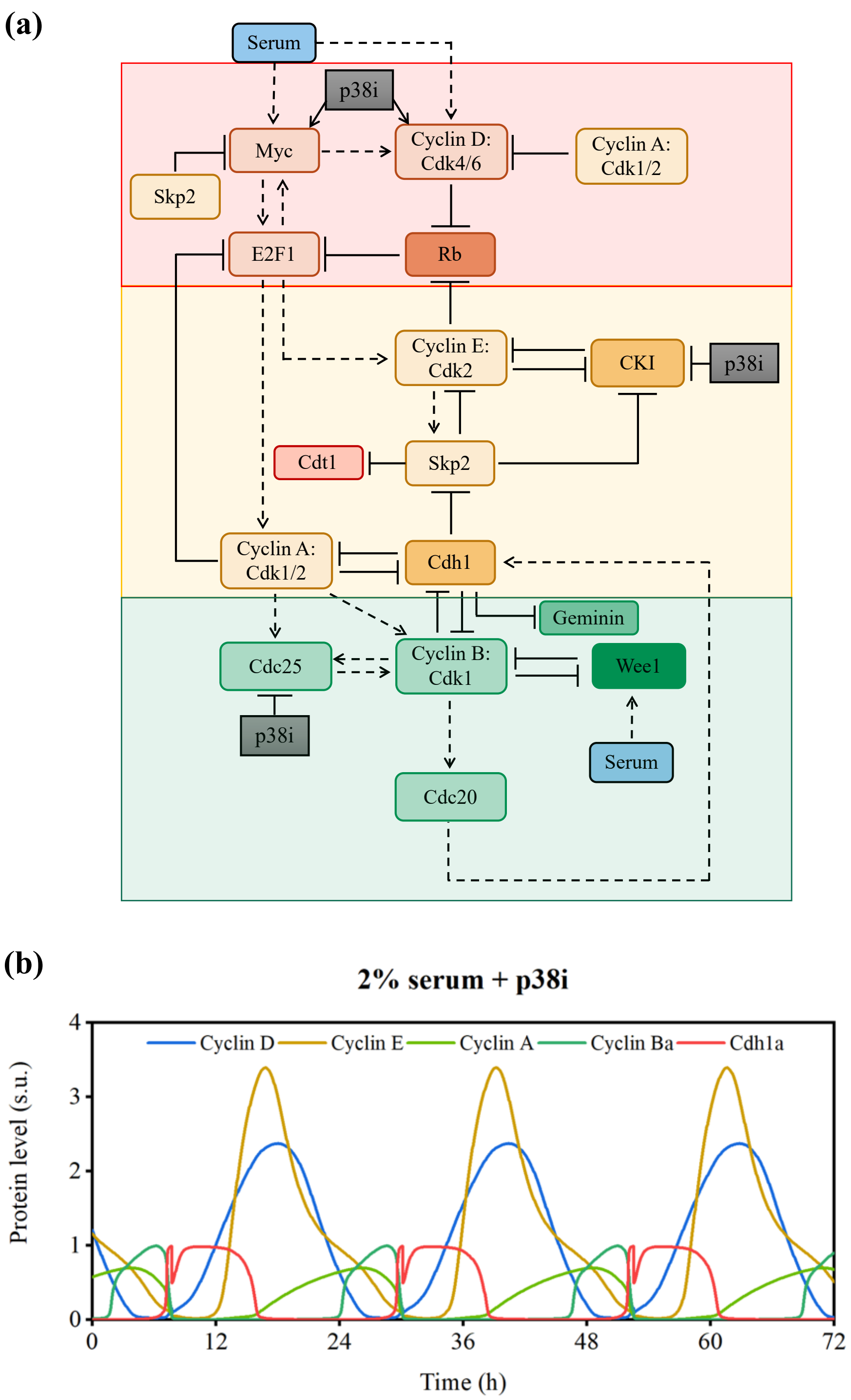


**Fig. S10. (a)** Effect of p38 protein in cell cycle regulation.(solid and dotted arrows represent biochemical reactions and catalytic processes respectively, while hammer head arrows represent inhibition processes).Detailed mechanism for the regulation of cell cycle network components by p38 is provided in **supplementary Text** and the corresponding differential equations are provided in **Table S10**. **(b)** Deterministic simulation time course trajectories for some of the network components at 2% serum in the presence of the effect of p38i as described in Table S10.

### Supplementary tables

**Table S1.** Table showing the data from the live cell imaging of FUCCI-HeLa cells grown at 2% and 10% serum concentration.

| Serum(%) | | $T_{CC}$ Mean (h) | $T_{CC}$ SD (h) | $T_{CC}$ CV (%) | $T_{G1}$ Mean (h) | $T_{G1}$ SD (h) | $T_{G1}$ CV (%) | $T_{S\text{-}G2\text{-}M}$ Mean (h) | $T_{S\text{-}G2\text{-}M}$ SD (h) | $T_{S\text{-}G2\text{-}M}$ CV (%) | $T_{CC}$ vs $T_{G1}$ $R^2$ | $T_{CC}$ vs $T_{S\text{-}G2\text{-}M}$ $R^2$ | $T_{G1}$ vs $T_{S\text{-}G2\text{-}M}$ $R^2$ | n |
|---|---|---|---|---|---|---|---|---|---|---|---|---|---|---|
| 2 | I | 26.8 | 5.3 | 19.8 | 12.0 | 4.1 | 34.2 | 14.8 | 3.1 | 20.9 | 0.656 | 0.419 | 0.006 | 300 |
| | II | 27.0 | 6.5 | 24.1 | 13.0 | 5.2 | 40.0 | 14.0 | 3.5 | 25.0 | 0.712 | 0.375 | 0.009 | 300 |
| | III | 24.3 | 4.7 | 19.3 | 10.7 | 3.5 | 32.7 | 13.6 | 2.7 | 19.9 | 0.687 | 0.471 | 0.026 | 315 |
| | IV | 24.4 | 4.5 | 18.4 | 10.8 | 3.4 | 31.5 | 13.6 | 2.5 | 18.2 | 0.689 | 0.452 | 0.021 | 320 |
| | avg | 25.6 | 5.3 | 20.4 | 11.6 | 4.1 | 34.6 | 14.0 | 3.0 | 21.0 | 0.686 | 0.429 | 0.016 | |
| 10 | I | 23.8 | 4.5 | 18.9 | 9.7 | 2.8 | 28.9 | 14.2 | 3.4 | 23.9 | 0.447 | 0.611 | 0.003 | 300 |
| | II | 25.1 | 4.7 | 18.7 | 9.8 | 3.0 | 30.6 | 15.3 | 3.3 | 21.6 | 0.516 | 0.599 | 0.013 | 300 |
| | III | 22.4 | 3.6 | 16.0 | 7.9 | 2.5 | 31.1 | 14.5 | 2.8 | 19.1 | 0.403 | 0.533 | 0.004 | 313 |
| | IV | 22.4 | 3.8 | 17.2 | 7.8 | 2.3 | 29.0 | 14.6 | 3.4 | 23.4 | 0.221 | 0.661 | 0.017 | 293 |
| | avg | 23.4 | 4.2 | 17.7 | 8.8 | 2.7 | 29.9 | 14.7 | 3.2 | 22.0 | 0.397 | 0.601 | 0.009 | |

| Serum(%) | | Sister pairs (Pearson, R) $T_{CC}$ | Sister $T_{G1}$ | Sister $T_{S\text{-}G2\text{-}M}$ | Sister n | Cousin pairs (Pearson, R) $T_{CC}$ | Cousin $T_{G1}$ | Cousin $T_{S\text{-}G2\text{-}M}$ | Cousin n | Mother-daughter pairs (Pearson, R) $T_{CC}$ | Mother-daughter $T_{G1}$ | Mother-daughter $T_{S\text{-}G2\text{-}M}$ | Mother-daughter n |
|---|---|---|---|---|---|---|---|---|---|---|---|---|---|
| 2 | I | 0.5873 | 0.71 | 0.4555 | 137 | 0.2707 | 0.2609 | 0.3724 | 89 | 0.2896 | 0.2784 | 0.18 | 131 |
| | II | 0.7453 | 0.8322 | 0.584 | 134 | 0.1619 | 0.3294 | 0.2962 | 121 | 0.0466 | 0.2824 | 0.0978 | 155 |
| | III | 0.7553 | 0.8086 | 0.5426 | 150 | 0.5374 | 0.5241 | 0.3179 | 166 | 0.3752 | 0.2896 | 0.5021 | 196 |
| | IV | 0.7442 | 0.7704 | 0.5585 | 150 | 0.5525 | 0.481 | 0.5473 | 187 | 0.247 | 0.326 | 0.3147 | 202 |
| | avg | 0.708 | 0.7803 | 0.5352 | | 0.3806 | 0.3989 | 0.3835 | | 0.2396 | 0.2941 | 0.2737 | |
| 10 | I | 0.7121 | 0.7952 | 0.6765 | 141 | 0.3975 | 0.3992 | 0.5077 | 142 | 0.3677 | 0.3089 | 0.4199 | 179 |
| | II | 0.729 | 0.786 | 0.5709 | 134 | 0.4723 | 0.5704 | 0.3851 | 144 | 0.2698 | 0.4657 | 0.3194 | 178 |
| | III | 0.7136 | 0.809 | 0.6268 | 145 | 0.4345 | 0.5634 | 0.5545 | 177 | 0.334 | 0.1791 | 0.3936 | 214 |
| | IV | 0.5471 | 0.8046 | 0.4594 | 135 | 0.2122 | 0.3419 | 0.3691 | 178 | 0.104 | 0.048 | 0.3745 | 201 |
| | avg | 0.6755 | 0.7987 | 0.5834 | | 0.3791 | 0.4687 | 0.4541 | | 0.2689 | 0.2504 | 0.3769 | |

**Table S2.** Reaction governing the proposed cell cycle network in Fig. 2(a):

| **S.No** | **Reaction** | **Kinetics** | **Description** |
|---|---|---|---|
| **MV variable** | | | |
| R1 | $\xrightarrow{Serum} MV$ | $\frac{k_{mV} \times s \times GF}{smv + GF}$ | Serum mediated synthesis of MV |
| R2 | $MV \rightarrow ::$ | $k_{dmv} \times MV$ | Degradation of MV |
| **Myc mRNA** | | | |
| R3 | $\rightarrow Myc_m$ | $b_{myc} \times s$ | Basal synthesis of Myc mRNA |
| R4 | $\xrightarrow{MV} Myc_m$ | $k_{Mcm1} \times MV$ | MV mediated synthesis of Myc mRNA |
| R5 | $\xrightarrow{DE} Myc_m$ | $\frac{k_{Mcm2} \times s \times DE^n}{(k_{mm}^n \times s^n) + DE^n}$ | DE mediated synthesis of Myc mRNA |
| R6 | $Myc_m \rightarrow ::$ | $k_{dMcm} \times Myc_m$ | Degradation of Myc mRNA |
| **Myc protein** | | | |
| R7 | $\xrightarrow{Myc_m} Myc_p$ | $k_{Mcp} \times Myc_m$ | Synthesis of Myc protein |
| R8 | $Myc_p \rightarrow ::$ | $k_{dMcp} \times Myc_p$ | Degradation of Myc protein |
| R9 | $Myc_p \xrightarrow{Skp2_a} ::$ | $\frac{k_{dMcp1} \times Myc_p \times Skp2}{(s \times J_{ms}) + Myc_p}$ | Skp2 mediated degradation of Myc protein |
| **E2F mRNA** | | | |
| R10 | $\rightarrow E2F_m$ | $b_{e2f} \times s$ | Basal synthesis of E2F mRNA |
| R11 | $\xrightarrow{Myc_p} E2F_m$ | $k_{Em1} \times Myc_p$ | Myc mediated synthesis of E2F mRNA |
| R12 | $\xrightarrow{DE} E2F_m$ | $\frac{k_{Em2} \times s \times DE^n}{(k_{mm}^n \times s^n) + DE^n}$ | DE mediated synthesis of E2F mRNA |
| R13 | $E2F_m \rightarrow ::$ | $k_{dEm} \times E2F_m$ | Degradation of E2F mRNA |
| **E2F protein** | | | |
| R14 | $\xrightarrow{E2F_m} E2F_p$ | $k_{Ep1} \times E2F_m$ | Synthesis of E2F protein |
| R15 | $E2F_p \rightarrow ::$ | $k_{dEp} \times E2F_p$ | Degradation of E2F protein |
| R16 | $E2F_p \xrightarrow{CycA} ::$ | $\frac{k_{dEp1} \times CycA \times E2F_p}{(s \times J_{DE}) + E2F_p}$ | CycA mediated degradation of E2F protein |
| **DE complex** | | | |
| R17 | $E2F_p + Dp1_p \rightarrow DE$ | $\frac{k_{DE1}}{s} \times Dp1_p \times E2F_p$ | Complex formation of Dp1 and E2F protein |
| R18 | $DE \rightarrow E2F_p + Dp1_p$ | $k_{DE2} \times DE$ | Dissociation of DE complex into Dp1 and E2F protein |
| R19 | $DE \rightarrow ::$ | $k_{dEp} \times DE$ | Degradation of DE protein |
| R20 | $DE \xrightarrow{CycA} ::$ | $\frac{k_{dEp1} \times CycA \times DE}{(s \times J_{DE}) + DE}$ | CycA mediated degradation of DE protein |

| | | | |
|---|---|---|---|
| **iDE complex** | | | |
| R21 | $DE + Rb \rightarrow iDE$ | $\frac{k_{RD1}}{s} \times DE \times Rb$ | Complex formation of Rb and DE protein |
| R22 | $iDE \rightarrow DE + Rb$ | $k_{RD2} \times iDE$ | Dissociation of inactive DE complex into Rb and E2F protein |
| R23 | $iDE \rightarrow ::$ | $k_{dEp} \times iDE$ | Degradation of inactive DE protein |
| **iDEP complex** | | | |
| R24 | $DE + iRb1 \rightarrow iDEP$ | $\frac{k_{RpD1}}{s} \times DE \times iRb1$ | Complex formation of single phosphorylated Rb and DE complex |
| R25 | $iDEP \rightarrow DE + iRb1$ | $k_{RpD2} \times iDEP$ | Dissociation of iDEP complex into single phosphorylated Rb and DE complex |
| R26 | $iDE \xrightarrow{CycD} iDEP$ | $\frac{k_{iR1} \times iDE \times CycD}{(J_{RD} \times s) + iDE}$ | First phosphorylation of Rb in iDE by cyclin D |
| R27 | $iDEP \longrightarrow iDE$ | $\frac{k_{iR2} \times s \times iDEP}{(J_{R1} \times s) + iDEP}$ | Dephosphorylation of single phosphorylated Rb in iDEP |
| R28 | $iDEP \xrightarrow{CycE} iRb2 + DE$ | $\frac{k_{iR3} \times iDEP \times CycE}{(J_{RE} \times s) + iDEP}$ | Second phosphorylation of Rb in iDEP by cyclin E |
| **Rb mRNA** | | | |
| R29 | $\rightarrow Rb_m$ | $k_{Rbm} \times s$ | Synthesis of Rb mRNA |
| R30 | $Rb_m \rightarrow ::$ | $k_{dRbm} \times Rb_m$ | Degradation of Rb mRNA |
| **Rb protein** | | | |
| R31 | $\xrightarrow{Rb_m} Rb$ | $k_{Rbp} \times Rb_m$ | Synthesis of Rb protein |
| R32 | $Rb \rightarrow ::$ | $k_{dRbp} \times Rb$ | Degradation of Rb protein |
| R33 | $Rb \xrightarrow{CycD} iRb1$ | $\frac{k_{iR1} \times Rb \times CycD}{(J_{RD} \times s) + Rb}$ | First phosphorylation of Rb by cyclin D |
| R34 | $iRb1 \longrightarrow Rb$ | $\frac{k_{iR2} \times s \times iRb1}{(J_{R1} \times s) + iRb1}$ | Dephosphorylation of single phosphorylated Rb |
| R35 | $iRb1 \xrightarrow{CycE} iRb2$ | $\frac{k_{iR3} \times iRb1 \times CycE}{(J_{RE} \times s) + iRb1}$ | Second phosphorylation of Rb by cyclin E |
| R36 | $iRb2 \longrightarrow iRb1$ | $\frac{k_{iR4} \times s \times iRb2}{(J_{R2} \times s) + iRb2}$ | Dephosphorylation of double phosphorylated Rb |
| R37 | $iRb1 \rightarrow ::$ | $k_{dRbp} \times iRb1$ | Degradation of single phosphorylated Rb |
| R38 | $iRb2 \rightarrow ::$ | $k_{dRbp} \times iRb2$ | Degradation of double phosphorylated Rb |

| | | | |
|---|---|---|---|
| **Cyclin D mRNA** | | | |
| R39 | $\xrightarrow{MV} CycD_m$ | $k_{CDm1} \times MV$ | MV mediated synthesis of Cyclin D mRNA |
| R40 | $\xrightarrow{Myc_p} CycD_m$ | $k_{CDm2} \times Myc_p$ | Myc mediated synthesis of Cyclin D mRNA |
| R41 | $CycD_m \rightarrow ::$ | $k_{dCDm} \times CycD_m$ | Degradation of Cyclin D mRNA |
| **Cyclin D** | | | |
| R42 | $\xrightarrow{CycD_m} CycD$ | $k_{CDp1} \times CycD_m$ | Synthesis of Cyclin D protein |
| R43 | $CycD \rightarrow ::$ | $k_{dCDp} \times CycD$ | Degradation of Cyclin D protein |
| R44 | $CycD \xrightarrow{CycA} ::$ | $\frac{k_{dCDp1} \times CycD \times CycA}{(s \times J_{CD}) + CycD}$ | Cyclin A mediated degradation of Cyclin D protein |
| **Cyclin E mRNA** | | | |
| R45 | $\rightarrow CycE_m$ | $b_{Cycem} \times s$ | Synthesis of Cyclin E mRNA |
| R46 | $\xrightarrow{DE} CycE_m$ | $k_{sem} \times DE$ | E2F mediated synthesis of Cyclin E mRNA |
| R47 | $CycE_m \rightarrow ::$ | $k_{dCEm} \times CycE_m$ | Degradation of Cyclin E mRNA |
| **Cyclin E** | | | |
| R48 | $\xrightarrow{CycE_m} CycE$ | $k_{se} \times CycE_m$ | Synthesis of Cyclin E protein |
| R49 | $CycE \rightarrow ::$ | $k_{dE} \times CycE$ | Degradation of Cyclin E protein |
| R50 | $CycE \xrightarrow{Skp2_a} ::$ | $\frac{k_{dEA}}{s} \times CycE \times Skp2_a$ | Skp2 mediated degradation of Cyclin E protein |
| **CKI mRNA** | | | |
| R51 | $\rightarrow I_m$ | $k_{sim} \times s$ | Synthesis of CKI mRNA |
| R52 | $I_m \rightarrow ::$ | $k_{dim} \times I_m$ | Degradation of CKI mRNA |
| **CKI protein** | | | |
| R53 | $\xrightarrow{I_m} I$ | $k_{si} \times I_m$ | Synthesis of CKI protein |
| R54 | $I \rightarrow ::$ | $k_{di} \times I$ | Degradation of CKI protein |
| **Cyclin E-CKI complex** | | | |
| R55 | $CycE + I \rightarrow EI$ | $\frac{k_{aei}}{s} \times CycE \times I$ | Complex formation of Cyclin E and CKI |
| R56 | $EI \rightarrow CycE + I$ | $k_{dei} \times EI$ | Dissociation of Cyclin E: CKI complex |
| R57 | $EI \rightarrow ::$ | $k_{di} \times EI$ | Degradation of Cyclin E: CKI complex |

| | | | |
|---|---|---|---|
| R58 | $EI \rightarrow ::$ | $k_{dE} \times EI$ | Degradation of Cyclin E: CKI complex |
| R59 | $EI \xrightarrow{Skp2_a} CycE+::$ | $\frac{k_{dip}}{s} \times EI \times Skp2_a$ | Skp2 mediated degradation of I in EI complex to give free CycE |
| **CKI Phosphorylated form** | | | |
| R60 | $I \xrightarrow{CycE} I_p$ | $\frac{k_{ip} \times CycE \times I}{(J_{CE} \times s) + I}$ | Cyclin E mediated phosphorylation of CKI |
| R61 | $I_p \rightarrow ::$ | $k_{dip1} \times I_p$ | Degradation of phosphorylated CKI |
| R62 | $I_p \xrightarrow{Skp2_a} ::$ | $\frac{k_{dip}}{s} \times I_p \times Skp2_a$ | SKp2 mediated degradation of phosphorylated CKI |
| **Skp2 mRNA** | | | |
| R63 | $\rightarrow Skp2_m$ | $k_{17m} \times s$ | Synthesis of Skp2 mRNA |
| R64 | $Skp2_m \rightarrow ::$ | $k_{17dm} \times Skp2_m$ | Degradation of Skp2 mRNA |
| **Skp2 protein** | | | |
| R65 | $\xrightarrow{Skp2_m} Skp2_i$ | $k_{17} \times Skp2_m$ | Synthesis of Skp2 protein |
| R66 | $Skp2_i \rightarrow ::$ | $k_{18a} \times (Skp2_T - Skp2_a)$ | Degradation of inactive form of Skp2 protein |
| R67 | $Skp2_i \xrightarrow{Cdh1_a} ::$ | $\frac{k_{18b} \times (Skp2_T - Skp2_a) \times Cdh1}{(J_{18} \times s) + Skp2_T - Skp2_a}$ | Cdh1 mediated degradation of inactive form of Skp2 protein |
| R68 | $Skp2_i \xrightarrow{CycE} Skp2_a$ | $\frac{k_{19a}}{s} \times CycE \times (Skp2_T - Skp2_a)$ | Cyclin E mediated phosphorylation of Skp2 |
| R69 | $Skp2_a \rightarrow Skp2_i$ | $k_{19b} \times Skp2_a$ | Dephosphorylation of phosphorylated Skp2 |
| R70 | $Skp2_a \rightarrow ::$ | $k_{18c} \times Skp2_a$ | Degradation of phosphorylated Skp2 protein |
| R71 | $Skp2_a \xrightarrow{Cdh1_a} ::$ | $\frac{k_{18d} \times Skp2_a \times Cdh1_a}{(J_{18b} \times s) + Skp2_a}$ | Cdh1 mediated degradation of phosphorylated Skp2 protein |
| **Cdh1 mRNA** | | | |
| R72 | $\rightarrow Cdh1_m$ | $k_{3m} \times s$ | Synthesis of Cdh1 mRNA |
| R73 | $Cdh1_m \rightarrow ::$ | $k_{3dm} \times Cdh1_m$ | Degradation of Cdh1 mRNA |
| **Cdh1 protein** | | | |
| R74 | $\xrightarrow{Cdh1_m} Cdh1_i$ | $k_3 \times Cdh1_m$ | Synthesis of Cdh1 protein |
| R75 | $Cdh1_i \rightarrow ::$ | $(k_{3d} \times Cdh1_T - Cdh1_a)$ | Degradation of inactive form of Cdh1 protein |
| R76 | $Cdh1_i \rightarrow Cdh1_a$ | $\frac{(k_{3a} \times s) \times (Cdh1_T - Cdh1_a)}{(J_3 \times s) + Cdh1_T - Cdh1_a}$ | Basal activation of Cdh1 protein |

| | | | |
|---|---|---|---|
| R77 | $Cdh1_i \xrightarrow{Cdc20_a} Cdh1_a$ | $\frac{k_{3b} \times Cdc20_A \times (Cdh1_T - Cdh1_{[illegible]}}{(J_3 \times s) + Cdh1_T - Cdh1_a}$ | Cdc20 mediated activation of Cdh1 protein |
| R78 | $Cdh1_a \xrightarrow{CycB_a} Cdh1_i$ | $\frac{k_4 \times CycB_a \times Cdh1_a}{(J_4 \times s) + Cdh1_a}$ | Cyclin A mediated inactivation of Cdh1 protein |
| R79 | $Cdh1_a \xrightarrow{CycA} Cdh1_i$ | $\frac{k_{4b} \times CycA \times Cdh1_a}{(J_4 \times s) + Cdh1_a}$ | Cyclin E mediated inactivation of Cdh1 protein |
| R80 | $Cdh1_a \rightarrow ::$ | $k_{3d} \times Cdh1_a$ | Degradation of inactive form of Cdh1 protein |
| **Cyclin A mRNA** | | | |
| R81 | $\rightarrow CycA_m$ | $b_{sam}$ | Basal synthesis rate of Cyclin A mRNA |
| R82 | $\xrightarrow{DE} CycA_m$ | $k_{sam} \times DE$ | E2F mediated synthesis of Cyclin A mRNA |
| R83 | $CycA_m \rightarrow ::$ | $k_{dam} \times CycA_m$ | Degradation of Cyclin A mRNA |
| **Cyclin A protein** | | | |
| R84 | $\xrightarrow{CycA_m} CycA$ | $k_{sa} \times CycA_m$ | Synthesis of Cyclin A protein |
| R85 | $CycA \rightarrow ::$ | $k_{da} \times CycA$ | Degradation of Cyclin A protein |
| R86 | $CycA \xrightarrow{Cdh1_a} ::$ | $k_{da1} \times CycA \times Cdh1_a$ | Cdh1 mediated Cyclin A protein |
| **Cyclin B mRNA** | | | |
| R87 | $\xrightarrow{CycA} CycB_m$ | $\frac{k_{scbm}}{s} \times CycA$ | Cyclin A mediated synthesis of Cyclin B mRNA |
| R88 | $CycB_m \rightarrow ::$ | $k_{dcbm} \times CycB_m$ | Degradation of Cyclin B mRNA |
| **Cyclin B protein** | | | |
| R89 | $\xrightarrow{CycB_m} CycB_a$ | $k_{scb} \times CycB_m$ | Synthesis of Cyclin B protein |
| R90 | $CycB_a \rightarrow ::$ | $k_{dcb} \times CycB_a$ | Degradation of Cyclin B protein |
| R91 | $CycB_a \xrightarrow{Cdh1_a} ::$ | $\frac{k_{dcb1}}{s} \times Cdh1 \times CycB_a$ | Cdh1 mediated Cyclin B protein |
| R92 | $CycB_a \xrightarrow{Wee1_a} CycB_i$ | $\frac{k_{cbi}}{s} \times Wee1_a \times CycB_a$ | Wee1 mediated dephosphorylation of Cyclin B protein |
| R93 | $CycB_a \xrightarrow{Cdc25_a} CycB_i$ | $\frac{k_{cba}}{s} \times C25_a \times CycB_i$ | Cdc25 mediated phosphorylation of Cyclin B protein |

| | | | |
|---|---|---|---|
| R94 | $CycB_i \rightarrow ::$ | $k_{dcb} \times CycB_i$ | Degradation of inactive form of Cyclin B protein |
| R95 | $CycB_i \xrightarrow{Cdh1_a} ::$ | $\frac{k_{dcb1}}{s} \times Cdh1 \times CycB_i$ | Degradation of active form of Cyclin B protein |
| **Wee1 mRNA** | | | |
| R96 | $\xrightarrow{Serum} Wee1_m$ | $\frac{k_{swm} \times s \times Serum}{sgw + (Serum * kgw)}$ | Synthesis of Wee1 mRNA |
| R97 | $Wee1_m \rightarrow ::$ | $k_{dwm} \times Wee1_m$ | Degradation of Wee1 mRNA |
| **Wee1 protein** | | | |
| R98 | $\xrightarrow{Wee1_m} Wee1_a$ | $k_{sw} \times Wee1_m$ | Serum mediated synthesis of Wee1 protein |
| R99 | $Wee1_a \rightarrow ::$ | $k_{dw} \times Wee1_a$ | Degradation of Wee1 protein |
| R100 | $Wee1_a \xrightarrow{CycB_a} Wee1_i$ | $\frac{k_{wi} \times Wee1_a \times CycB_a}{(k_{mw2} \times s) + Wee1_a}$ | Cyclin B mediated phosphorylation of Wee1 protein |
| R101 | $Wee1_i \rightarrow Wee1_a$ | $\frac{k_{wa} \times s \times Wee1_i}{(k_{mc1} \times s) + Wee1_i}$ | Dephosphorylation of Wee1 protein |
| R102 | $Wee1_i \rightarrow ::$ | $k_{dw} \times Wee1_i$ | Degradation of inactive form of Wee1 protein |
| **Cdc25 mRNA** | | | |
| R103 | $\xrightarrow{CycA} Cdc25_m$ | $k_{sc25m} \times CycA$ | Cyclin A mediated synthesis of Cdc25 mRNA |
| R104 | $Cdc25_m \rightarrow ::$ | $k_{dc25m} \times C25_m$ | Degradation of Cdc25 mRNA |
| **Cdc25 protein** | | | |
| R105 | $\xrightarrow{Cdc25_m} Cdc25_i$ | $k_{sc25} \times C25_m$ | Synthesis of Cdc25 protein |
| R106 | $Cdc25_i \rightarrow ::$ | $k_{dc25} \times C25_i$ | Degradation of Cdc25 protein |
| R107 | $Cdc25_i \xrightarrow{CycB_a} Cdc25_a$ | $\frac{k_{c25a} \times C25_i \times CycB_a}{(k_{mc1} \times s) + C25_i}$ | Cyclin B mediated phosphorylation of Cdc25 protein |
| R108 | $Cdc25_a \rightarrow Cdc25_i$ | $\frac{k_{c25i} \times s \times C25_a}{(k_{mc2} \times s) + C25_a}$ | Dephosphorylation of Cdc25 protein |
| R109 | $Cdc25_a \rightarrow ::$ | $k_{dc25} \times C25_a$ | Degradation of inactive form of Cdc25 protein |
| **Cdc20 mRNA protein** | | | |
| R110 | $\rightarrow Cdc20_m$ | $k_{5am} \times s$ | Synthesis of Cdc20 mRNA |
| R111 | $\xrightarrow{CycB_a} Cdc20_m$ | $\frac{k_{5bm} \times s \times \left(\frac{CycB_a}{J_5 \times s}\right)^m}{1 + \left(\frac{CycB_a}{(J_5 \times s)}\right)^m}$ | MV and Cyclin A mediated synthesis of Cdc20 mRNA |
| R112 | $Cdc20_m \rightarrow ::$ | $k_{5dm} \times Cdc20_m$ | Degradation of Cdc20 mRNA |

| | | | |
|---|---|---|---|
| **Cdc20 protein** | | | |
| R113 | $\xrightarrow{Cdc20_m} Cdc20_i$ | $k_{5a} \times Cdc20_m$ | Synthesis of Cdc20 protein |
| R114 | $Cdc20_i \rightarrow ::$ | $k_6 \times (Cdc20_T - Cdc20_a)$ | Degradation of inactive Cdc20 protein |
| R115 | $Cdc20_i \xrightarrow{IEP_a} Cdc20_a$ | $\frac{k_7 \times IEP_a \times (Cdc20_T - Cdc20_a)}{(J_7 \times s) + Cdc20_T - Cdc20_a}$ | IEP mediated activation of Cdc20 protein |
| R116 | $Cdc20_a \xrightarrow{Mad} Cdh1_i$ | $\frac{k_8 \times Mad \times s \times Cdc20_a}{(J_8 \times s) + Cdc20_a}$ | Mad mediated inactivation of Cdc20 protein |
| R117 | $Cdc20_a \rightarrow ::$ | $k_6 \times Cdc20_a$ | Degradation of active Cdc20 protein |
| **IEP protein** | | | |
| R118 | $IEP_i \xrightarrow{CycB_a} IEP_a$ | $\frac{k_9}{s} \times CycB_a \times ((IEP_T \times s) - IEP_a)$ | Cyclin A mediated Phosphorylation of IEP protein |
| R119 | $IEP_a \rightarrow IEP_i$ | $k_{10} \times IEP_a$ | Dephosphorylation of phosphorylated IEP protein |
| **Cdt1** | | | |
| R120 | $\rightarrow Cdt1_m$ | $k_{21m} \times s$ | Synthesis of Cdt1 mRNA |
| R121 | $Cdt1_m \rightarrow ::$ | $k_{21dm} \times Cdt1_m$ | Degradation of Cdt1 mRNA |
| R122 | $\xrightarrow{Cdt1_m} Cdt1$ | $k_{21} \times Cdt1_m$ | Synthesis of Cdt1 protein |
| R123 | $Cdt1 \rightarrow ::$ | $k_{22a} \times Cdt1$ | Degradation of Cdt1 protein |
| R124 | $Cdt1 \xrightarrow{Skp2_a} ::$ | $\frac{k_{22b} \times Cdt1 \times Skp2_a}{(J_{22} \times s) + Cdt1}$ | Skp2 mediated degradation of Cdt1 protein |
| **Geminin** | | | |
| R125 | $\rightarrow Gem_m$ | $k_{20m} \times s$ | Synthesis of Geminin mRNA |
| R126 | $Gem_m \rightarrow ::$ | $k_{20dm} \times Gem_m$ | Degradation of Geminin mRNA |
| R127 | $\xrightarrow{Gem_m} Gem$ | $k_{19} \times Gem_m$ | Synthesis of Geminin protein |
| R128 | $Gem \rightarrow ::$ | $k_{20a} \times Gem$ | Degradation of Geminin protein |
| R129 | $Gem \xrightarrow{Cdh1_a} ::$ | $\frac{k_{20b} \times Gem \times Cdh1_a}{(J_{20} \times s) + Gem}$ | Cdh1 mediated degradation of Geminin protein |

**Table S3**. Equations governing reactions in Table S2 for the proposed cell cycle network in Fig. 2(a)

| | |
|---|---|
| $\frac{dMV}{dt} = \frac{k_{mV} \times s \times Serum \times cf}{smv + (Serum * gmv)} - k_{dmv} \times MV$ | 1 |
| $\frac{dMyc_m}{dt} = (b_{myc} \times s) + k_{Mcm1} \times MV + \frac{k_{Mcm2} \times s \times DE^n}{(k_{mm}^n \times s^n) + DE^n} - k_{dMcm} \times Myc_m$ | 2 |
| $\frac{dMyc_p}{dt} = k_{Mcp} \times Myc_m - k_{dMcp} \times Myc_p - \frac{k_{dMcp1} \times Myc_p \times Skp2_a}{(s \times J_{ms}) + Myc_p}$ | 3 |
| $\frac{dE2F_m}{dt} = (b_{e2f} \times s) + k_{Em1} \times Myc_p + \frac{k_{Em2} \times s \times DE^n}{(k_{mm}^n \times s^n) + DE^n} - k_{dEm} \times E2F_m$ | 4 |
| $\frac{dE2F_T}{dt} = k_{Ep1} \times E2F_m - k_{dEp} \times E2F_p - k_{dEp} \times DE - k_{dEp} \times iDE - \frac{k_{dEp1} \times CycA \times E2F_p}{(s \times J_{DE}) + E2F_p} - \frac{k_{dEp1} \times CycA \times DE}{(s \times J_{DE}) + DE}$ | 5 |
| $\frac{dDE}{dt} = \frac{k_{DE1}}{s} \times Dp1_p \times E2F_p - k_{DE2} \times DE - \frac{k_{RD1}}{s} \times DE \times Rb + k_{RD2} \times iDE - k_{dEp} \times DE + \frac{k_{iR3} \times iDEP \times CycE}{(J_{RE} \times s) + iDEP} - \frac{k_{RpD1}}{s} \times DE \times iRb1 + k_{RpD2} \times iDEP - \frac{k_{dEp1} \times CycA \times DE}{(s \times J_{DE}) + DE}$ | 6 |
| $\frac{diDE}{dt} = \frac{k_{RD1}}{s} \times DE \times Rb - k_{RD2} \times iDE - k_{dEp} \times iDE - \frac{k_{iR1} \times iDE \times CycD}{(J_{RD} \times s) + iDE} + \frac{k_{iR2} \times s \times iDEP}{(J_{R1} \times s) + iDEP}$ | 7 |
| $\frac{diDEP}{dt} = \frac{k_{iR1} \times iDE \times CycD}{(J_{RD} \times s) + iDE} - \frac{k_{iR2} \times s \times iDEP}{(J_{R1} \times s) + iDEP} - \frac{k_{iR3} \times iDEP \times CycE}{(J_{RE} \times s) + iDEP} + \frac{k_{RpD1}}{s} \times DE \times iRb1 - k_{RpD2} \times iDEP$ | 8 |
| $\frac{dRb_m}{dt} = k_{Rbm} \times s - k_{dRbm} \times Rb_m$ | 9 |
| $\frac{dRb_T}{dt} = k_{Rbp} \times Rb_m - k_{dRbp} \times (Rb + iRb1 + iRb2) - k_{dEp} \times iDE$ | 10 |
| $\frac{diRb1}{dt} = \frac{k_{iR1} \times Rb \times CycD}{(J_{RD} \times s) + Rb} - \frac{k_{iR2} \times s \times iRb1}{(J_{R1} \times s) + iRb1} - \frac{k_{iR3} \times iRb1 \times CycE}{(J_{RE} \times s) + iRb1} + \frac{k_{iR4} \times s \times iRb2}{(J_{R2} \times s) + iRb2} - \frac{k_{RpD1}}{s} \times DE \times iRb1 + k_{RpD2} \times iDEP - k_{dRbp} \times iRb1$ | 11 |
| $\frac{diRb2}{dt} = \frac{k_{iR3} \times iRb1 \times CycE}{(J_{RE} \times s) + iRb1} + \frac{k_{iR4} \times s \times iRb2}{(J_{R2} \times s) + iRb2} + \frac{k_{iR3} \times iDEP \times CycE}{(J_{RE} \times s) + iDEP} - k_{dRbp} \times iRb2$ | 12 |
| $\frac{dCycD_m}{dt} = k_{CDm1} \times MV + k_{CDm2} \times Myc_p - k_{dCDm} \times CycD_m$ | 13 |
| $\frac{dCycD}{dt} = k_{CDp1} \times CycD_m - k_{dCDp} \times CycD - \frac{k_{dCDp1} \times CycD \times CycA}{(s \times J_{CD}) + CycD}$ | 14 |
| $\frac{dCycE_m}{dt} = (b_{Cycem} \times s) + k_{sem} \times DE - k_{dCEm} \times CycE_m$ | 15 |
| $\frac{dCycE}{dt} = k_{se} \times CycE_m - k_{dE} \times CycE - \frac{k_{dEA}}{s} \times CycE \times Skp2 - \frac{k_{aei}}{s} \times CycE \times I + k_{dei} \times EI + \frac{k_{dip}}{s} \times EI \times Skp2_a$ | 16 |
| $\frac{dI_m}{dt} = k_{sim} \times s - k_{dim} \times I_m$ | 17 |
| $\frac{dI}{dt} = k_{si} \times I_m - k_{di} \times I - \frac{k_{ip} \times CycE \times I}{(J_{CE} \times s) + I} - \frac{k_{aei}}{s} \times CycE \times I + k_{dei} \times EI$ | 18 |
| $\frac{dI_p}{dt} = \frac{k_{ip} \times CycE \times I}{(J_{CE} \times s) + I} - k_{dip1} \times I_p - \frac{k_{dip}}{s} \times I_p \times Skp2_a$ | 19 |

| | |
|---|---|
| $\frac{dEI}{dt} = \frac{k_{aei}}{s} \times CycE \times I - k_{dei} \times EI - k_{di} \times EI - k_{dE} \times EI - \frac{k_{dip}}{s} \times EI \times Skp2_a$ | 20 |
| $\frac{dSkp2_m}{dt} = k_{17m} \times s - k_{17dm} \times Skp2_m$ | 21 |
| $\frac{dSkp2_T}{dt} = k_{17} \times Skp2_m - k_{18a} \times (Skp2_T - Skp2_a) - \frac{k_{18b} \times (Skp2_T - Skp2_a) \times Cdh1_a}{(J_{18} \times s) + Skp2_T - Skp2_a} - k_{18c} \times Skp2_a - \frac{k_{18d} \times Skp2_a \times Cdh1_a}{(J_{18b} \times s) + Skp2_a}$ | 22 |
| $\frac{dSkp2_a}{dt} = \frac{k_{19a}}{s} \times CycE \times (Skp2_T - Skp2_a) - k_{19b} \times Skp2_a - k_{18c} \times Skp2_a - \frac{k_{18d} \times Skp2_a \times Cdh1_a}{(J_{18b} \times s) + Skp2_a}$ | 23 |
| $\frac{dCycA_m}{dt} = b_{sam} \times s + k_{sam} \times DE - k_{dam} \times CycA_m$ | 24 |
| $\frac{dCycA}{dt} = k_{sa} \times CycA_m - k_{da} \times CycA - \frac{k_{da1}}{s} \times CycA \times Cdh1_a$ | 25 |
| $\frac{dCdh1_m}{dt} = k_{3m} \times s - k_{3dm} \times Cdh1_m$ | 26 |
| $\frac{dCdh1_T}{dt} = k_3 \times Cdh1_m - k_{3d} \times Cdh1_T$ | 27 |
| $\frac{dCdh1_a}{dt} = \frac{((k_{3a} \times s) + k_{3b} \times Cdc20_a) \times (Cdh1_T - Cdh1_a)}{(J_3 \times s) + Cdh1_T - Cdh1_a} - \frac{k_4 \times CycB_a \times Cdh1_a}{(J_4 \times s) + Cdh1_a} - \frac{k_{4b} \times CycA \times Cdh1_a}{(J_4 \times s) + Cdh1_a} - k_{3d} \times Cdh1_a$ | 28 |
| $\frac{dCdc20_m}{dt} = k_{5am} \times s + \frac{k_{5bm} \times s \times \left(\frac{CycB_a}{J_5 \times s}\right)^n}{1 + \left(\frac{CycB_a}{(J_5 \times s)}\right)^n} - k_{5dm} \times Cdc20_m$ | 29 |
| $\frac{dCdc20_T}{dt} = k_{5a} \times Cdc20_m - k_6 \times Cdc20_T$ | 30 |
| $\frac{dCdc20_a}{dt} = \frac{k_7 \times IEP \times (Cdc20_T - Cdc20_a)}{(J_7 \times s) + Cdc20_T - Cdc20_a} - \frac{k_8 \times Mad \times s \times Cdc20_a}{(J_8 \times s) + Cdc20_a} - k_6 \times Cdc20_a$ | 31 |
| $\frac{dIEP_a}{dt} = \frac{k_9}{s} \times CycB_a \times ((IEP_T \times s) - IEP_a) - k_{10} \times IEP_a$ | 32 |
| $\frac{dCycB_m}{dt} = k_{scbm} \times CycA - k_{dcbm} \times CycB_m$ | 33 |
| $\frac{dCycB_T}{dt} = k_{scb} \times CycB_m - k_{dcb} \times CycB_T - \frac{k_{dcb1}}{s} \times Cdh1_a \times CycB_T$ | 34 |
| $\frac{dCycB_a}{dt} = k_{scb} \times CycB_m + \frac{k_{cba}}{s} \times C25_a \times CycB_i - \frac{k_{cbi}}{s} \times Wee1_a \times CycB_a - k_{dcb} \times CycB_a - \frac{k_{dcb1}}{s} \times Cdh1_a \times CycB_a$ | 35 |
| $\frac{dC25_m}{dt} = k_{sc25m} \times CycA - k_{dc25m} \times C25_m$ | 36 |
| $\frac{dC25_T}{dt} = k_{sc25} \times C25_m - k_{dc25} \times C25_T$ | 37 |
| $\frac{dC25_a}{dt} = \frac{k_{c25a} \times C25_i \times CycB_a}{(k_{mc1} \times s) + C25_i} - \frac{k_{c25i} \times s \times C25_a}{(k_{mc2} \times s) + C25_a} - k_{dc25} \times C25_a$ | 38 |
| $\frac{dWee1_m}{dt} = \frac{k_{swm} \times s \times Serum}{sgw + (Serum * kgw)} - k_{dwm} \times Wee1_m$ | 39 |
| $\frac{dWee1_T}{dt} = k_{sw} \times Wee1_m - k_{dw} \times Wee1_T$ | 40 |
| $\frac{dWee1_a}{dt} = k_{sw} \times Wee1_m + \frac{k_{wa} \times s \times Wee1_i}{(k_{mc1} \times s) + Wee1_i} - \frac{k_{wi} \times Wee1_a \times CycB_a}{(k_{mw2} \times s) + Wee1_a} - k_{dw} \times Wee1_a$ | 41 |

| | |
|---|---|
| $\frac{dCdt1_m}{dt} = k_{21m} \times s - k_{21dm} \times Cdt1_m$ | 42 |
| $\frac{dCdt1}{dt} = k_{21} \times Cdt1_m - k_{22a} \times Cdt1 - \frac{k_{22b} \times Cdt1 \times Skp2_a}{(J_{22} \times s) + Cdt1}$ | 43 |
| $\frac{dGem_m}{dt} = k_{20m} \times s - k_{20dm} \times Gem_m$ | 44 |
| $\frac{dGem}{dt} = k_{19} \times Gem_m - k_{20a} \times Gem - \frac{k_{20b} \times Gem \times Cdh1_a}{(J_{20} \times s) + Gem}$ | 45 |

Algebraic equations

| |
|---|
| $E2F_p = E2F_T - DE - iDE - iDEP$; $Dp1_p = (Dp_t \times s) - DE - iDE - iDEP$; $Rb = Rb_T - iRb1 - iRb2 - iDE - iDEP$ |
| $CycB_i = CycB_T - CycB_a$; $C25_i = C25_T - C25_a$; $Wee1_i = Wee1_T - Wee1_a$ |

**Table S4.** Description of the model parameters and their values:

| Description | Parameter | Values | Unit |
|---|---|---|---|
| Serum | *Serum* | 2,10 | % |
| Synthesis rate of MV Protein | $k_{mv}$ | 3 | s.u. $h^{-1}$ |
| Michaelis-Menten constant associated with MV synthesis | *smv* | 2.2 | - |
| General serum mediated activation rate of MV synthesis | *gmv* | 0.2 | - |
| Degradation rate of MV protein | $k_{dmv}$ | 4 | $h^{-1}$ |
| Basal transcription rate of Myc mRNA | $b_{myc}$ | $3.3\times10^{-5}$ | s.u. $h^{-1}$ |
| MV mediated transcription rate of Myc mRNA | $k_{Mcm1}$ | 0.033 | $h^{-1}$ |
| DE mediated transcription rate of Myc mRNA | $k_{Mcm2}$ | $1.5\times10^{-5}$ | s.u. $h^{-1}$ |
| Hill constant associated with transcription of Myc mRNA, E2F1 mRNA | $k_{mm}$ | 0.33 | s.u. |
| Hill coefficient associated with transcription of Myc mRNA, E2F1 mRNA | *n* | 2 | - |
| Degradation rate of Myc mRNA | $k_{dMcm}$ | 1.38 | $h^{-1}$ |
| Translation rate of Myc protein | $k_{Mcp}$ | 40 | $h^{-1}$ |
| Degradation rate of Myc protein | $k_{dMcp}$ | 0.7 | $h^{-1}$ |
| Skp2 mediated degradation rate of Myc protein | $k_{dMcp1}$ | 2 | $h^{-1}$ |
| Michaelis-Menten constant associated with Skp2 mediated Myc degradation | $J_{MS}$ | 0.001 | s.u. |
| Basal transcription rate of E2F1 mRNA | $b_{E2F}$ | $3.3\times10^{-6}$ | s.u. $h^{-1}$ |
| Myc mediated transcription rate of E2F1 mRNA | $k_{Em1}$ | 0.005 | $h^{-1}$ |
| DE Mediated transcription rate of E2F1 mRNA | $k_{Em2}$ | 0.0005 | s.u. $h^{-1}$ |
| Degradation rate of E2F1 mRNA | $k_{dEm}$ | 0.25 | $h^{-1}$ |
| Translation rate of E2F1 protein | $k_{Ep1}$ | 50 | $h^{-1}$ |
| Degradation rate of E2F1 protein, DE, IDE | $k_{dEp}$ | 0.25 | $h^{-1}$ |
| Cyclin A mediated degradation rate of E2F1 protein | $k_{dEp1}$ | 1 | $h^{-1}$ |
| Michaelis-Menten constant associated with Cyclin A mediated DE degradation | $J_{DE}$ | 1 | s.u. |
| Dp1 and E2F1 association constant | $k_{DE1}$ | 871.4 | $s.u.^{-1}$ $h^{-1}$ |
| DE dissociation constant | $k_{DE2}$ | 55 | $h^{-1}$ |
| Total Dp1 protein | $Dp1_t$ | 1 | s.u. |
| Transcription rate of Rb mRNA | $k_{Rbm}$ | 0.05 | s.u. $h^{-1}$ |
| Degradation rate of Rb mRNA | $k_{dRbm}$ | 1.04 | $h^{-1}$ |
| Translation rate of Rb protein | $k_{Rbp}$ | 5 | $h^{-1}$ |
| Degradation rate of Rb, Rbp, Rbpp protein | $k_{dRbp}$ | 0.231 | $h^{-1}$ |
| DE and Rb association constant | $k_{RD1}$ | 100 | $s.u.^{-1}$ $h^{-1}$ |
| IDE dissociation constant | $k_{RD2}$ | 0.5 | $h^{-1}$ |
| DE and Rbp association constant | $k_{RpD1}$ | 0.1 | $s.u.^{-1}$ $h^{-1}$ |

| | | | |
|---|---|---|---|
| IDEP dissociation constant | $k_{RpD2}$ | 0.05 | $h^{-1}$ |
| Cyclin D mediated phosphorylation rate of free Rb and Rb bound to DE | $k_{iR1}$ | 0.8 | $h^{-1}$ |
| Michaelis-Menten constant associated with Cyclin D mediated phosphorylation of free Rb and Rb bound to DE | $J_{RD}$ | 0.01 | s.u. |
| Dephosphorylation rate of free Rbp and Rbp bound to DE | $k_{iR2}$ | 1 | s.u. $h^{-1}$ |
| Michaelis-Menten constant associated with dephosphorylation of free Rbp and Rbp bound to DE | $J_{R1}$ | 0.01 | s.u. |
| Cyclin E mediated phosphorylation rate of free Rbp and Rbp bound to DE | $k_{iR3}$ | 60 | $h^{-1}$ |
| Michaelis-Menten constant associated with Cyclin E mediated phosphorylation of free Rbp and Rbp bound to DE | $J_{RE}$ | 0.001 | s.u. |
| Dephosphorylation rate of free Rbpp | $k_{iR4}$ | 25 | s.u. $h^{-1}$ |
| Michaelis-Menten constant associated with dephosphorylation of free Rbpp | $J_{R2}$ | 0.01 | s.u. |
| MV mediated transcription rate of Cyclin D mRNA | $k_{CDm1}$ | $1.0\times10^{-6}$ | $h^{-1}$ |
| Myc mediated transcription rate of Cyclin D mRNA | $k_{CDm2}$ | 0.01 | $h^{-1}$ |
| Degradation rate of Cyclin D mRNA | $k_{dCDm}$ | 0.173 | $h^{-1}$ |
| Translation rate of Cyclin D protein | $k_{CDp1}$ | 56.67 | $h^{-1}$ |
| Degradation rate of Cyclin D protein | $k_{dCDp}$ | 1.386 | $h^{-1}$ |
| Cyclin A mediated degradation rate of Cyclin D protein | $k_{dCDp1}$ | 2 | $h^{-1}$ |
| Michaelis-Menten constant associated with Cyclin A mediated Cyclin D degradation | $J_{CD}$ | 0.01 | s.u. |
| Basal transcription rate of Cyclin E mRNA | $b_{CycEm}$ | $2.0\times10^{-6}$ | s.u. $h^{-1}$ |
| DE mediated transcription rate of Cyclin E mRNA | $k_{sem}$ | 0.04 | $h^{-1}$ |
| Degradation rate of Cyclin E mRNA | $k_{dcem}$ | 1 | $h^{-1}$ |
| Translation rate of Cyclin E protein | $k_{se}$ | 100 | $h^{-1}$ |
| Degradation rate of Cyclin E protein | $k_{de}$ | 1 | $h^{-1}$ |
| Skp2 mediated degradation rate of Cyclin E protein | $k_{dea}$ | 2 | s.u.$^{-1}$ $h^{-1}$ |
| Transcription rate of CKI mRNA | $k_{sim}$ | 0.0142 | s.u. $h^{-1}$ |
| Degradation rate of CKI mRNA | $k_{dim}$ | 0.693 | $h^{-1}$ |
| Translation rate of CKI protein | $k_{si}$ | 100 | $h^{-1}$ |
| Degradation rate of CKI protein | $k_{di}$ | 1 | $h^{-1}$ |
| Cyclin E and CKI association constant | $k_{aei}$ | 10 | s.u.$^{-1}$ $h^{-1}$ |
| Cyclin E: CKI dissociation constant | $k_{dei}$ | 1 | $h^{-1}$ |
| Cyclin E mediated phosphorylation rate of CKI | $k_{ip}$ | 5 | $h^{-1}$ |

| | | | |
|---|---|---|---|
| Michaelis-Menten constant associated with Cyclin E mediated CKI phosphorylation | $J_{ce}$ | 0.1 | s.u. |
| Skp2 mediated degradation rate of CKI protein from Cyclin E: CKI complex | $k_{dip1}$ | 1 | $h^{-1}$ |
| Skp2 mediated degradation rate of phosphorylated CKI | $k_{dip}$ | 20 | $s.u.^{-1}$ $h^{-1}$ |
| Transcription rate of Skp2 mRNA | $k_{17m}$ | 0.0036 | s.u. $h^{-1}$ |
| Degradation rate of Skp2 mRNA | $k_{17dm}$ | 0.17 | $h^{-1}$ |
| Translation rate of Skp2 protein | $k_{17}$ | 26.471 | $h^{-1}$ |
| Degradation rate of dephosphorylated Skp2 protein | $k_{18a}$ | 19.412 | $h^{-1}$ |
| Cdh1 mediated degradation rate of dephosphorylated Skp2 protein | $k_{18b}$ | 52.941 | $h^{-1}$ |
| Michaelis-Menten constant associated with Cdh1 mediated dephosphorylated Skp2 degradation | $J_{18}$ | 0.001 | s.u. |
| Cyclin E mediated phosphorylation rate of Skp2 | $k_{19a}$ | 70.588 | $s.u.^{-1}$ $h^{-1}$ |
| Dephosphorylation rate of Skp2 | $k_{19b}$ | 0.35294 | $h^{-1}$ |
| Degradation rate of phosphorylated Skp2 | $k_{18c}$ | 0.028235 | $h^{-1}$ |
| Cdh1 mediated degradation rate of phosphorylated Skp2 | $k_{18d}$ | 0.35294 | $h^{-1}$ |
| Michaelis-Menten constant associated with Cdh1 mediated phosphorylated Skp2 degradation | $J_{18b}$ | 0.01 | s.u. |
| Basal transcription rate of Cyclin A mRNA | $b_{sam}$ | $1.0 \times 10^{-6}$ | s.u. $h^{-1}$ |
| DE mediated transcription rate of Cyclin A mRNA | $k_{sam}$ | 0.02 | $h^{-1}$ |
| Degradation rate of Cyclin A mRNA | $k_{dam}$ | 1 | $h^{-1}$ |
| Translation rate of Cyclin A protein | $k_{sa}$ | 7.0588 | $h^{-1}$ |
| Degradation rate of Cyclin A protein | $k_{da}$ | 0.14118 | $h^{-1}$ |
| Cdh1 mediated degradation rate of Cyclin A protein | $k_{da1}$ | 3.5294 | $s.u.^{-1}$ $h^{-1}$ |
| Transcription rate of Cdh1 mRNA | $k_{3m}$ | 0.06 | s.u. $h^{-1}$ |
| Degradation rate of Cdh1 mRNA | $k_{3dm}$ | 3 | $h^{-1}$ |
| Translation rate of Cdh1protein | $k_{3}$ | 176.47 | $h^{-1}$ |
| Degradation rate of Cdh1 protein | $k_{3d}$ | 3.5294 | $h^{-1}$ |
| Basal dephosphorylation rate of Cdh1 protein | $k_{3a}$ | 3.5294 | s.u. $h^{-1}$ |
| Cdc20 mediated dephosphorylation rate of Cdh1 protein | $k_{3b}$ | 176.47 | $h^{-1}$ |
| Michaelis-Menten constant associated with Cdc20 mediated Cdh1 dephosphorylation | $J_{3}$ | 0.04 | s.u. |
| Cyclin B mediated phosphorylation rate of Cdh1 | $k_{4}$ | 123.53 | $h^{-1}$ |
| Cyclin A mediated phosphorylation rate of Cdh1 | $k_{4b}$ | 123.53 | $h^{-1}$ |
| Michaelis-Menten constant associated with Cyclin A mediated Cdh1 phosphorylation | $J_{4}$ | 0.04 | s.u. |
| Basal synthesis rate of Cdc20 mRNA | $k_{5am}$ | 0.00035294 | s.u. $h^{-1}$ |
| Cyclin B mediated transcription rate of Cdc20 mRNA | $k_{5bm}$ | 0.014118 | s.u. $h^{-1}$ |

| | | | |
|---|---|---|---|
| Hill constant associated with Cyclin B mediated Cdc20 transcription | $J_5$ | 0.3 | s.u. |
| Hill coefficient associated with Cyclin B mediated Cdc20 transcription | $m$ | 4 | - |
| Degradation rate of Cdc20 mRNA | $k_{5dm}$ | 0.35294 | $h^{-1}$ |
| Translation rate of Cdc20 protein | $k_{5a}$ | 17.647 | $h^{-1}$ |
| Degradation rate of Cdc20 protein | $k_6$ | 0.35294 | $h^{-1}$ |
| IEP mediated activation rate of Cdc20 protein | $k7$ | 3.5294 | $h^{-1}$ |
| Michaelis-Menten constant associated with IEP mediated activation of Cdc20 protein | $J_7$ | 0.05 | s.u. |
| Mad mediated deactivation rate of Cdc20 protein | $k_8$ | 1.7647 | $h^{-1}$ |
| Michaelis-Menten constant associated with Mad mediated deactivation of Cdc20 protein | $J_8$ | 0.05 | s.u. |
| Cyclin B mediated phosphorylation rate of IEP | $k_9$ | 0.35294 | $s.u.^{-1}\ h^{-1}$ |
| Dephosphorylation rate of IEP | $k_{10}$ | 0.070588 | $h^{-1}$ |
| Total Mad1 protein | $Mad$ | 1 | s.u. |
| Total IEP protein | $IEP_T$ | 1 | s.u. |
| Transcription rate of Cdt1 mRNA | $k_{21m}$ | 0.0075 | $s.u.\ h^{-1}$ |
| Degradation rate of Cdt1 mRNA | $k_{21dm}$ | 0.35 | $h^{-1}$ |
| Translation rate of Cdt1 protein | $k_{21}$ | 8.8235 | $h^{-1}$ |
| Degradation rate of Cdt1 protein | $k_{22a}$ | 0.21176 | $h^{-1}$ |
| Skp2 mediated degradation rate of Cdt1protein | $k_{22b}$ | 7.0588 | $h^{-1}$ |
| Michaelis-Menten constant associated with Skp2 mediated degradation of Cdt1 protein | $J_{22}$ | 0.04 | s.u. |
| Transcription rate of Geminin mRNA | $k_{20m}$ | 0.0075 | $s.u.\ h^{-1}$ |
| Degradation rate of Geminin mRNA | $k_{20dm}$ | 0.35 | $h^{-1}$ |
| Translation rate of Geminin protein | $k_{19}$ | 8.8235 | $h^{-1}$ |
| Degradation rate of Geminin protein | $k_{20a}$ | 0.21176 | $h^{-1}$ |
| Cdh1 mediated degradation rate of Geminin protein | $k_{20b}$ | 3.5294 | $h^{-1}$ |
| Michaelis-Menten constant associated with Cdh1 mediated degradation of Geminin protein | $J_{20}$ | 0.5 | s.u. |
| Cyclin A mediated transcription rate of Cyclin B mRNA | $k_{scbm}$ | 0.02 | $h^{-1}$ |
| Degradation rate of Cyclin B mRNA | $k_{dcbm}$ | 0.1 | $h^{-1}$ |
| Translation rate of Cyclin B protein | $k_{scb}$ | 3.5294 | $h^{-1}$ |
| Degradation rate of Cyclin B protein | $k_{dcb}$ | 0.14118 | $h^{-1}$ |
| Cdh1 mediated degradation rate of Cyclin B protein | $k_{dcb1}$ | 3.5294 | $s.u.^{-1}\ h^{-1}$ |
| Cdc25 mediate phosphorylation rate of Cyclin B | $k_{cba}$ | 35.294 | $s.u.^{-1}\ h^{-1}$ |
| Wee1 mediated dephosphorylation rate of Cyclin B | $k_{cbi}$ | 35.294 | $s.u.^{-1}\ h^{-1}$ |
| Cyclin A mediated transcription rate of Cdc25 mRNA | $k_{sc25m}$ | 0.02 | $h^{-1}$ |
| Degradation rate of Cdc25 mRNA | $k_{dc25m}$ | 1 | $h^{-1}$ |

| | | | |
|---|---|---|---|
| Translation rate of Cdc25 protein | $k_{sc25}$ | 176.47 | $h^{-1}$ |
| Degradation rate of Cdc25 protein | $k_{dc25}$ | 3.5294 | $h^{-1}$ |
| Cyclin B mediate phosphorylation rate of Cdc25 | $k_{c25a}$ | 35.294 | $h^{-1}$ |
| Dephosphorylation rate of Cdc25 | $k_{c25i}$ | 1.4118 | s.u. $h^{-1}$ |
| Michaelis-Menten constant associated with Cyclin B mediate phosphorylation of Cdc25 | $k_{mc1}$ | 0.05 | s.u. |
| Michaelis-Menten constant associated with dephosphorylation of Cdc25 | $k_{mc2}$ | 0.05 | s.u. |
| Transcription rate of Wee1 mRNA | $k_{swm}$ | 0.016 | s.u. $h^{-1}$ |
| Michaelis-Menten constant associated with Wee1 mRNA transcription | *sgw* | 1 | - |
| General serum mediated activation rate of Wee1 transcription | *kgw* | 0.5 | - |
| Degradation rate of Wee1 mRNA | $k_{dwm}$ | 1 | $h^{-1}$ |
| Translation rate of Wee1 protein | $k_{sw}$ | 176.47 | $h^{-1}$ |
| Degradation rate of Wee1 protein | $k_{dw}$ | 3.5294 | $h^{-1}$ |
| Dephosphorylation rate of Wee1 | $k_{wa}$ | 14.118 | s.u. $h^{-1}$ |
| Cyclin B mediate phosphorylation rate of Wee1 | $k_{wi}$ | 176.47 | $h^{-1}$ |
| Michaelis-Menten constant associated with dephosphorylation of Wee1 | $k_{mw1}$ | 0.2 | s.u. |
| Michaelis-Menten constant associated with Cyclin B mediate phosphorylation of Wee1 | $k_{mw2}$ | 0.2 | s.u. |
| General p38i mediated inhibition rate of Myc, Cyclin D and CKI transcription | $k_{p1}$ | 0.1 | - |
| General p38i mediated inhibition rate of Cdc25 translation | $k_{p2}$ | 0.35 | - |
| Amount of p38 inhibitor | *p38i* | 0,10 | - |
| Scaling factor<br>( **1 s.u.** is considered as 10000 molecules) | *s* | 10000 | Molecules |
| Confluency factor (increases in 0.4 in each cycle) | *cf* | 1 (increases in 0.4 for each cycle) | - |

**Table S5.** Table showing the data from stochastic simulation (Model, M-1, Table-1 with unequal division) at 2% and 10% serum**.**

| Serum (%) | | $T_{CC}$ | | | $T_{G1}$ | | | $T_{S\text{-}G2\text{-}M}$ | | | $T_{CC}$ vs $T_{G1}$ | $T_{CC}$ vs $T_{S\text{-}G2\text{-}M}$ | $T_{G1}$ vs $T_{S\text{-}G2\text{-}M}$ | n |
|---|---|---|---|---|---|---|---|---|---|---|---|---|---|---|
| | | Mean (h) | SD (h) | CV (%) | Mean (h) | SD (h) | CV (%) | Mean (h) | SD (h) | CV (%) | $R^2$ | $R^2$ | $R^2$ | |
| 2 | I | 29.3 | 7.4 | 25.3 | 13.3 | 4.9 | 37.1 | 16.0 | 4.6 | 29.0 | 0.624 | 0.574 | 0.039 | 300 |
| | II | 27.7 | 7.0 | 25.4 | 12.7 | 4.8 | 37.5 | 15.1 | 4.0 | 26.7 | 0.700 | 0.582 | 0.081 | 300 |
| | III | 27.0 | 7.3 | 26.9 | 11.8 | 4.4 | 36.9 | 15.3 | 4.4 | 28.8 | 0.689 | 0.694 | 0.146 | 300 |
| | avg | 28.0 | 7.2 | 25.9 | 12.6 | 4.7 | 37.2 | 15.5 | 4.3 | 28.2 | 0.671 | 0.617 | 0.089 | |

| Serum (%) | | $T_{CC}$ | | | $T_{G1}$ | | | $T_{S\text{-}G2\text{-}M}$ | | | $T_{CC}$ vs $T_{G1}$ | $T_{CC}$ vs $T_{S\text{-}G2\text{-}M}$ | $T_{G1}$ vs $T_{S\text{-}G2\text{-}M}$ | n |
|---|---|---|---|---|---|---|---|---|---|---|---|---|---|---|
| | | Mean (h) | SD (h) | CV (%) | Mean (h) | SD (h) | CV (%) | Mean (h) | SD (h) | CV (%) | $R^2$ | $R^2$ | $R^2$ | |
| 10 | I | 25.0 | 6.3 | 25 | 8.9 | 3.0 | 33.9 | 16.1 | 5.3 | 33.2 | 0.271 | 0.769 | 0.002 | 300 |
| | II | 24.2 | 5.5 | 22.6 | 8.5 | 2.6 | 31.0 | 15.7 | 4.6 | 29.6 | 0.281 | 0.768 | 0.003 | 300 |
| | III | 23.7 | 5.1 | 21.6 | 8.4 | 2.7 | 32.8 | 15.3 | 4.1 | 27.0 | 0.357 | 0.714 | 0.006 | 300 |
| | avg | 24.3 | 5.6 | 23.1 | 8.6 | 2.8 | 32.6 | 15.7 | 4.7 | 29.9 | 0.303 | 0.750 | 0.004 | |

| Serum (%) | | Sister pairs (Pearson, R) | | | | Cousin pairs (Pearson, R) | | | | Mother-daughter pairs (Pearson, R) | | | |
|---|---|---|---|---|---|---|---|---|---|---|---|---|---|
| | | $T_{CC}$ | $T_{G1}$ | $T_{S\text{-}G2\text{-}M}$ | n | $T_{CC}$ | $T_{G1}$ | $T_{S\text{-}G2\text{-}M}$ | n | $T_{CC}$ | $T_{G1}$ | $T_{S\text{-}G2\text{-}M}$ | n |
| 2 | I | 0.698 | 0.575 | 0.65 | 131 | 0.219 | 0.282 | 0.12 | 73 | 0.391 | 0.356 | 0.475 | 129 |
| | II | 0.832 | 0.854 | 0.614 | 136 | 0.394 | 0.363 | 0.352 | 83 | 0.322 | 0.343 | 0.391 | 146 |
| | III | 0.835 | 0.822 | 0.75 | 130 | 0.146 | 0.073 | 0.266 | 81 | 0.419 | 0.382 | 0.494 | 147 |
| | avg | 0.788 | 0.750 | 0.671 | | 0.253 | 0.239 | 0.246 | | 0.377 | 0.360 | 0.453 | |

| Serum (%) | | Sister pairs (Pearson, R) | | | | Cousin pairs (Pearson, R) | | | | Mother-daughter pairs (Pearson, R) | | | |
|---|---|---|---|---|---|---|---|---|---|---|---|---|---|
| | | $T_{CC}$ | $T_{G1}$ | $T_{S\text{-}G2\text{-}M}$ | n | $T_{CC}$ | $T_{G1}$ | $T_{S\text{-}G2\text{-}M}$ | n | $T_{CC}$ | $T_{G1}$ | $T_{S\text{-}G2\text{-}M}$ | n |
| 10 | I | 0.707 | 0.566 | 0.509 | 132 | 0.388 | 0.579 | 0.384 | 123 | 0.491 | 0.568 | 0.591 | 191 |
| | II | 0.806 | 0.881 | 0.752 | 130 | 0.163 | 0.202 | 0.382 | 98 | 0.456 | 0.406 | 0.667 | 182 |
| | III | 0.626 | 0.756 | 0.556 | 134 | 0.284 | 0.375 | 0.289 | 133 | 0.532 | 0.513 | 0.607 | 195 |
| | avg | 0.713 | 0.734 | 0.606 | | 0.279 | 0.385 | 0.352 | | 0.493 | 0.496 | 0.622 | |

**Table S6.** Table showing the simulation data from ***Replicate I*** of all model variants (Table-1) with unequal division at 2% and 10% serum.

| Model Type | Serum Level (%) | $T_{CC}$ | | | $T_{G1}$ | | | $T_{S\text{-}G2\text{-}M}$ | | | $T_{CC}$ vs $T_{G1}$ | $T_{CC}$ vs $T_{S\text{-}G2\text{-}M}$ | $T_{G1}$ vs $T_{S\text{-}G2\text{-}M}$ | n |
|---|---|---|---|---|---|---|---|---|---|---|---|---|---|---|
| | | Mean (h) | SD (h) | CV (%) | Mean (h) | SD (h) | CV (%) | Mean (h) | SD (h) | CV (%) | $R^2$ | $R^2$ | $R^2$ | |
| **M-1** | 2 | 29.3 | 7.4 | 25.3 | 13.3 | 4.9 | 37.1 | 16 | 4.6 | 29.0 | 0.624 | 0.574 | 0.039 | 300 |
| | 10 | 25 | 6.3 | 25.0 | 8.9 | 3 | 33.9 | 16.1 | 5.3 | 33.2 | 0.271 | 0.769 | 0.002 | 300 |
| **M-2** | 2 | 28.1 | 2.2 | 7.7 | 13.9 | 1.7 | 11.9 | 14.2 | 0.8 | 5.7 | 0.888 | 0.531 | 0.209 | 300 |
| | 10 | 23.4 | 0.8 | 3.6 | 8.5 | 0.8 | 9.6 | 14.8 | 0.5 | 3.4 | 0.663 | 0.1 | 0.086 | 300 |
| **M-3** | 2 | 26.4 | 3.4 | 12.9 | 12.3 | 2.1 | 16.9 | 14.1 | 1.8 | 12.9 | 0.787 | 0.723 | 0.262 | 300 |
| | 10 | 23.6 | 2.8 | 11.8 | 8.5 | 1.3 | 15.3 | 15 | 2.1 | 14.1 | 0.467 | 0.797 | 0.079 | 300 |
| **M-4** | 2 | 25.5 | 4.2 | 16.3 | 11.6 | 3.1 | 26.5 | 13.9 | 2.0 | 14.3 | 0.795 | 0.511 | 0.103 | 300 |
| | 10 | 24.2 | 4.4 | 18.1 | 8.5 | 2.2 | 25.5 | 15.7 | 3.6 | 23.0 | 0.326 | 0.753 | 0.008 | 300 |
| **M-5** | 2 | 27.9 | 3.5 | 12.5 | 13.6 | 2.5 | 18.2 | 14.3 | 1.9 | 13.4 | 0.72 | 0.531 | 0.066 | 300 |
| | 10 | 23.3 | 2.4 | 10.3 | 8.6 | 1.3 | 14.9 | 14.7 | 1.9 | 12.9 | 0.382 | 0.718 | 0.011 | 300 |
| **M-6** | 2 | 29.3 | 7.4 | 25.3 | 13.3 | 4.9 | 37.1 | 16.0 | 4.6 | 29.0 | 0.624 | 0.574 | 0.039 | 300 |
| | 10 | 23.5 | 5.0 | 21.3 | 8.7 | 3.1 | 35.2 | 14.8 | 3.8 | 25.7 | 0.425 | 0.629 | 0.003 | 300 |

| Model Type | Serum Level (%) | Sister pairs (Pearson, R) | | | | Cousin pairs (Pearson, R) | | | | Mother-daughter pairs (Pearson, R) | | | |
|---|---|---|---|---|---|---|---|---|---|---|---|---|---|
| | | $T_{CC}$ | $T_{G1}$ | $T_{S\text{-}G2\text{-}M}$ | n | $T_{CC}$ | $T_{G1}$ | $T_{S\text{-}G2\text{-}M}$ | n | $T_{CC}$ | $T_{G1}$ | $T_{S\text{-}G2\text{-}M}$ | n |
| **M-1** | 2 | 0.698 | 0.575 | 0.65 | 131 | 0.219 | 0.282 | 0.12 | 73 | 0.391 | 0.356 | 0.475 | 129 |
| | 10 | 0.707 | 0.566 | 0.509 | 132 | 0.388 | 0.579 | 0.384 | 123 | 0.491 | 0.568 | 0.591 | 191 |
| **M-2** | 2 | 0.671 | 0.638 | 0.274 | 148 | -0.164 | -0.14 | 0.003 | 192 | -0.016 | 0.063 | -0.043 | 198 |
| | 10 | -0.185 | 0.047 | 0.029 | 150 | -0.009 | -0.021 | 0.035 | 200 | 0.171 | 0.122 | 0.082 | 200 |
| **M-3** | 2 | 0.998 | 0.992 | 0.987 | 149 | 0.998 | 0.996 | 0.992 | 194 | 0.97 | 0.965 | 0.967 | 196 |
| | 10 | 0.94 | 0.679 | 0.995 | 146 | 0.969 | 0.799 | 0.997 | 204 | 0.957 | 0.769 | 0.973 | 209 |
| **M-4** | 2 | 0.934 | 0.991 | 0.69 | 136 | 0.565 | 0.587 | 0.195 | 98 | 0.507 | 0.527 | 0.541 | 168 |
| | 10 | 0.686 | 0.864 | 0.595 | 136 | 0.464 | 0.395 | 0.32 | 157 | 0.528 | 0.492 | 0.582 | 201 |
| **M-5** | 2 | 0.863 | 0.832 | 0.762 | 139 | 0.805 | 0.771 | 0.671 | 165 | 0.784 | 0.816 | 0.745 | 180 |
| | 10 | 0.836 | 0.577 | 0.921 | 145 | 0.892 | 0.735 | 0.919 | 197 | 0.882 | 0.619 | 0.924 | 211 |
| **M-6** | 2 | 0.698 | 0.575 | 0.65 | 131 | 0.219 | 0.282 | 0.12 | 73 | 0.391 | 0.356 | 0.475 | 129 |
| | 10 | 0.617 | 0.629 | 0.494 | 136 | 0.582 | 0.364 | 0.53 | 117 | 0.556 | 0.573 | 0.659 | 195 |

**Table S7.** Table showing the effect of varying lognormal distribution CV for transcription rates in Model, M-3 (Table-1) with unequal division.

| CV (%) | Serum Level (%) | $T_{CC}$ | | | $T_{G1}$ | | | $T_{S\text{-}G2\text{-}M}$ | | | $T_{CC}$ vs $T_{G1}$ | $T_{CC}$ vs $T_{S\text{-}G2\text{-}M}$ | $T_{G1}$ vs $T_{S\text{-}G2\text{-}M}$ | n |
|---|---|---|---|---|---|---|---|---|---|---|---|---|---|---|
| | | Mean (h) | SD (h) | CV (%) | Mean (h) | SD (h) | CV (%) | Mean (h) | SD (h) | CV (%) | $R^2$ | $R^2$ | $R^2$ | |
| **10** | 2 | 26.9 | 2.5 | 9.2 | 12.9 | 1.7 | 13.5 | 14 | 1.0 | 7.2 | 0.888 | 0.665 | 0.332 | 300 |
| | 10 | 23.3 | 1.3 | 5.5 | 8.5 | 0.9 | 10.6 | 14.8 | 0.9 | 6 | 0.53 | 0.514 | 0.002 | 300 |
| **20** | 2 | 26.4 | 3.4 | 12.9 | 12.3 | 2.1 | 16.9 | 14.1 | 1.8 | 12.9 | 0.787 | 0.723 | 0.262 | 300 |
| | 10 | 23.6 | 2.8 | 11.8 | 8.5 | 1.3 | 15.3 | 15 | 2.1 | 14.1 | 0.467 | 0.797 | 0.079 | 300 |
| **30** | 2 | 25.7 | 3.7 | 14.5 | 11.9 | 2.6 | 22.1 | 13.8 | 2.0 | 14.6 | 0.732 | 0.541 | 0.078 | 300 |
| | 10 | 23.3 | 3.7 | 16.1 | 8.8 | 1.9 | 21.9 | 14.5 | 2.6 | 17.8 | 0.588 | 0.77 | 0.133 | 300 |
| **40** | 2 | 25.2 | 4.8 | 18.9 | 11.6 | 3.1 | 27 | 13.7 | 2.7 | 19.5 | 0.726 | 0.627 | 0.126 | 300 |
| | 10 | 22.9 | 3.6 | 16 | 8.4 | 1.8 | 21.1 | 14.5 | 2.9 | 20.2 | 0.378 | 0.771 | 0.026 | 300 |

| CV (%) | Serum Level (%) | Sister pairs (Pearson, R) | | | | Cousin pairs (Pearson, R) | | | | Mother-daughter pairs (Pearson, R) | | | |
|---|---|---|---|---|---|---|---|---|---|---|---|---|---|
| | | $T_{CC}$ | $T_{G1}$ | $T_{S\text{-}G2\text{-}M}$ | n | $T_{CC}$ | $T_{G1}$ | $T_{S\text{-}G2\text{-}M}$ | n | $T_{CC}$ | $T_{G1}$ | $T_{S\text{-}G2\text{-}M}$ | n |
| **10** | 2 | 0.998 | 0.993 | 0.958 | 150 | 0.999 | 0.997 | 0.979 | 200 | 0.98 | 0.977 | 0.946 | 200 |
| | 10 | 0.762 | 0.347 | 0.984 | 149 | 0.884 | 0.508 | 0.99 | 204 | 0.86 | 0.454 | 0.973 | 206 |
| **20** | 2 | 0.998 | 0.992 | 0.987 | 149 | 0.998 | 0.996 | 0.992 | 194 | 0.97 | 0.965 | 0.967 | 196 |
| | 10 | 0.94 | 0.679 | 0.995 | 146 | 0.969 | 0.799 | 0.997 | 204 | 0.957 | 0.769 | 0.973 | 209 |
| **30** | 2 | 0.996 | 0.988 | 0.988 | 149 | 0.998 | 0.994 | 0.994 | 189 | 0.963 | 0.961 | 0.957 | 190 |
| | 10 | 0.955 | 0.811 | 0.996 | 149 | 0.976 | 0.898 | 0.996 | 210 | 0.968 | 0.881 | 0.979 | 214 |
| **40** | 2 | 0.996 | 0.994 | 0.987 | 150 | 0.996 | 0.992 | 0.992 | 204 | 0.959 | 0.948 | 0.963 | 204 |
| | 10 | 0.967 | 0.836 | 0.994 | 148 | 0.983 | 0.914 | 0.995 | 216 | 0.961 | 0.871 | 0.975 | 224 |

**Table S8.** Table showing the effect of variation in % change of transcription rate during cell cycle in Model, M-4 (Table-1) with unequal division.

| % change in TR | Serum Level (%) | $T_{CC}$ | | | $T_{G1}$ | | | $T_{S\text{-}G2\text{-}M}$ | | | $T_{CC}$ vs $T_{G1}$ | $T_{CC}$ vs $T_{S\text{-}G2\text{-}M}$ | $T_{G1}$ vs $T_{S\text{-}G2\text{-}M}$ | n |
|---|---|---|---|---|---|---|---|---|---|---|---|---|---|---|
| | | Mean (h) | SD (h) | CV (%) | Mean (h) | SD (h) | CV (%) | Mean (h) | SD (h) | CV (%) | $R^2$ | $R^2$ | $R^2$ | |
| **1 to 5** | 2 | 26.7 | 3.1 | 11.6 | 12.4 | 2.3 | 18.1 | 14.3 | 1.7 | 11.7 | 0.729 | 0.501 | 0.056 | 300 |
| | 10 | 23.5 | 2.3 | 9.8 | 8.7 | 1.2 | 13.7 | 14.8 | 1.7 | 11.8 | 0.457 | 0.747 | 0.045 | 300 |
| **2 to 10** | 2 | 25.8 | 3.0 | 11.7 | 11.9 | 2.1 | 17.5 | 13.9 | 1.6 | 11.4 | 0.759 | 0.582 | 0.121 | 300 |
| | 10 | 23.9 | 2.9 | 12.3 | 8.5 | 1.4 | 16.4 | 15.4 | 2.1 | 13.6 | 0.566 | 0.810 | 0.152 | 300 |
| **3 to 15** | 2 | 25.9 | 3.7 | 14.2 | 12.4 | 2.9 | 23.3 | 13.5 | 1.7 | 12.8 | 0.787 | 0.408 | 0.045 | 300 |
| | 10 | 23.3 | 3.3 | 13.9 | 8.2 | 1.4 | 17.8 | 15.2 | 2.6 | 16.8 | 0.425 | 0.815 | 0.069 | 300 |
| **4 to 20** | 2 | 25.5 | 3.5 | 13.7 | 11.8 | 2.6 | 21.7 | 13.7 | 1.7 | 12.6 | 0.781 | 0.517 | 0.096 | 300 |
| | 10 | 23.4 | 3.2 | 13.7 | 8.4 | 1.6 | 19.4 | 15.0 | 2.6 | 17.2 | 0.361 | 0.747 | 0.014 | 300 |
| **5 to 25** | 2 | 26.7 | 4.0 | 14.9 | 12.4 | 3.0 | 24.6 | 14.3 | 2.0 | 14.1 | 0.750 | 0.431 | 0.037 | 300 |
| | 10 | 23.4 | 3.6 | 15.5 | 8.2 | 2.0 | 24.0 | 15.2 | 2.8 | 18.4 | 0.412 | 0.708 | 0.016 | 300 |
| **6 to 30** | 2 | 25.5 | 4.2 | 16.3 | 11.6 | 3.1 | 26.5 | 13.9 | 2.0 | 14.3 | 0.795 | 0.511 | 0.103 | 300 |
| | 10 | 24.2 | 4.4 | 18.1 | 8.5 | 2.2 | 25.5 | 15.7 | 3.6 | 23.0 | 0.326 | 0.753 | 0.008 | 300 |

| % change in TR | Serum Level (%) | Sister pairs (Pearson, R) | | | | Cousin pairs (Pearson, R) | | | | Mother-daughter pairs (Pearson, R) | | | |
|---|---|---|---|---|---|---|---|---|---|---|---|---|---|
| | | $T_{CC}$ | $T_{G1}$ | $T_{S\text{-}G2\text{-}M}$ | n | $T_{CC}$ | $T_{G1}$ | $T_{S\text{-}G2\text{-}M}$ | n | $T_{CC}$ | $T_{G1}$ | $T_{S\text{-}G2\text{-}M}$ | n |
| **1 to 5** | 2 | 0.987 | 0.990 | 0.947 | 147 | 0.907 | 0.907 | 0.890 | 186 | 0.897 | 0.927 | 0.903 | 193 |
| | 10 | 0.912 | 0.670 | 0.966 | 146 | 0.89 | 0.742 | 0.904 | 203 | 0.894 | 0.702 | 0.942 | 214 |
| **2 to 10** | 2 | 0.975 | 0.985 | 0.889 | 146 | 0.714 | 0.666 | 0.647 | 168 | 0.816 | 0.785 | 0.84 | 189 |
| | 10 | 0.882 | 0.696 | 0.902 | 148 | 0.770 | 0.657 | 0.753 | 198 | 0.868 | 0.733 | 0.878 | 208 |
| **3 to 15** | 2 | 0.972 | 0.994 | 0.858 | 144 | 0.503 | 0.617 | 0.453 | 156 | 0.641 | 0.718 | 0.748 | 185 |
| | 10 | 0.868 | 0.708 | 0.859 | 143 | 0.695 | 0.626 | 0.628 | 195 | 0.799 | 0.589 | 0.838 | 211 |
| **4 to 20** | 2 | 0.946 | 0.991 | 0.782 | 143 | 0.501 | 0.468 | 0.395 | 150 | 0.732 | 0.720 | 0.713 | 190 |
| | 10 | 0.812 | 0.738 | 0.771 | 141 | 0.402 | 0.356 | 0.437 | 177 | 0.628 | 0.500 | 0.698 | 209 |
| **5 to 25** | 2 | 0.902 | 0.991 | 0.607 | 136 | 0.459 | 0.456 | 0.333 | 125 | 0.595 | 0.623 | 0.646 | 172 |
| | 10 | 0.761 | 0.837 | 0.729 | 141 | 0.432 | 0.556 | 0.344 | 167 | 0.652 | 0.682 | 0.683 | 208 |
| **6 to 30** | 2 | 0.934 | 0.991 | 0.690 | 136 | 0.565 | 0.587 | 0.195 | 98 | 0.507 | 0.527 | 0.541 | 168 |
| | 10 | 0.686 | 0.864 | 0.595 | 136 | 0.464 | 0.395 | 0.320 | 157 | 0.528 | 0.492 | 0.582 | 201 |

**Table S9.** Table showing simulation data from ***Replicate I*** of all model variants (Table-1) with equal division at 2% and 10% serum.

| Model Type | Serum Level (%) | $T_{CC}$ | | | $T_{G1}$ | | | $T_{S\text{-}G2\text{-}M}$ | | | $T_{CC}$ vs $T_{G1}$ | $T_{CC}$ vs $T_{S\text{-}G2\text{-}M}$ | $T_{G1}$ vs $T_{S\text{-}G2\text{-}M}$ | n |
|---|---|---|---|---|---|---|---|---|---|---|---|---|---|---|
| | | Mean (h) | SD (h) | CV (%) | Mean (h) | SD (h) | CV (%) | Mean (h) | SD (h) | CV (%) | $R^2$ | $R^2$ | $R^2$ | |
| **M-1** | 2 | 28.3 | 6.8 | 23.9 | 12.2 | 4.1 | 33.4 | 16.1 | 4.8 | 29.6 | 0.516 | 0.649 | 0.028 | 300 |
| | 10 | 24.9 | 6.5 | 26.3 | 9.0 | 3.5 | 38.6 | 15.9 | 5.0 | 31.6 | 0.426 | 0.724 | 0.025 | 300 |
| **M-2** | 2 | 28.1 | 2.1 | 7.4 | 13.9 | 1.6 | 11.5 | 14.1 | 0.8 | 5.6 | 0.885 | 0.525 | 0.201 | 300 |
| | 10 | 23.3 | 0.7 | 3.1 | 8.5 | 0.7 | 8.1 | 14.8 | 0.5 | 3.0 | 0.641 | 0.167 | 0.048 | 300 |
| **M-3** | 2 | 26.9 | 3.2 | 11.9 | 12.8 | 2.2 | 17.5 | 14.1 | 1.6 | 11.7 | 0.767 | 0.573 | 0.121 | 300 |
| | 10 | 22.7 | 2.7 | 12.1 | 8.4 | 1.2 | 14.9 | 14.3 | 1.8 | 12.6 | 0.724 | 0.867 | 0.361 | 300 |
| **M-4** | 2 | 25.6 | 4.2 | 16.3 | 11.8 | 3.1 | 26.3 | 13.8 | 2.1 | 15.3 | 0.761 | 0.49 | 0.069 | 300 |
| | 10 | 23.2 | 4.2 | 17.9 | 8.3 | 2.2 | 26.1 | 14.9 | 3.4 | 22.4 | 0.355 | 0.731 | 0.009 | 300 |
| **M-5** | 2 | 27.5 | 4.3 | 15.6 | 13.1 | 2.7 | 20.8 | 14.4 | 2.3 | 16.1 | 0.766 | 0.68 | 0.201 | 300 |
| | 10 | 24.2 | 2.6 | 10.7 | 8.7 | 1.2 | 14.3 | 15.4 | 1.9 | 12.4 | 0.506 | 0.789 | 0.096 | 300 |
| **M-6** | 2 | 29.3 | 7.4 | 25.3 | 13.3 | 4.9 | 37.1 | 16 | 4.6 | 29 | 0.624 | 0.574 | 0.039 | 300 |
| | 10 | 23.5 | 5 | 21.3 | 8.7 | 3.1 | 35.2 | 14.8 | 3.8 | 25.7 | 0.425 | 0.629 | 0.003 | 300 |

| Model Type | Serum Level (%) | Sister pairs (Pearson, R) | | | | Cousin pairs (Pearson, R) | | | | Mother-daughter pairs (Pearson, R) | | | |
|---|---|---|---|---|---|---|---|---|---|---|---|---|---|
| | | $T_{CC}$ | $T_{G1}$ | $T_{S\text{-}G2\text{-}M}$ | n | $T_{CC}$ | $T_{G1}$ | $T_{S\text{-}G2\text{-}M}$ | n | $T_{CC}$ | $T_{G1}$ | $T_{S\text{-}G2\text{-}M}$ | n |
| **M-1** | 2 | 0.685 | 0.91 | 0.46 | 132 | 0.294 | 0.349 | 0.29 | 77 | 0.264 | 0.392 | 0.557 | 141 |
| | 10 | 0.604 | 0.571 | 0.602 | 128 | 0.453 | 0.488 | 0.383 | 131 | 0.505 | 0.591 | 0.529 | 182 |
| **M-2** | 2 | 0.722 | 0.652 | 0.396 | 150 | 0.052 | -0.04 | -0.027 | 200 | 0.014 | 0.075 | 0.066 | 200 |
| | 10 | 0.435 | 0.789 | 0.149 | 150 | -0.082 | -0.076 | -0.004 | 200 | -0.068 | -0.065 | -0.026 | 200 |
| **M-3** | 2 | 1 | 1 | 1 | 150 | 1 | 1 | 1 | 188 | 0.96 | 0.972 | 0.962 | 188 |
| | 10 | 1 | 1 | 1 | 150 | 1 | 1 | 1 | 224 | 0.991 | 0.95 | 0.98 | 224 |
| **M-4** | 2 | 0.925 | 1 | 0.728 | 136 | 0.422 | 0.282 | 0.429 | 112 | 0.617 | 0.532 | 0.73 | 172 |
| | 10 | 0.748 | 1 | 0.631 | 137 | 0.502 | 0.454 | 0.359 | 166 | 0.623 | 0.639 | 0.68 | 206 |
| **M-5** | 2 | 0.88 | 0.828 | 0.848 | 137 | 0.795 | 0.762 | 0.801 | 159 | 0.805 | 0.791 | 0.813 | 175 |
| | 10 | 0.935 | 0.895 | 0.933 | 148 | 0.875 | 0.893 | 0.828 | 204 | 0.887 | 0.848 | 0.846 | 210 |
| **M-6** | 2 | 0.698 | 0.575 | 0.65 | 131 | 0.219 | 0.282 | 0.12 | 73 | 0.391 | 0.356 | 0.475 | 129 |
| | 10 | 0.617 | 0.629 | 0.494 | 136 | 0.582 | 0.364 | 0.53 | 117 | 0.556 | 0.573 | 0.659 | 195 |

**Table S10.** Changes in the model (**Table S3**) to incorporate the effect of the p38-inihibitor in a phenomenological manner.

| | |
|---|---|
| $\frac{dMyc_m}{dt} = (b_{myc} \times s) + k_{Mcm1} \times MV \times (1 + (k_{p1} \times p38i)) + \frac{k_{Mcm2} \times s \times DE^n}{(k_{mm}^n \times s^n) + DE^n} - k_{dMcm} \times Myc_m$ | 46 |
| $\frac{dCycD_m}{dt} = k_{CDm1} \times MV \times (1 + (k_{p1} \times p38i)) + k_{CDm2} \times Myc_p - k_{dCDm} \times CycD_m$ | 47 |
| $\frac{dI_m}{dt} = \frac{k_{sim} \times s}{(1 + (k_{p1} \times p38i))} - k_{dim} \times I_m$ | 48 |
| $\frac{dC25_T}{dt} = \frac{k_{sc25} \times C25_m}{(1 + (k_{p2} \times p38i))} - k_{dc25} \times C25_T$ | 49 |

**Table S11.** Table showing the data from stochastic simulation (Model, M-1 with unequal division) at 2% serum in the presence and absence of p38 inhibitor.

| 2% serum | | $T_{CC}$ | | | $T_{G1}$ | | | $T_{S\text{-}G2\text{-}M}$ | | | $T_{CC}$ vs $T_{G1}$ | $T_{CC}$ vs $T_{S\text{-}G2\text{-}M}$ | $T_{G1}$ vs $T_{S\text{-}G2\text{-}M}$ | n |
|---|---|---|---|---|---|---|---|---|---|---|---|---|---|---|
| | | Mean (h) | SD (h) | CV (%) | Mean (h) | SD (h) | CV (%) | Mean (h) | SD (h) | CV (%) | $R^2$ | $R^2$ | $R^2$ | |
| Control | I | 29.3 | 7.4 | 25.3 | 13.3 | 4.9 | 37.1 | 16.0 | 4.6 | 29 | 0.624 | 0.574 | 0.039 | 300 |
| | II | 27.7 | 7 | 25.4 | 12.7 | 4.8 | 37.5 | 15.1 | 4.0 | 26.7 | 0.7 | 0.582 | 0.081 | 300 |
| | III | 27.0 | 7.3 | 26.9 | 11.8 | 4.4 | 36.9 | 15.3 | 4.4 | 28.8 | 0.689 | 0.694 | 0.146 | 300 |
| | avg | 28.0 | 7.2 | 25.9 | 12.6 | 4.7 | 37.2 | 15.5 | 4.3 | 28.2 | 0.671 | 0.617 | 0.089 | |
| 2% serum | | $T_{CC}$ | | | $T_{G1}$ | | | $T_{S\text{-}G2\text{-}M}$ | | | $T_{CC}$ vs $T_{G1}$ | $T_{CC}$ vs $T_{S\text{-}G2\text{-}M}$ | $T_{G1}$ vs $T_{S\text{-}G2\text{-}M}$ | n |
| | | Mean (h) | SD (h) | CV (%) | Mean (h) | SD (h) | CV (%) | Mean (h) | SD (h) | CV (%) | $R^2$ | $R^2$ | $R^2$ | |
| p38i | I | 26.7 | 8.2 | 30.7 | 9.8 | 2.9 | 29.2 | 16.8 | 7.3 | 43.6 | 0.212 | 0.879 | 0.015 | 300 |
| | II | 24.2 | 5.2 | 21.6 | 9.4 | 2.7 | 28.7 | 14.9 | 4.1 | 27.4 | 0.416 | 0.746 | 0.029 | 300 |
| | III | 25.5 | 6.7 | 26.2 | 9.3 | 2.7 | 29.1 | 16.2 | 5.6 | 34.4 | 0.329 | 0.844 | 0.041 | 300 |
| | avg | 25.5 | 6.7 | 26.2 | 9.5 | 2.8 | 29.0 | 16.0 | 5.7 | 35.1 | 0.319 | 0.823 | 0.028 | |

| 2% serum | | Sister pairs (Pearson, R) | | | | Cousin pairs (Pearson, R) | | | | Mother-daughter pairs (Pearson, R) | | | |
|---|---|---|---|---|---|---|---|---|---|---|---|---|---|
| | | $T_{CC}$ | $T_{G1}$ | $T_{S\text{-}G2\text{-}M}$ | n | $T_{CC}$ | $T_{G1}$ | $T_{S\text{-}G2\text{-}M}$ | n | $T_{CC}$ | $T_{G1}$ | $T_{S\text{-}G2\text{-}M}$ | n |
| Control | I | 0.698 | 0.575 | 0.65 | 131 | 0.219 | 0.282 | 0.120 | 73 | 0.391 | 0.356 | 0.475 | 129 |
| | II | 0.832 | 0.854 | 0.614 | 136 | 0.394 | 0.363 | 0.352 | 83 | 0.322 | 0.343 | 0.391 | 146 |
| | III | 0.835 | 0.822 | 0.75 | 130 | 0.146 | 0.073 | 0.266 | 81 | 0.419 | 0.382 | 0.494 | 147 |
| | avg | 0.788 | 0.75 | 0.671 | | 0.253 | 0.239 | 0.246 | | 0.377 | 0.36 | 0.453 | |
| 2% serum | | Sister pairs (Pearson, R) | | | | Cousin pairs (Pearson, R) | | | | Mother-daughter pairs (Pearson, R) | | | |
| | | $T_{CC}$ | $T_{G1}$ | $T_{S\text{-}G2\text{-}M}$ | n | $T_{CC}$ | $T_{G1}$ | $T_{S\text{-}G2\text{-}M}$ | n | $T_{CC}$ | $T_{G1}$ | $T_{S\text{-}G2\text{-}M}$ | n |
| p38i | I | 0.474 | 0.904 | 0.329 | 133 | 0.305 | 0.57 | 0.032 | 80 | 0.489 | 0.52 | 0.572 | 154 |
| | II | 0.581 | 0.636 | 0.461 | 135 | 0.345 | 0.278 | 0.428 | 113 | 0.469 | 0.384 | 0.637 | 177 |
| | III | 0.578 | 0.902 | 0.378 | 131 | 0.375 | 0.34 | 0.274 | 95 | 0.544 | 0.501 | 0.551 | 167 |
| | avg | 0.544 | 0.814 | 0.389 | | 0.342 | 0.396 | 0.245 | | 0.501 | 0.468 | 0.587 | |

**Table S12.** Table showing the data from HeLa cells grown at 2% serum in DMSO control and 10 μM p38 inhibitor

| 2% serum | | $T_{CC}$ | | | $T_{G1}$ | | | $T_{S\text{-}G2\text{-}M}$ | | | $T_{CC}$ vs $T_{G1}$ | $T_{CC}$ vs $T_{S\text{-}G2\text{-}M}$ | $T_{G1}$ vs $T_{S\text{-}G2\text{-}M}$ | n |
|---|---|---|---|---|---|---|---|---|---|---|---|---|---|---|
| | | Mean (h) | SD (h) | CV (%) | Mean (h) | SD (h) | CV (%) | Mean (h) | SD (h) | CV (%) | $R^2$ | $R^2$ | $R^2$ | |
| DMSO control | I | 24.6 | 4.5 | 18.3 | 11.1 | 3.8 | 34.2 | 13.5 | 2.5 | 18.5 | 0.696 | 0.279 | 0.001 | 297 |
| | II | 23.9 | 4.8 | 20.1 | 9.6 | 3.3 | 34.4 | 14.2 | 3.3 | 23.2 | 0.543 | 0.540 | 0.007 | 308 |
| | III | 24.5 | 5.2 | 21.2 | 11.7 | 4.2 | 36.0 | 12.8 | 2.7 | 21.2 | 0.732 | 0.347 | 0.007 | 309 |
| | avg | 24.3 | 4.8 | 19.9 | 10.8 | 3.8 | 34.9 | 13.5 | 2.8 | 21.0 | 0.657 | 0.389 | 0.005 | |

| 2% serum | | $T_{CC}$ | | | $T_{G1}$ | | | $T_{S\text{-}G2\text{-}M}$ | | | $T_{CC}$ vs $T_{G1}$ | $T_{CC}$ vs $T_{S\text{-}G2\text{-}M}$ | $T_{G1}$ vs $T_{S\text{-}G2\text{-}M}$ | n |
|---|---|---|---|---|---|---|---|---|---|---|---|---|---|---|
| | | Mean (h) | SD (h) | CV (%) | Mean (h) | SD (h) | CV (%) | Mean (h) | SD (h) | CV (%) | | | | |
| 10 μM p38i | I | 26.0 | 4.5 | 17.3 | 10.6 | 2.8 | 26.4 | 15.4 | 3.1 | 20.1 | 0.515 | 0.619 | 0.018 | 301 |
| | II | 26.3 | 4.4 | 16.7 | 9.6 | 2.2 | 22.9 | 16.8 | 3.7 | 22.0 | 0.305 | 0.744 | 0.003 | 246 |
| | III | 23.0 | 4.0 | 17.4 | 9.2 | 3.0 | 32.3 | 13.8 | 2.4 | 17.2 | 0.652 | 0.464 | 0.014 | 307 |
| | avg | 25.1 | 4.3 | 17.1 | 9.8 | 2.7 | 27.2 | 15.3 | 3.1 | 19.8 | 0.491 | 0.609 | 0.012 | |

| 2% serum | | Sister pairs (Pearson, R) | | | | Cousin pairs (Pearson, R) | | | | Mother-daughter pairs (Pearson, R) | | | |
|---|---|---|---|---|---|---|---|---|---|---|---|---|---|
| | | $T_{CC}$ | $T_{G1}$ | $T_{S\text{-}G2\text{-}M}$ | n | $T_{CC}$ | $T_{G1}$ | $T_{S\text{-}G2\text{-}M}$ | n | $T_{CC}$ | $T_{G1}$ | $T_{S\text{-}G2\text{-}M}$ | n |
| DMSO control | I | 0.7504 | 0.8341 | 0.5961 | 136 | 0.3758 | 0.4861 | 0.3309 | 151 | 0.3758 | 0.4861 | 0.3309 | 181 |
| | II | 0.7766 | 0.8683 | 0.7343 | 141 | 0.2249 | 0.4219 | 0.2347 | 129 | 0.2785 | 0.3649 | 0.3214 | 186 |
| | III | 0.7077 | 0.7683 | 0.4324 | 143 | 0.2955 | 0.3344 | 0.4320 | 124 | 0.2547 | 0.1920 | 0.5520 | 159 |
| | avg | 0.7449 | 0.8236 | 0.5876 | | 0.2987 | 0.4141 | 0.3325 | | 0.303 | 0.3477 | 0.4014 | |

| 2% serum | | Sister pairs (Pearson, R) | | | | Cousin pairs (Pearson, R) | | | | Mother-daughter pairs (Pearson, R) | | | |
|---|---|---|---|---|---|---|---|---|---|---|---|---|---|
| | | $T_{CC}$ | $T_{G1}$ | $T_{S\text{-}G2\text{-}M}$ | n | $T_{CC}$ | $T_{G1}$ | $T_{S\text{-}G2\text{-}M}$ | n | $T_{CC}$ | $T_{G1}$ | $T_{S\text{-}G2\text{-}M}$ | n |
| 10 μM p38i | I | 0.6636 | 0.6716 | 0.6026 | 137 | 0.3741 | 0.1224 | 0.5591 | 103 | 0.3397 | 0.3266 | 0.4681 | 130 |
| | II | 0.6278 | 0.7772 | 0.614 | 112 | 0.5494 | 0.3432 | 0.6278 | 95 | 0.3258 | 0.2402 | 0.323 | 126 |
| | III | 0.7891 | 0.8161 | 0.6525 | 106 | 0.4542 | 0.4882 | 0.3126 | 153 | 0.2972 | 0.2886 | 0.1365 | 179 |
| | avg | 0.6935 | 0.7550 | 0.623 | | 0.4592 | 0.3179 | 0.4998 | | 0.3209 | 0.2851 | 0.3092 | |

**Movie S1.** Time-Lapse Imaging of FUCCI-Expressing HeLa cells, related to Fig. 1. Asynchronously growing HeLa expressing FUCCI were imaged for 72 h at 2% serum. Images were collected every 15 minutes in 15 different positions in the culture dish. This is one of the representative positions.

**Movie S2.** Time-Lapse Imaging of FUCCI-Expressing HeLa cells, related to Fig. 1. Asynchronously growing HeLa expressing FUCCI were imaged for 72 h at 2% serum from another representative position other than shown in Movie S1.

**Movie S3.** Time-Lapse Imaging of FUCCI-Expressing HeLa cells, related to Fig. 1. Asynchronously growing HeLa expressing FUCCI were imaged for 72 h at 10% serum. Images were collected every 15 minutes in 15 different positions in the culture dish. This is one of the representative positions.

**Movie S4.** Time-Lapse Imaging of FUCCI-Expressing HeLa cells, related to Fig. 1. Asynchronously growing HeLa expressing FUCCI were imaged for 72 h at 10% serum from another representative position other than shown in Movie S3.
